# Synthesis and characterization of 5-(2-fluoro-4-[^11^C]methoxyphenyl)-2,2-dimethyl-3,4-dihydro-2*H*-pyrano[2,3-*b*]pyridine-7-carboxamide as a PET imaging ligand for metabotropic glutamate receptor 2

**DOI:** 10.1101/2021.06.29.450413

**Authors:** Gengyang Yuan, Maeva Dhaynaut, Yu Lan, Nicolas J. Guehl, Dalena Huynh, Suhasini Iyengar, Sepideh Afshar, Hao Wang, Sung-Hyun Moon, Mary Jo Ondrechen, Changning Wang, Timothy Shoup, Georges El Fakhri, Marc D. Normandin, Anna-Liisa Brownell

## Abstract

Metabotropic glutamate receptor 2 (mGluR2) is a therapeutic target for the treatment of several neuropsychiatric disorders and conditions. The role of mGluR2 function in etiology could be unveiled by *in vivo* imaging using positron emission tomography (PET). In this regard, 5-(2- fluoro-4-[^11^C]methoxyphenyl)-2,2-dimethyl-3,4-dihydro-2*H*-pyrano[2,3-*b*]pyridine-7- carboxamide ([^11^C]**13**), a potent negative allosteric modulator (NAM), was developed to support this endeavor. Radioligand [^11^C]**13** was synthesized via the *O*-[^11^C]methylation of phenol **24** with a high molar activity of 212 ± 76 GBq/µmol (n = 5) and excellent radiochemical purity (> 99%). PET imaging of [^11^C]**13** in rats demonstrated its superior brain heterogeneity, particularly in the regions of striatum, thalamus, hippocampus, and cortex. Accumulation of [^11^C]**13** in these regions of interest (ROIs) was reduced with pretreatment of mGluR2 NAMs, VU6001966 (**9**) and MNI-137 (**26**), the extent of which revealed a time-dependent drug effect of the blocking agents. In a nonhuman primate, [^11^C]**13** selectively accumulated in mGluR2-rich regions, especially in different cortical areas, putamen, thalamus, and hippocampus, and resulted in high-contrast brain images. The regional total volume of distribution (*V_T_*) estimates of [^11^C]**13** decreased by 14% after the pretreatment with **9**. Therefore, [^11^C]**13** is a potential candidate for translational PET imaging studies of mGluR2 function.

## INTRODUCTION

As the most abundant endogenous neurotransmitter in the central nervous system (CNS), glutamate has an important role in regulating several neurological functions in the brain.^1–2^ There are two families of glutamate receptors, namely the ionotropic glutamate receptors (iGluRs) and the metabotropic glutamate receptors (mGluRs).^3^ The mGluRs are further divided into three groups based on their sequence homology, pharmacological effects, and distribution.^4^ Among them, the group II mGluRs, including mGluR2 and mGluR3, are promising targets for drug discovery, especially for the treatment of schizophrenia,^5–6^ anxiety,^7–8^ depression,^9^ pain,^10^ and Alzheimer’s disease^11^. The rationale is that mGluR2 and mGluR3 are highly distributed in the forebrain at the presynaptic nerve terminals and activation of these receptors reduces the excessive glutamatergic signaling that is implicated in the pathophysiology of these diseases.^9, 12^ Despite the setback of LY2140023,^6^ a group II agonist prodrug, in clinical trials for the treatment of schizophrenia,^13–14^ it demonstrated the disease-modifying potential of targeting the mGluR2- focused glutamatergic signaling and emphasized the importance of mGluR2-subtype selectivity for successful drug candidates.^15–16^ As a result, allosteric modulators that bind to the more lipophilic and structurally less conserved seven transmembrane (7-TM) region are developed to afford ligands with more favorable physiochemical properties and enhanced selectivity for mGluR2 binding.^17–18^ Similarly, development of positron emission tomography (PET) radioligands targeting mGluR2 have shifted from the early group II orthosteric ligands, such as the mGluR2/3 antagonists [^11^C]MMMHC (**1**)^19^ and [^11^C]CMGDE (**2**)^20^, to the recent allosteric modulators-derived radiotracers, such as the positive allosteric modulators (PAMs) of [^11^C]JNJ- 42491293 (**3**)^21^, [^11^C]mG2P001 (**4**),^22–23^ [^18^F]JNJ-46356479 (**5**)^24^ and [^18^F]mG2P026 (**6**)^25^ (Fig. 1).

As a non-invasive *in vivo* imaging technique, PET will enable the visualization and quantification of mGluR2 under normal and disease conditions as well as the evaluation of target engagement and the dose occupancy studies of drug candidates. Currently, there is no suitable mGluR2 PET tracer for humans. [^11^C]JNJ-42491293 (**3**), the only structurally disclosed PET tracer that entered clinical trials, showed unexpected binding in the myocardium and off-target binding in the brain.^21^ Besides our current efforts of developing a new series of mGluR2 PAM radiotracers for clinical use,^22–25^ we are also devoted to identifying negative allosteric modulators (NAMs)- based radiotracers due to their distinct allosteric mode of action and pharmacology.^26–29^ As noted by O’Brien *et al*, mGluR2 PAMs had both affinity and efficacy cooperativity with glutamate, whereas mGluR2 NAMs showed predominantly efficacy cooperativity.^26^ The development of NAM-based PET tracers is still nascent without any viable tracers reported in higher species. At the beginning of our work, only [^11^C]QCA (**7**, IC_50_ = 45 nM)^30^ and [^11^C]MMP (**8**, IC_50_ = 59 nM)^31^ were disclosed. However, these tracers suffered poor brain permeability in rats with a SUV_max_ value of 0.3 and 0.7, respectively. Further studies of these radiotracers in the P-glycoprotein and the breast cancer resistance protein (Pgp-BCRP) knock-out mouse model indicated that they are likely substrates of the efflux pumps on the blood-brain barrier (BBB).^30–31^ QCA (**7**) was presented in a patent application filed by Merck Research Laboratories in 2013^32^ and the structure-activity relationship was further explored by Felts *et al* in 2015.^33^ Compound MMP (**8**) is an analogous NAM of VU6001966 (**9**, IC_50_ = 78 nM)^34^, developed by Bollinger *et al*, has higher brain permeability than QCA (**7**), thus it was deemed as a promising PET imaging candidate.^34^ However, according to the recent publication, the ^11^C-labeled VU6001966 (**9**) has the same issues as [^11^C]QCA (**7**).^35^ Since there are no explanations on the structural basis of the poor brain permeability for these NAM tracers, we searched for a completely new chemical scaffold to avoid this issue. The recent surge on the development of mGluR2 NAMs has resulted in a number of patent applications and research publications, providing an ample reservoir of PET imaging candidates with distinct structures. For instance, the recently published NAM tracers of [^11^C]MG- 1904 (**10**, IC_50_ = 24 nM),^36^ which was selected from a series of tetrahydronaphthyridine derivatives patented by Merck in 2016,^37^ and [^11^C]MG2-1812 (**11**, IC_50_ = 21 nM),^35^ which was a close analog of VU6001966 (**9**),^34^ were brain permeable in rats.

After a comprehensive examination of the chemical scaffolds, we selected the 3,4-dihydro-2*H*- pyrano[2,3-*b*]pyridine derivative, 5-(2,4-difluorophenyl)-2,2-dimethyl-3,4-dihydro-2*H*-pyrano [2,3-*b*]pyridine-7-carboxamide (**12**), as a lead compound. Compound **12** was reported as a potent mGluR2 NAM (IC_50_ = 6.0 nM) by Merck in 2018.^38^ Although only limited information was provided for compound **12**, we envisioned this compound to be an ideal starting point due to its potent modulatory affinity, absence of a chiral center, and ease to introduce structural variances for future structure-activity analysis (SAR). As a proof-of-concept study, compound **12** and/or its analogs are expected to be radiolabeled with convenient methods to allow a rapid PET imaging evaluation of their brain permeability and kinetics. With this in mind, replacement of the *para*- fluoride at compound **12** with a phenolic methyl ether led to the 5-(2-fluoro-4-methoxyphenyl)- 2,2-dimethyl-3,4-dihydro-2*H*-pyrano[2,3-*b*]pyridine-7-carboxamide (**13**). This chemical modification allowed the radiolabeling of **13** with [^11^C]CH_3_I via the *O*-methylation of the corresponding phenol precursor. Herein, the synthesis, *in vitro* characterization and radiolabeling of compounds **12** and **13** as well as the *in vivo* evaluation of [^11^C]**13** in rats and a non-human primate are disclosed.

**Figure 1.**
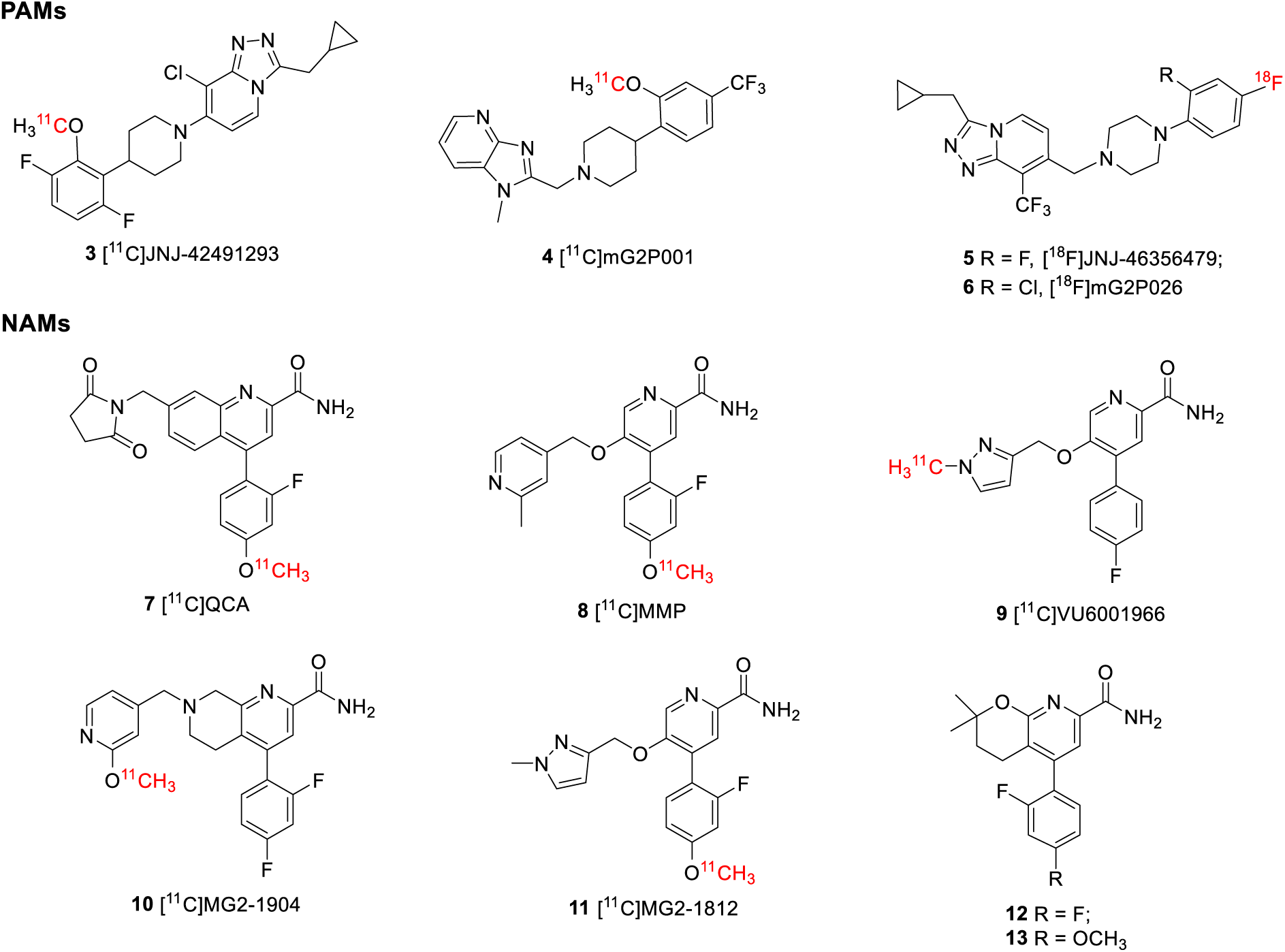
Structures of the mGluR2 allosteric modulators.

## RESULTS AND DISCUSSION

### Chemistry

Syntheses of compounds **12**, **13** and the phenolic precursor **24** are shown in Scheme 1.^38^ The syntheses started from the Wittig reaction between aldehyde **14** and phosphorous ylide **15** to give compound **16**. Hydrogenation of compound **16** under 40 psi of hydrogen at room temperature led to compound **17**, which was used in the subsequent Suzuki coupling reaction with the boronic acid species **18a**-**18c** to furnish compounds **19a**-**19c**. The ester groups in compounds **19a**-**19c** were converted to tertiary alcohol moieties in compounds **20a**-**20c** at 0 °C in the presence of a Grignard reagent. After cyclization of the tertiary alcohols under basic conditions, aryl chlorides **21a**-**21c** were obtained, which were cyanated with Zn(CN)_2_ in a microwave reactor to give aryl nitriles **22a**-**22c**. Finally, hydrolysis of **22a** and **22b** led to compounds **12** and **13**, whereas compound **22c** was deprotected before hydration to afford the radiolabeling precursor **24**.

During the syntheses of these compounds, several modifications were made to the previous methods.^38^ First, the more reactive 4-iodo-2,6-dichloronicotinaldehyde (**14**) instead of 4-bromo- 2,6-dichloronicotinaldehyde was used as starting material. Second, compound **16** was hydrogenated to **17** prior to the Suzuki coupling with **18a**-**18c**. Third, the carboxamide group in compounds **12**, **13** and **24** was introduced by a microwave-assisted cyanation with Zn(CN)_2_ at 160 °C for 30 min followed by hydration with Na_2_CO_3_·1.5H_2_O_2_. Previously, this function group was installed via the palladium-catalyzed esterification of aryl chlorides **21a**-**21c** under 50 psi of carbon monoxide at 80 °C for 30 h and subsequent amidation with ammonia. The new synthetic methods in scheme 1 were robust and gave compounds **12**, **13** and **24** with overall yields of 2.7%, 7.1%, and 1.2%, respectively, whereas the yield of compound **12** was not disclosed previously.^38^

**Scheme 1.**
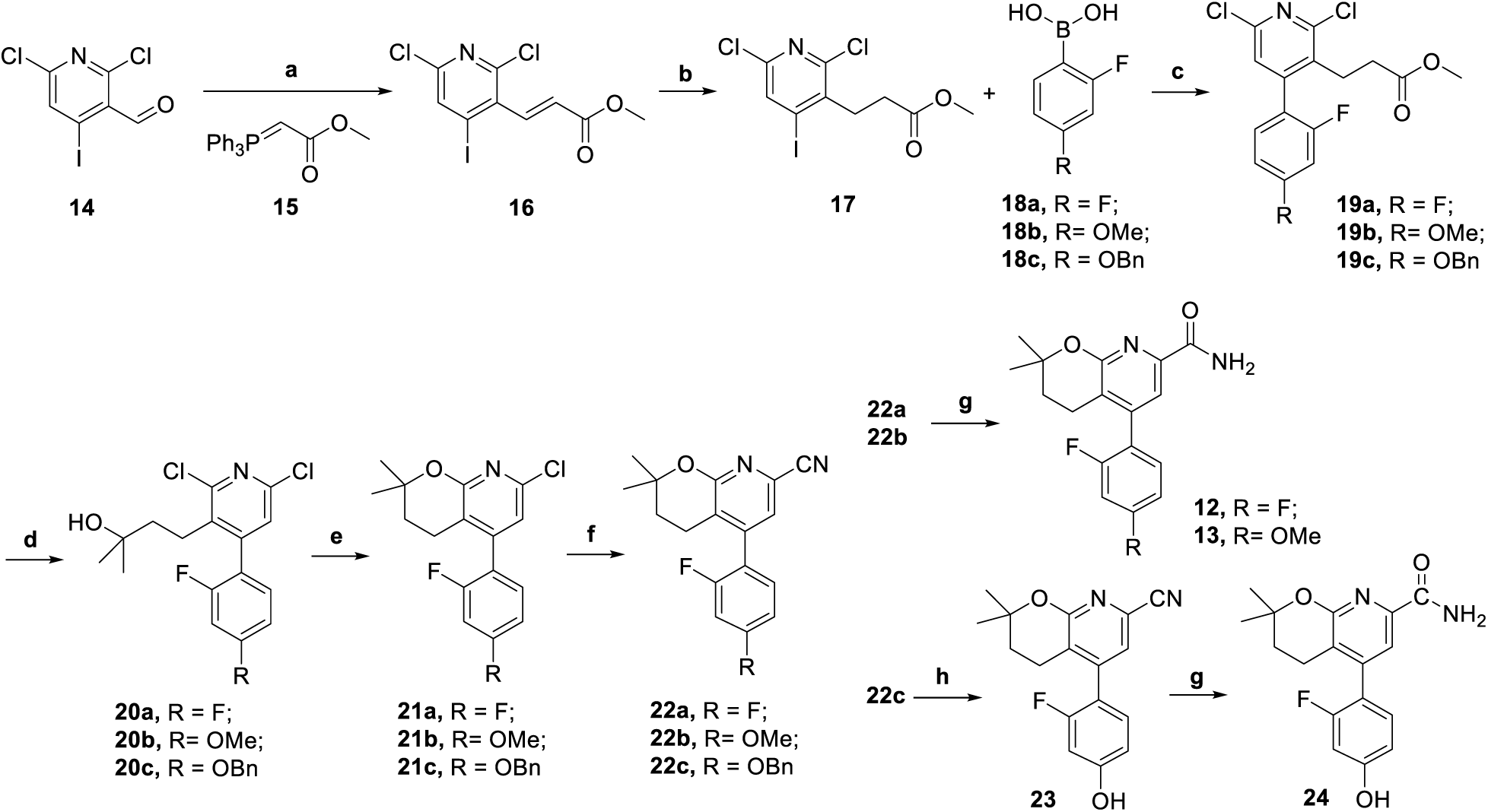
Synthesis of compounds **12**, **13** and **24**. Reagents and conditions: (a) THF, N_2_, 80 °C, 2 h; (b) RhCl(PPh_3_)_3_, H_2_, 40 psi, rt, 2 d; (c) Pd(dppf)Cl_2_, NaHCO_3_, 1,4-dioxane/water, 100 °C, 3 h; (d) MeMgBr (3.0 M in diethyl ether), THF, 0 °C, 1 h; (e) Cs_2_CO_3_, DMA, 120 °C, overnight; (f) Zn(CN)_2_, microwave, 160 °C, 30 min; (g) Na_2_CO_3_·1.5H_2_O_2_, acetone, water, rt, overnight; (h) EtOAc, H_2_, Pd/C (10 wt.%), rt, overnight.

### Pharmacology and Physiochemical Properties

As previously disclosed, compound **12** had a potent mGluR2 negative allosteric modulatory activity (IC_50_ = 6 nM).^38^ The IC_50_ value was determined by measuring the inhibition of glutamate-induced calcium mobilization in Chinese Hamster Ovary (CHO) cells expressing recombinant human mGluR2. Herein, the modulatory activity of compound **13** was tested by monitoring the cAMP modulation using the DiscoverX HitHunter cAMP XS+ assay. The CHO cells expressing recombinant human mGluR2 were used. Compound **13** was determined as a potent mGluR2 NAM with an IC_50_ value of 93.2 nM (Fig. 2A). Besides the mGluR2 binding, the physicochemical properties of compounds **12** and **13** were also characterized using our previously described assays.^22^ The assays assessed their lipophilicity, plasma stability, liver microsome stability, and their effect on recombinant human P-glycoprotein (Pgp).^22^ The lipophilicity of **12** and **13** was initially predicted in ChemDraw 16.0 with a cLogP value of 4.3 and 4.25, respectively (Table 1). This property was further tested using the “shake flask method” to give a LogD_7.4_ value of 2.81 and 2.94 for compounds **12** and **13**, respectively, which are in the preferred range of 1.0-3.5 for brain permeable compounds (Table 1).^39–40^ Compounds **12** showed excellent stabilities in rat plasma and rat liver microsome assays (> 92%), whereas compound **13** had excellent rat plasma stability (94.5%) but moderate rat liver microsome stability (47.8%, Table 1). In addition, compounds **12** and **13** were evaluated by the Pgp-Glo^TM^ assay. The assay detects the effects of a tested compound toward recombinant human Pgp protein in a cell membrane fraction. If the compound is a transport substrate of Pgp, it stimulates the Pgp ATPase reaction, resulting in ATP consumption and subsequent decrease of the luciferase-generated luminescent signal. The basal Pgp ATPase activity was measured by the change in luminescence between sodium orthovanadate (Na_3_VO_4_)-treated controls and untreated samples. Verapamil, a known transport substrate of Pgp, was used as a positive control. As shown in Fig. 2B and Table 1, the change in luminescence for compounds **12** and **13** was similar to that of the basal condition, suggesting neither compound **12** nor compound **13** had any effect with this protein. Therefore, compounds **12** and **13** were employed as candidates for PET imaging ligands.

To probe the ligand-protein binding of compounds **12** and **13**, we have prepared an mGluR2 homology model for NAMs via YASARA^41^ and performed the molecular docking. As shown in Figs. 2C, compounds **12** and **13** adopted similar binding poses in the allosteric binding pocket. For both compounds, the oxygen atom in the carboxamide sidechain forms a hydrogen bond with Asn735 and the nitrogen atom in the carboxamide side chain forms a hydrogen bond with R636. Moreover, compound **13** forms an extra hydrogen bond with its methyl ether oxygen atom to Ser797 and an additional π-π stacking with its phenyl ring toward Phe643. The docking score of compounds **12** and **13** were -11.74 kcal/mol and -11.00 kcal/mol, respectively, indicating their potential nanomolar binding affinity for mGluR2.

**Figure 2.**
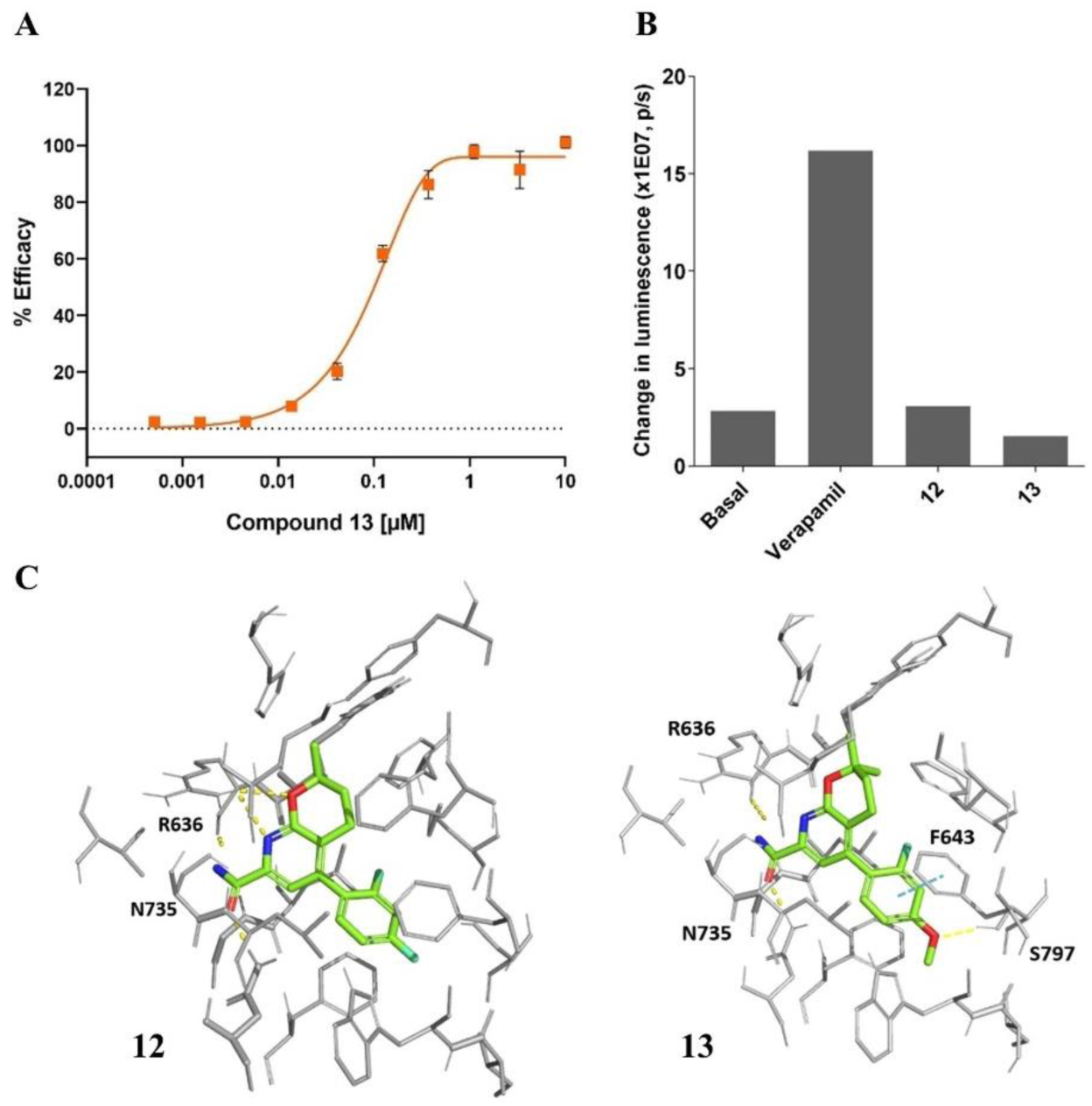
Characterization of compounds **12** and **13**. (**A**) GPCR cAMP modulation result for compound **13**; (**B**) Pgp-Glo™ assay; (**C**) snapshots of the docking poses for compounds **12** and **13**. The key binding residues are shown in gray and the ones interacting with the ligand are labelled. The ligand atoms are rendered as carbon in green, nitrogen in blue, oxygen in red, and fluorine in cyan. Yellow dotted lines represent H-bonds and cyan dotted lines show π-π stacking. Pictures were rendered in PyMol 2.3.3.

**Table 1.** Summary of the *in vitro* and *in silico* results of compounds **12** and **13**

| Comp. | cLogP | LogD<sub>7.4</sub> | IC<sub>50</sub> (nM) | Plasma stability | Microsome stability | Change in luminescence (p/s) | Docking (kcal/mol) |
| --- | --- | --- | --- | --- | --- | --- | --- |
| **12** | 4.3 | 2.81 | 6.0 | 95.3 | 94.5 | 3.07E7 | -11.74 |
| **13** | 4.25 | 2.94 | 93.2 | 92.2 | 47.8 | 1.53E7 | -11.00 |

### Radiochemistry

Although compound **12** had better pharmacological and physicochemical properties than compound **13**, radiolabeling of this compound was challenging. As shown in scheme 2, the first attempted method utilized the palladium catalyzed cyanation of compound **21a** with [^11^C]HCN^42–43^ and subsequent amidation of [^11^C]**22a** with hydrogen peroxide to get [^11^C]**12**. Unfortunately, although the unlabeled compound **22a** could be prepared from compound **21a** with a 71% yield at 160 °C for 30 min under the microwave conditions, [^11^C]**22a** was not obtained under the conventional heating at 160 °C for 5- or 10-min. Precursor **21a** was intact at 180 °C for 20 min, indicating its insufficient reactivity for such radiosynthesis. In addition, the significantly changed stoichiometry between the precursor **21a** and the cyanide source might contribute to this failure considering [^11^C]CN^-^ was in the nano- or pico-molar scale. We then tried the Ru-mediated deoxyfluorination of compound **23** using [^18^F]fluoride.^44–45^ However, under the typical radiofluorination conditions, **23** readily decomposed without forming [^18^F]**22a**. Alternatively, replacement of the aryl chloride in compound **21a** with aryl bromide or iodide could allow the incorporation of [^11^C]CN group due to the enhanced reactivity. Moreover, the Cu-mediated radiofluorination of the corresponding boronic acid/ester or alkyl tin precursor **25** may give the desired [^18^F]**22a**.^46–47^ This method has been successfully applied to the automated radiosynthesis of [^18^F]JNJ-46356479 (**5**) in our group.^48^ On the other hand, radiolabeling of compound **13** was less troublesome. [^11^C]**13** was prepared via the one-step *O*-methylation of phenol **24** (1.6 µmol) in anhydrous dimethylformamide (DMF, 0.35 mL) using [^11^C]CH_3_I in the presence of 0.5N NaOH (3.0 µL). The reaction was carried out at 80 °C for 3 min, quenched by addition of 1.0 mL water, and purified by a semipreparative HPLC system. Noteworthy, the HPLC fractions containing [^11^C]**13** could be trapped on a C-18 cartridge and released via 0.6-1.0 mL ethanol with more than 95% recovery rate (n = 5). In the previous radiotracer synthesis, such as [^11^C]QCA (**7**), the product was enriched by removing the HPLC solvents under reduced pressure.^30–31, 35–36^ At the end of synthesis (EOS = 45 min), [^11^C]**13** was obtained with a radiochemical yield of 42 ± 5% (n = 5, non-decay corrected) calculated from starting [^11^C]CO_2_, excellent chemical and radiochemical purities (> 99%), and a high molar activity (A_m_) of 212 ± 76 GBq/µmol (n = 5). As a representative 3,4-dihydro-2*H*-pyrano[2,3-*b*]pyridine NAM tracer, [^11^C]**13** was characterized using *in vivo* PET imaging studies in rats and a non-human primate.

**Scheme 2.**
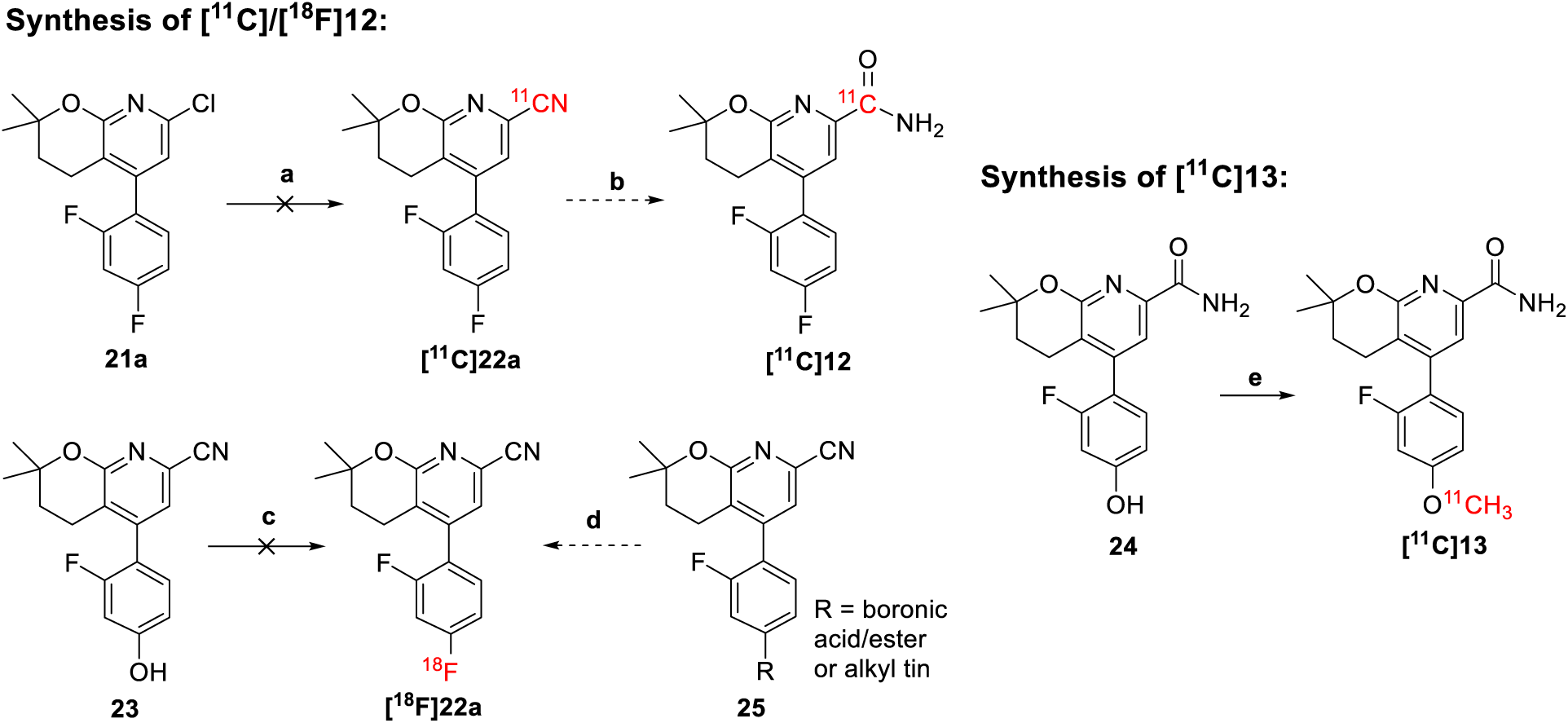
Radiolabeling strategies for compounds **12** and **13**. Reagents and conditions: (a) **21a** (0.32 µmol), Pd(PPh_3_)_4_ (0.9 µmol), [^11^C]HCN (3.5 GBq), DMF (0.1 mL), 160 °C, 5 or 10 min; (b) Na_2_CO_3_·1.5H_2_O_2_ or 35% H_2_O_2_, rt, 2 min; (c) i. **23** (15 µmol), CpRu(COD)Cl (45 µmol), EtOH (50 µL), 85 °C, 30 min; ii. chloroimidazolium chloride (45 µmol), CH_3_CN (150 µL), ^18^F^-^ (1.5 GBq), DMSO (150 µL), 125 °C, 30 min; (d) tetraethylammonium bicarbonate, ^18^F^-^, [Cu(OTf)_2_py_4_], DMF, 130 °C, 10 min; (e) **24** (1.6 µmol), [^11^C]CH_3_I (7.4-74 GBq), 0.5N NaOH (3.0 µL), DMF (0.35 mL), 80 °C, 3 min.

### PET Imaging Studies in Rats

Preliminary PET imaging studies of [^11^C]**13** were carried out in Sprague Dawley rats. Representative TACs and summed PET images at time interval of 1-30 min are shown in Fig. 3. [^11^C]**13** showed excellent brain permeability with a maximum SUV value of 3.6 at 3 min in striatum, which was higher than that of [^11^C]MG-1904 (**10**, SUV_max_ = 1.7)^36^ and [^11^C]MG2-1812 (**11**, SUV_max_ = 1.2)^35^. [^11^C]**13** had a satisfactory tracer kinetics with most of the radioactivity washed out at 30 min (SUV_3min_/SUV_30min_ = 2.7). Accumulation of [^11^C]**13** was high at the mGluR2-rich regions of striatum, thalamus, cortex, hypothalamus, hippocampus, and cerebellum. [^11^C]**13** showed improved brain heterogeneity compared to those of [^11^C]MG-1904 (**10**)^36^ and [^11^C]MG2-1812 (**11**)^35^.

The binding specificity of [^11^C]**13** was examined by pretreatment studies with the selective mGluR2 NAM VU6001966 (**9**)^34^ and the potent group II NAM MNI-137^49^ (**26**, IC_50_ = 8.3 nM). Pretreatment with both compounds were investigated using two different time points, namely, 1 min and 20 min before radioactivity. Pretreatment with **9** (0.5 mg/kg, iv.) 1 min before tracer injection decreased the radioactivity accumulations by 22.4 ± 7.3% across these regions of interest (ROIs) with the cortex having the highest decrease of 38.5% and thalamus the least decrease of 17.1%. However, the blocking effect significantly decreased when this agent was administered 20 min before radioactivity, where the total average decrease was 14.5 ± 1.5%. Administration of **26** (0.2 mg/kg, iv.) 1 min before [^11^C]**13** induced a higher radioactivity decrease among these ROIs by 41.7 ± 1.1% with the hypothalamus having the highest decrease of 42.6% and the cerebellum the least decrease of 39.6%. When compound **26** was administered 20 min before [^11^C]**13**, the blocking effect significantly diminished with an average decrease of 12.7 ± 1.7%. The highest decrease was observed in the cerebellum (15.2%) and the lowest decrease was seen in the striatum (11.1%). Therefore, both compounds **9** and **26** showed a similar blocking pattern where the highest blocking effect occurred when these blocking agents were administered 1 min before radioactivity, whereas this blocking effect diminished with extended time gap of 20 min. It is hypothesized the blocking agents and/or their induced pharmacological effects might wash out over time. Altogether, [^11^C]**13** demonstrated a moderate-to-high level of specific binding toward mGluR2 in rat studies.

**Figure 3.**
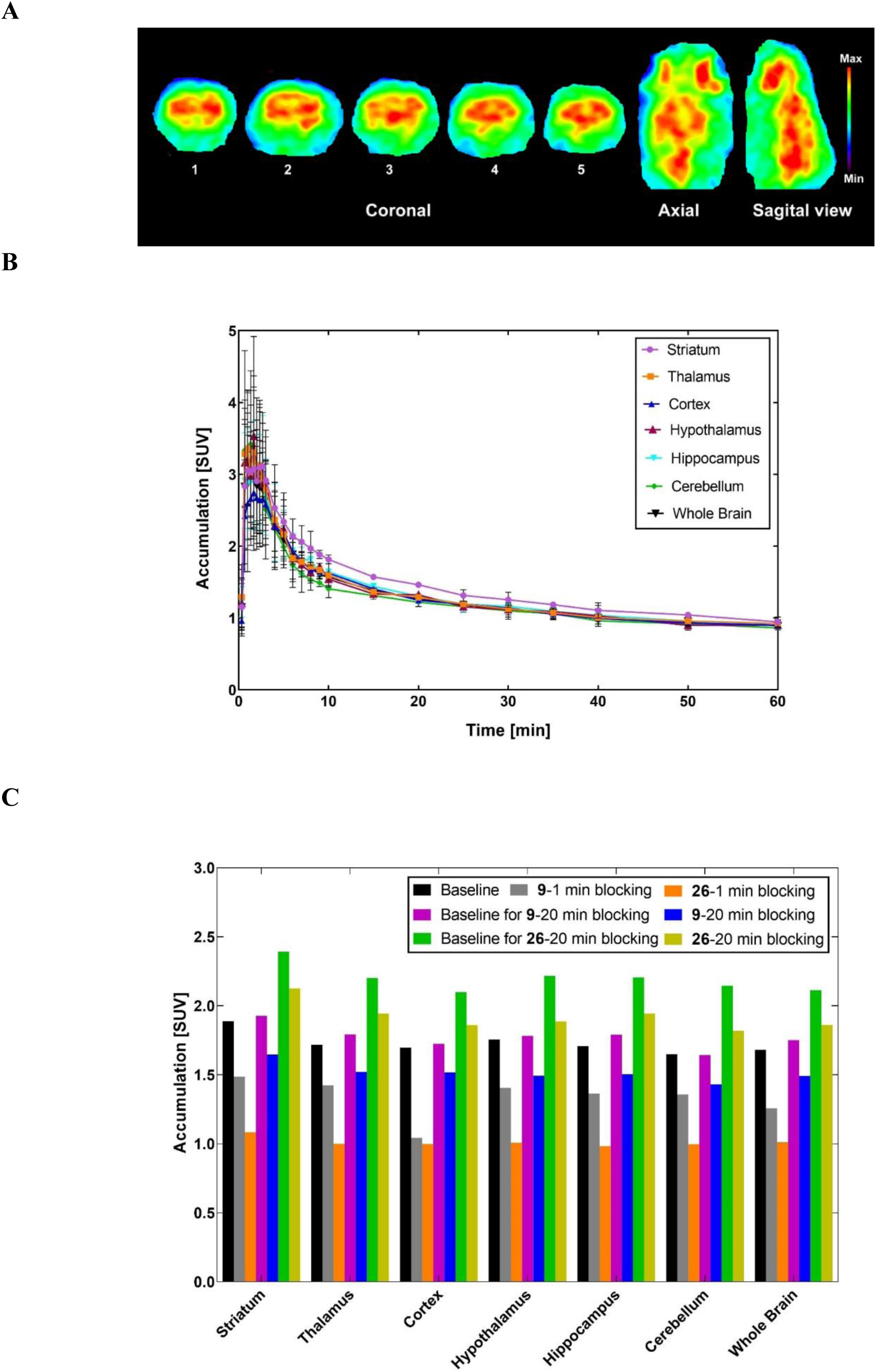
Preliminary PET imaging results of [^11^C]**13** in rat brain. (**A**) Summed PET images at the time interval of 1-30 min. Coronal levels show striatum (1), cingular cortex, striatum, thalamus, hypothalamus (2), cortex, hippocampus, thalamus (3), and cerebellar structures (4 and 5); (**B**) Representative time-activity curves of [^11^C]**13** across the regions of interest; (**C**) Accumulation of radioactivity during the 2-30 min window after pretreatment with VU6001966 (**9**) or MNI-137 (**26**) administered 1 min or 20 min before radioligand. The “Baseline” SUV values were the average of three baseline studies. Pictures were rendered from Prism 9.0.

### PET Imaging Studies in A Non-human Primate

To further characterize [^11^C]**13** as an imaging tool for mGluR2, we performed the PET imaging studies in a cynomolgus monkey. Brain imaging in non-human primate (NHPs) is a pivotal translational approach to study the etiology of human neuropsychiatric diseases, such as schizophrenia^5–6^ and drug addiction^50^. Herein, [^11^C]**13** was characterized for its *in vivo* metabolism in arterial whole-blood (WB) and plasma (PL) as well as for its binding in brain tissues by using kinetic modeling techniques. This effort will facilitate the future application of [^11^C]**13** in humans.

Fig. 4 shows analyses of [^11^C]**13** in arterial blood during the experimental PET imaging studies under the baseline and blocking conditions. The PL/WB ratio was similar in both studies and reached a plateau after 30 min of [^11^C]**13** injection with a mean value of 1.19 ± 0.013. Fig. 4B shows a representative radiometabolite analysis of [^11^C]**13** with selected plasma samples. It revealed the presence of a highly polar metabolite with a retention time (t_R_) of 2.0 min, which was likely the by-product of the [^11^C]CH_3_^-^ cleaved from the phenolic methyl ether of [^11^C]**13**. Besides, there is another polar metabolite near [^11^C]**13** with a t_R_ of 6 min, the structure of which was difficult to identify due to its extremely low amount as a tracer and its absence in neither the *in vitro* plasma nor microsome stability assays. We predicted the top possible sites for the metabolism of **13** via SMARTCyp,^51^ where the phenolic methyl ether was ranked as the first labile group followed by the C3-C4 bond on 3,4-dihydro-2*H*-pyran and the pyridine nitrogen (Supporting Information, Section 5). Measurement of the percent parent (%PP) in plasma revealed a moderate metabolism stability with 53 ± 5.3% of radioactivity attributable to unmetabolized [^11^C]**13** at 30 min and 24.8 ± 1.23% at 120 min (Fig. 4C). The individual metabolite-corrected [^11^C]**13** SUV time courses in plasma is shown in Fig. 4D. The plasma-free fraction (*fp*) of [^11^C]**13** at baseline condition (0.131 ± 0.006) was slightly higher than that in the blocking study (0.099 ± 0.011). The parent fraction curve of [^11^C]**13** fitted well with a Hill function.

**Figure 4.**
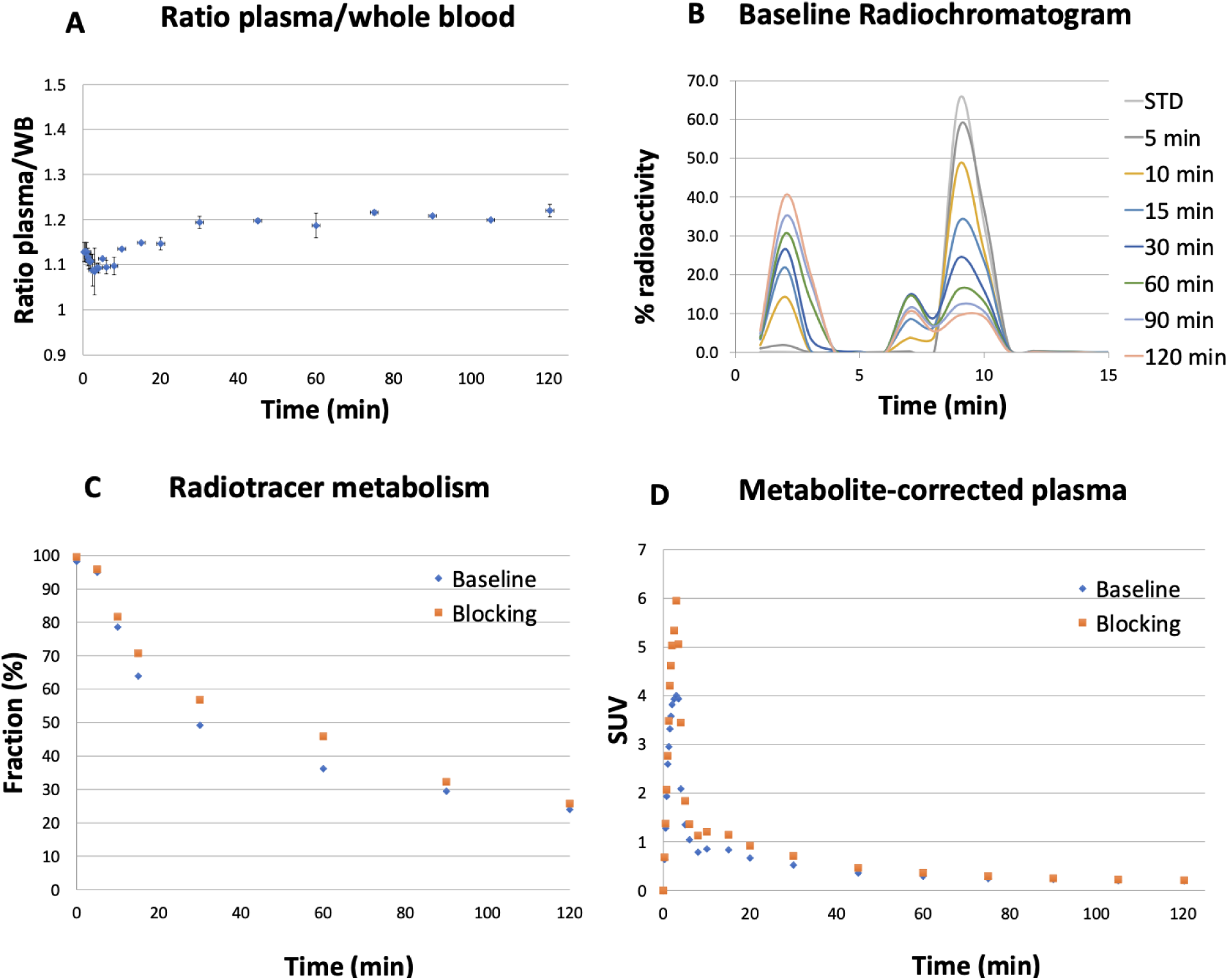
[^11^C]**13** analysis in arterial blood. (**A**) Plasma/Whole blood ratios. (**B**) Representative radiochromatogram of plasma samples from baseline study. (**C**) Individual time course of percent parent in plasma (%PP). (**D**) Individual metabolite-corrected SUV time courses in plasma.

As shown in Fig. 5A, [^11^C]**13** readily crossed the BBB and peaked at 4 min after tracer injection with a SUV value of 7.5 in the striatum in baseline condition. Selected brain regions of striatum, cerebellum non vermis, thalamus, frontal cortex and hippocampus are shown. Pharmacokinetic modeling of [^11^C]**13** was best described by a reversible 2-tissue compartment model (2T4k1v) with a fixed vascular contribution *v* included. According to the Akaike information criteria (AIC),^52^ the 2T4k1v model provided stable regional total volume of distribution (*V_T_*) estimates, which symbolize the equilibrium ratio of [^11^C]**13** in tissue to plasma as shown in Fig. 5A (left). Meanwhile, the Logan plots linearized well with t*30 min and resulted in *V_T_* estimates that were well correlated with those derived from the 2T model despite an underestimation (mean difference equals to 20 ± 6%) as depicted in Fig. 5A (right). The high *K_1_* values (0.7 mL/min/cc) based on the 2T4k1v model indicated high brain penetration. In the pretreatment study, compound **9** was administered 20 min before tracer injection at a dose of 1.0 mg/kg (iv.) considering the species and metabolic rate differences between rodents and NHPs. The *V_T_* estimates decreased in all ROIs over the entire acquisition. Representative Logan *V_T_* estimates obtained when using 120 min and t* of 30 min are shown in Fig. 5B-C, where the decrease of *V_T_* estimates ranges from 16.8% in the cerebellum gray to 3.2% in the occipital gyrus with the average decrease in the whole brain as 14.1%.

**Figure 5.**
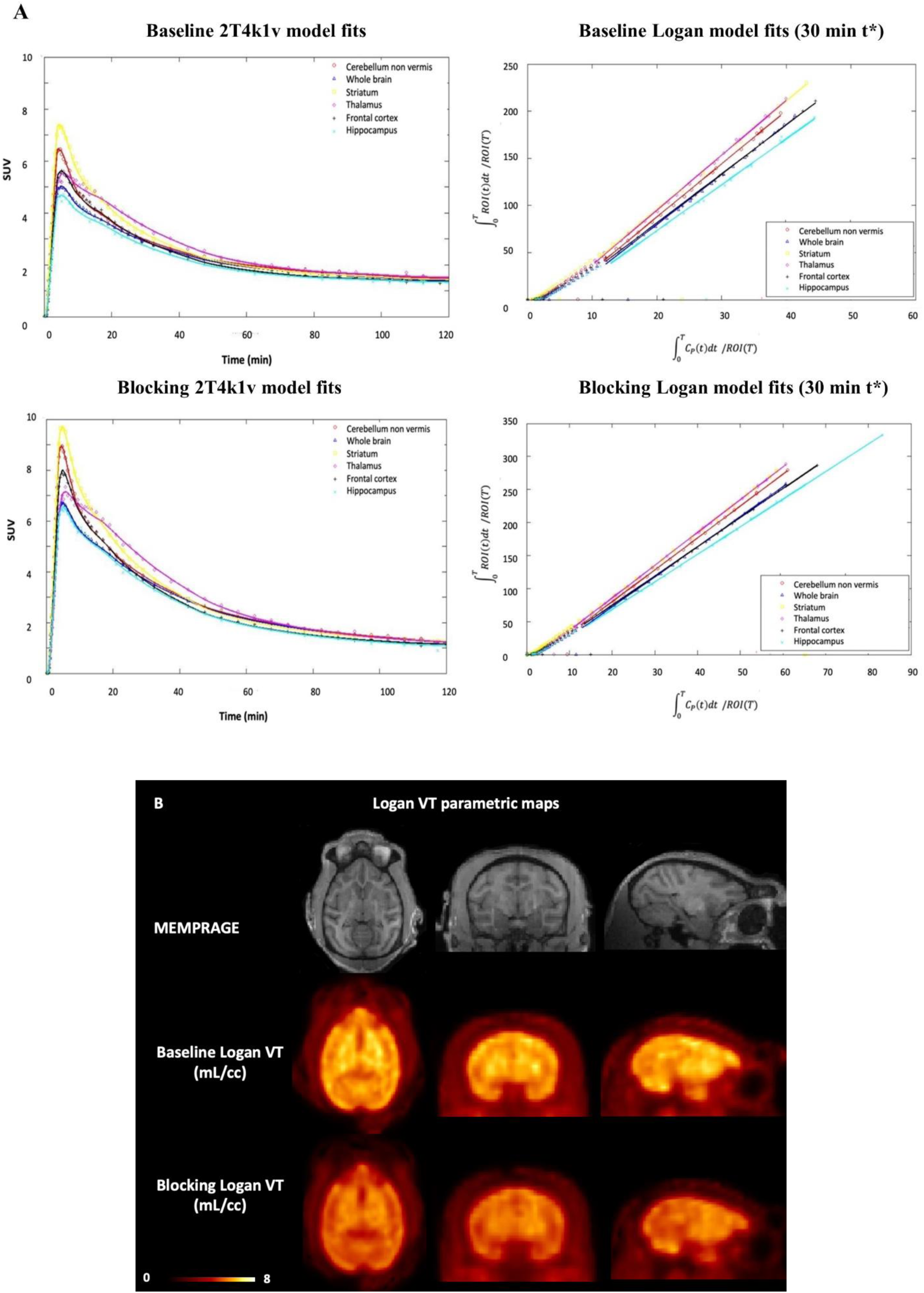
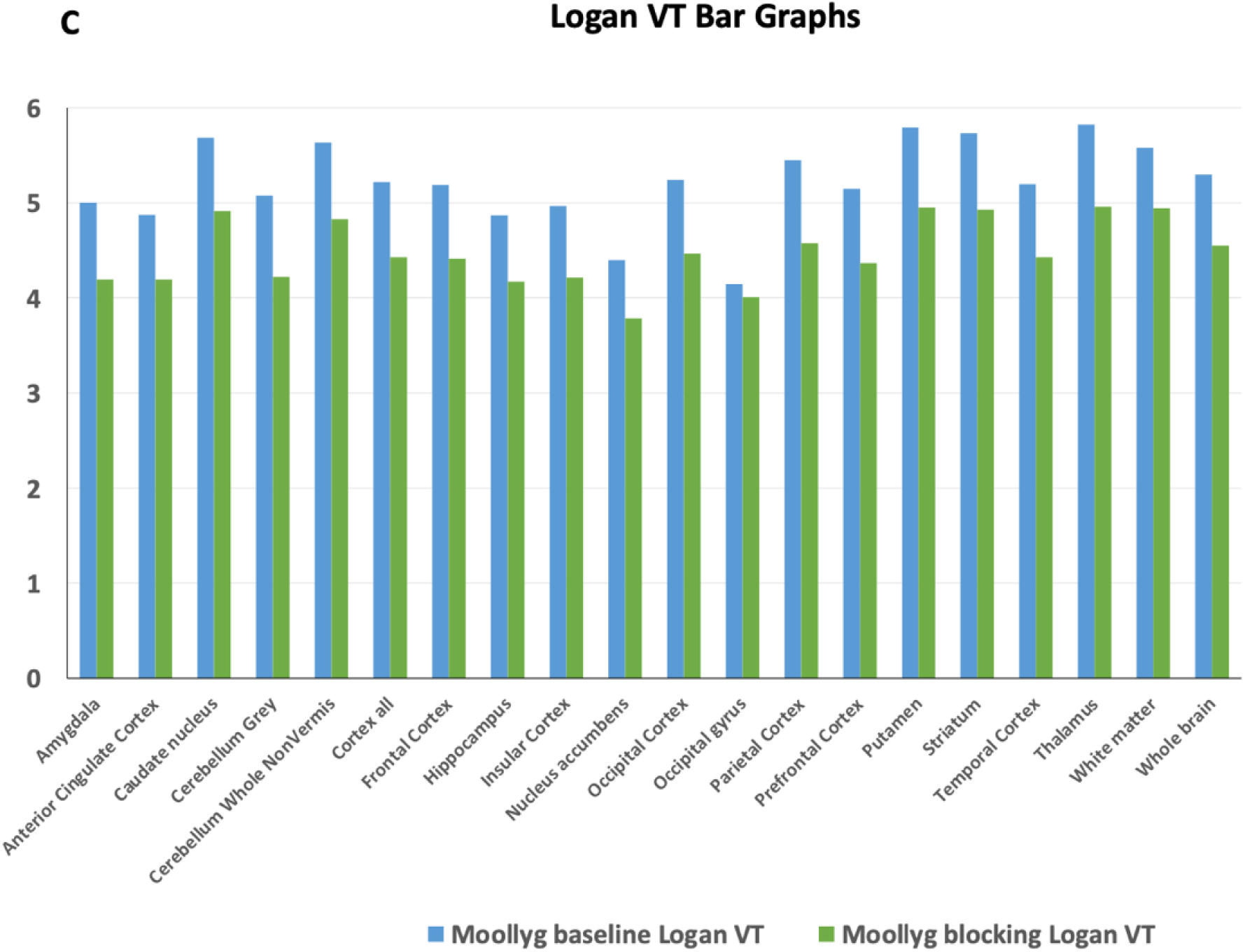
Characterization of [^11^C]**13** in the nonhuman primate brain. (**A**) 2-tissue compartment model (2T4k1v) fits in the six brain regions (left) and Logan plots (right) for [^11^C]**13** in the baseline and blocking experiments. (**B**) Structural MRI (MEMPRAGE) and [^11^C]**13** Logan *V_T_* images for the baseline (middle) and blocking studies (bottom). (**C**) Logan *V_T_* values bar graph obtained when using 120 min of data and t* of 30 min under baseline and blocking conditions.

## CONCLUSION

We have synthesized and characterized a 3,4-dihydro-2*H*-pyrano[2,3-*b*]pyridine NAM **13** as a PET imaging ligand for mGluR2. Compound **13** has a potent negative allosteric modulatory activity and suitable physiochemical properties as a PET imaging candidate. Radiolabeling of compound **13** was achieved via the *O*-methylation of phenol **24** using [^11^C]CH_3_I with a high radiochemical yield and a high molar activity. Preliminary PET imaging studies in rats confirmed the superior brain heterogeneity of [^11^C]**13**, particularly in striatum and cortex, as well as its favorable binding specificity and binding kinetics. Subsequent characterization of [^11^C]**13** in a non-human primate confirmed its capability of generating high-contrast images to map the biodistribution of mGluR2 in monkey brain. Using the 2-tissue compartment model, the accumulation of [^11^C]**13** was quantified in mGluR2-enriched brain regions, where the regional total volume of distributions (*V_T_*) was selectively reduced following the pretreatment of VU6001966 (**9**). Therefore, the experimental imaging studies conducted in two different species provided similar results in revealing the biological function of [^11^C]**13**. Altogether, [^11^C]**13** is a promising PET imaging ligand for mGluR2 to be further developed for translational studies.

## EXPERIMENTAL SECTION

All reagents and starting materials were obtained from the commercial sources including Sigma-Aldrich (St. Louis, MO), Thermo Fisher Scientific, Combi-Blocks (San Diego, CA), Ambeed (Arlington Hts, IL), and used as received. The commercially available compounds VU6001966 (**9**) and MNI-137 (**26**) were purchased from Tocris Bioscience (Minneapolis, MN). Silica gel flash column chromatography was performed using silica gel, particle size 60 Å, 230-400 mesh (Supelco). Microwave reactions were carried out in a CEM Discover microwave synthesizer. ^1^H and ^13^C nuclear magnetic resonance (NMR) spectra were collected with a JEOL 500 MHz spectrometer using tetramethylsilane (TMS) as an internal standard. All chemical shifts (δ) are assigned as parts in per million (ppm) downfield from TMS. Signals are described as s (singlet), d (doublet), t (triplet), q (quartet), or m (multiplet). Coupling constants (*J*) are quoted in hertz. Liquid chromatography-mass spectrometry (LCMS) was used to determine the mass and purity of all compounds (≥ 95%). The LCMS is equipped with a 1200 series HPLC system (Agilent Technologies, Canada), a multi-wavelength UV detector, a model 6310 ion trap mass spectrometer (Santa Clara, CA), and an analytical column (Agilent Eclipse C8, 150 mm × 4.6 mm, 5 μm). High-Resolution Mass Spectrometry (HRMS) was obtained from the Harvard Center for Mass Spectrometry at the Harvard University, Cambridge, using electrospray ionization (ESI) technique (Thermo_q-Exactive_Plus_I Mass Spectrometer).

### Chemistry

#### Methyl (E)-3-(2,6-dichloro-4-iodopyridin-3-yl)acrylate (16)

To a solution of 2,6- dichloro-4-iodonicotinaldehyde (**14**, 5.0 g, 16.56 mmol) in anhydrous tetrahydrofuran (105 mL) was added methyl 2-(triphenylphosphoranylidene)acetate (**15**, 8.31g, 24.84 mmol) under nitrogen. The mixture was stirred at 80 °C for 2h. After the reaction was completed, the solvent was evaporated under vacuum and the residue was purified by silica flash column chromatography to give the product as pale-yellow solid (14.16 mmol, 5.07 g, 85.5% yield). ^1^H NMR (500 MHz, CD_3_OD): *δ* 7.84 (s, 1H), 7.54 (d, *J* = 16.3 Hz, 1H), 6.44 (d, *J* = 16.3 Hz, 1H), 3.84 (s, 3H). ^13^C NMR (125 MHz, CDCl_3_): *δ* 165.8, 149.7, 147.6, 141.3, 133.7, 133.1, 128.1, 112.4, 52.33. HRMS (ESI^+^) for C_9_H_7_Cl_2_INO_2_^+^ [M+H]^+^ requires m/z = 357.8893, found 357.8891.

#### Methyl 3-(2,6-dichloro-4-iodopyridin-3-yl)propanoate (17)

To a solution of **16** (5.5g, 15.36 mmol) in anhydrous tetrahydrofuran /*tert*-Butanol (21 mL/21 mL) was added the RhCl(PPh_3_)_3_ (2.82g, 3.05 mmol). The mixture was stirred at room temperature under 42 psi H_2_ for 48 h. After the reaction was completed. The solvent was removed under vacuum and the residue was purified by silica flash column chromatography to give the product as white solid (3.14 g, 56.8% yield). ^1^H NMR (500 MHz, CDCl_3_): *δ* 7.74 (s, 1H), 3.72 (s, 3H), 3.27 (t, *J* = 8.4 Hz, 2H), 2.56 (t, *J* = 8.4 Hz, 2H). ^13^C NMR (125 MHz, CDCl_3_): *δ* 172.1, 148.6, 148.4, 136.4, 133.7, 114.0, 52.13, 33.1, 31.6. HRMS (ESI^+^) for C_9_H_9_Cl_2_INO_2_^+^ [M+H]^+^ requires m/z = 359.9050, found 359.9049.

#### Methyl 3-(2,6-dichloro-4-(2,4-difluorophenyl)pyridin-3-yl)propanoate (19a)

To a solution of **17** (0.5 g, 1.39 mmol) in 1,4-dioxane/water (3.0 mL/0.6 mL) was added (2,4- difluorophenyl)boronic acid (**18a**, 0.24 g, 1.53 mmol), Pd(dppf)Cl_2_ (0.10 g, 0.139 mmol), and NaHCO_3_ (0.234 g, 2.78 mmol). The mixture was stirred at 100 °C for 3 h. The solvent was removed under vacuum and the residue was purified by silica flash column chromatography to give the product as a yellow oil (0.23 g, 47.8% yield). ^1^H NMR (500 MHz, CDCl_3_): *δ* 7.17 (dd, *J* = 6.7, 14.8 Hz, 1H), 7.12 (s, 1H), 7.01 (t, *J* = 8.2 Hz, 1H), 6.96 (t, *J* = 9.1 Hz, 1H), 3.60 (s, 3H), 2.88- 2.91 (m, 2H), 2.48 (t, *J* = 7.7 Hz, 2H). ^13^C NMR (125 MHz, CDCl_3_): *δ* 172.3, 163.6 (dd, *J* = 11.8, 252.3 Hz), 159.0 (dd, *J* = 12.1, 250.5 Hz), 151.2, 148.4, 148.0, 132.6, 131.3 (dd, *J* = 4.2, 9.7 Hz), 125.0, 120.9 (d, *J* = 12.5 Hz), 112.3 (dd, *J* = 3.4, 21.4 Hz), 104.9 (t, *J* = 25.4 Hz), 51.9, 32.2, 25.4. ^19^F NMR (470 MHz, CDCl_3_): δ -106.99 (dd, *J* = 5.9, 13.0 Hz), -109.20 (dd, *J* = 7.6, 16.1 Hz). HRMS (ESI^+^) for C_15_H_12_Cl_2_F_2_NO_2_ [M+H] requires m/z = 346.0208, found 346.0208.

#### Methyl 3-(2,6-dichloro-4-(2-fluoro-4-methoxyphenyl)pyridin-3-yl)propanoate (19b)

The procedure described for compound **19a** was applied to (2-fluoro-4-methoxyphenyl)boronic acid (**18b**) to give compound **19b** as a waxy pale-yellow solid (0.383 g, 77.0% yield). ^1^H NMR (500 MHz, CDCl_3_): *δ* 7.11 (s, 1H), 7.06 (t, *J* = 8.5 Hz, 1H), 6.78 (dd, *J* = 2.2, 8.5 Hz, 1H), 6.71 (dd, *J* = 2.3, 11.5 Hz, 1H), 3.84 (s, 3H), 3.60 (s, 3H), 2.92 (t, *J* = 8.0 Hz, 2H), 2.47 (t, *J* = 8.2 Hz, 2H). ^13^C NMR (125 MHz, CDCl_3_): *δ* 172.5, 161.9 (d, *J* = 10.8 Hz), 159.5 (d, *J* = 246.9 Hz), 151.0, 149.4, 147.8, 132.8, 130.8 (d, *J* = 4.4 Hz), 125.2, 116.7 (d, *J* = 16.4 Hz), 110.8, 102.1 (d, *J* = 25.3 Hz), 55.8, 51.9, 32.2, 25.5. ^19^F NMR (470 MHz, CDCl_3_): δ -111.6 (t, J = 9.0 Hz). HRMS (ESI^+^) for C_16_H_15_Cl_2_FNO_3_^+^ [M+H]^+^ requires m/z = 358.0408, found 358.0408.

#### Methyl 3-(4-(4-(benzyloxy)-2-fluorophenyl)-2,6-dichloropyridin-3-yl)propanoate (19c)

The procedure described for compound **19a** was applied to (4-(benzyloxy)-2-fluorophenyl)boronic acid (**18c**) to give compound **19c** as a colorless oil (0.46 g, 84.0%). ^1^H NMR (500 MHz, CDCl_3_): *δ* 7.40-7.44 (m, 4H), 7.34-7.37 (m, 1H), 7.12 (s, 1H), 7.07 (t, *J* = 8.5 Hz, 1H), 6.86 (dd, *J* = 2.4, 8.5 Hz, 1H), 6.79 (dd, *J* = 2.4, 11.5 Hz, 1H), 5.09 (s, 2H), 3.60 (s, 3H), 2.93 (t, *J* = 8.2 Hz, 2H), 2.48 (t, *J* = 8.2 Hz, 2H). ^13^C NMR (125 MHz, CDCl_3_): *δ* 172.5, 161.0 (d, *J* = 10.8 Hz), 159.4 (d, *J* = 247.3 Hz), 151.0, 149.4, 147.8, 136.0, 132.8, 130.8, 128.9, 128.5, 127.7 (m), 125.2 (m), 117.0 (d, *J* = 16.6 Hz), 111.5, 103.1 (d, *J* = 25.2 Hz), 70.6, 51.9, 32.3, 25.5. ^19^F NMR (470 MHz, CDCl_3_): δ -111.4 (t, J = 10.0 Hz). HRMS (ESI^+^) for C_22_H_19_Cl_2_FNO_3_^+^ [M+H]^+^ requires m/z = 434.0721, found 434.0722.

#### 4-(2,6-dichloro-4-(2,4-difluorophenyl)pyridin-3-yl)-2-methylbutan-2-ol (20a)

To a solution of **19a** (0.23 g, 0.66 mmol) in anhydrous tetrahydrofuran (6.3 mL) was added methylmagnesium bromide (3.0 M in diethyl ether, 1.33 mL, 4.0 mmol) dropwise at 0 °C under nitrogen. The mixture was stirred at 0 °C for 1 h. After the reaction was completed, the mixture was quenched with saturated aqueous NH_4_Cl solution (30 mL) and extracted with ethyl acetate (20 mL x 3). The combined organic layers were dried over anhydrous MgSO_4_. The solvent was removed under reduced pressure and the residue was purified by silica flash column chromatography to give the product as a colorless oil (0.22 g, 95.6% yield). ^1^H NMR (500 MHz, CDCl_3_): *δ* 7.17 (td, *J* = 6.3, 8.3 Hz, 1H), 7.10 (s, 1H), 6.99 (td, *J* = 2.1, 7.9 Hz, 1H), 6.95 (td, *J* = 2.4, 9.3 Hz, 1H), 2.63 (m, 2H), 1.53 (m, 2H), 1.08 (s, 6H), 1.03 (s, 1H). ^13^C NMR (125 MHz, CDCl_3_): *δ* 163.5 (dd, *J* = 11.6, 252.1 Hz), 159.1 (dd, *J* = 11.9, 250.2 Hz), 151.0, 148.1, 147.4, 134.6, 131.3 (dd, *J* = 4.3, 9.6 Hz), 124.8, 121.1 (dd, *J* = 3.8, 16.6 Hz), 112.0 (dd, *J* = 3.4, 21.3 Hz), 104.7 (t, *J* = 25.5 Hz), 70.5, 42.0, 28.8, 25.4. ^19^F NMR (470 MHz, CDCl_3_): δ -107.27 (m), -109.00 (dd, *J* = 8.2, 16.2 Hz). HRMS (ESI^+^) for C_16_H_16_Cl_2_F_2_NO^+^ [M+H]^+^ requires m/z = 346.0572, found 346.0570.

#### 4-(2,6-dichloro-4-(2-fluoro-4-methoxyphenyl)pyridin-3-yl)-2-methylbutan-2-ol (20b)

The procedure described for compound **20a** was applied to **19b** to give compound **20b** as a colorless oil (0.28 g, 95.2% yield). ^1^H NMR (500 MHz, CDCl_3_): *δ* 7.10 (s, 1H), 7.07 (t, *J* = 8.5 Hz, 1H), 6.77 (dd, *J* = 2.4, 8.5 Hz, 1H), 6.71 (dd, *J* = 2.4, 11.5 Hz, 1H), 3.83 (s, 3H), 2.67 (t, *J* = 8.2 Hz, 2H), 1.55 (t, *J* = 8.2 Hz, 2H), 1.08 (s, 6H), 1.02 (s, 1H). ^13^C NMR (125 MHz, CDCl_3_): *δ* 161.8 (d, *J* = 10.7 Hz), 159.5 (d, *J* = 247.2 Hz), 150.8, 149.1, 147.2, 134.8, 130.8 (d, *J* = 4.1 Hz), 125.1, 117.0 (d, *J* = 16.8 Hz), 110.5, 102.0 (d, *J* = 25.2 Hz), 70.7, 55.8, 42.1, 28.8, 25.4. ^19^F NMR (470 MHz, CDCl_3_): δ -111.4 (t, *J* = 10.1 Hz). HRMS (ESI^+^) for C_17_H_19_Cl_2_FNO_2_^+^ [M+H]^+^ requires m/z = 358.0771, found 358.0771.

#### 4-(4-(4-(benzyloxy)-2-fluorophenyl)-2,6-dichloropyridin-3-yl)-2-methylbutan-2-ol (20c)

The procedure described for compound **20a** was applied to **19c** to give compound **20c** as a pale-yellow oil (0.37g, 92.5%). ^1^H NMR (500 MHz, CDCl_3_): *δ* 7.38-7.43 (m, 4H), 7.33-7.36 (m, 1H), 7.10 (s, 1H), 7.07 (t, *J* = 8.5 Hz, 1H), 6.85 (dd, *J* = 2.4, 8.5 Hz, 1H), 6.79 (dd, *J* = 2.4, 11.4 Hz, 1H), 5.09 (s, 2H), 2.64-2.68 (m, 2H), 1.51-1.54 (m, 2H), 1.07 (s, 6H), 1.01 (s, 1H). ^13^C NMR (125 MHz, CDCl_3_): *δ* 160.8 (d, *J* = 10.8 Hz), 159.4 (d, *J* = 247.2 Hz), 150.8, 149.0, 147.2, 136.0, 134.8, 130.8, 128.8 (m), 128.4 (m), 127.6 (m), 125.2 (m), 117.3 (d, *J* = 16.8 Hz), 111.3, 103.0 (d, *J* = 25.4 Hz), 70.6, 70.5, 42.1, 28.7, 25.4. ^19^F NMR (470 MHz, CDCl_3_): δ -111.3 (t, *J* = 10.0 Hz). HRMS (ESI^+^) for C_23_H_23_Cl_2_FNO_2_^+^ [M+H]^+^ requires m/z = 434.1084, found 434.1086.

#### 7-chloro-5-(2,4-difluorophenyl)-2,2-dimethyl-3,4-dihydro-2H-pyrano[2,3-b]pyridine (21a)

To a solution of **20a** (0.22 g, 0.64 mmol) in *N,N*-dimethylacetamide (10.0 mL) was added cesium carbonate (0.417 g, 1.28 mmol). The mixture was stirred at 120 °C overnight. The mixture was washed with water (30 mL) and extracted with ethyl acetate (20 mL x 3). The combined organic layers were dried over anhydrous MgSO_4_. The solvent was removed under reduced pressure and the residue was purified by silica flash column chromatography to give the product as a pale-yellow solid (0.068 g, 34.3% yield). ^1^H NMR (500 MHz, CDCl_3_): *δ* 7.20 (td, *J* = 6.3, 8.4 Hz, 1H), 6.97 (ddd, *J* = 1.2, 2.5, 8.0 Hz, 1H), 6.91 (ddd, *J* = 2.5, 8.9, 9.8 Hz, 1H), 6.79 (s, 1H), 2.50 (m, 2H), 1.75 (t, *J* = 6.7 Hz, 2H), 1.41 (s, 6H). ^13^C NMR (125 MHz, CDCl_3_): *δ* 163.3 (dd, *J* = 11.8, 251.1 Hz), 160.4, 159.3 (dd, *J* = 11.9, 238.6 Hz), 147.4, 147.3, 131.4 (dd, *J* = 4.8, 9.6 Hz), 121.4 (d, *J* = 20.2 Hz), 117.8, 113.6, 111.9 (dd, *J* = 3.5, 21.6 Hz), 104.5 (t, *J* = 25.7 Hz), 32.0, 29.8, 27.0, 20.1, 20.0. ^19^F NMR (470 MHz, CDCl_3_): δ -108.44 (m), -109.32 (m). HRMS (ESI^+^) for C_16_H_15_ClF_2_NO^+^ [M+H]^+^ requires m/z = 310.0805, found 310.0807.

#### 7-chloro-5-(2-fluoro-4-methoxyphenyl)-2,2-dimethyl-3,4-dihydro-2H-pyrano[2,3-b]pyridine (21b)

The procedure described for compound **21a** was applied to **20b** to give compound **21b** as a yellow oil (0.13 g, 48.1% yield). ^1^H NMR (500 MHz, CDCl_3_): *δ* 7.09-7.13 (m, 1H), 6.78 (s, 1H), 6.75-6.79 (m, 1H), 6.69 (d, *J* = 11.7 Hz, 1H), 3.83 (s, 3H), 2.52-2.54 (m, 2H), 1.72-1.74 (m, 2H), 1.39 (s, 6H). ^13^C NMR (125 MHz, CDCl_3_): *δ* 161.6 (d, *J* = 10.9 Hz), 160.3, 159.8 (d, *J* = 247.7 Hz), 148.4, 147.0, 131.0 (d, *J* = 5.0 Hz), 118.0, 117.4 (d, *J* = 16.7 Hz), 113.7, 110.4, 102.0 (d, *J* = 25.6 Hz), 77.2, 55.8, 32.1, 27.0, 20.2, 20.1. ^19^F NMR (470 MHz, CDCl_3_): δ -111.4. HRMS (ESI^+^) for C_17_H_18_ClFNO_2_^+^ [M+H]^+^ requires m/z = 322.1005, found 322.1007.

#### 5-(4-(benzyloxy)-2-fluorophenyl)-7-chloro-2,2-dimethyl-3,4-dihydro-2H-pyrano[2,3-b]pyridine (21c)

The procedure described for compound **21a** was applied to **20c** to give compound **21c** as a colorless oil (0.13 g, 40.4%). ^1^H NMR (500 MHz, CDCl_3_): *δ* 7.39-7.44 (m, 4H), 7.34-7.37 (m, 1H), 7.12 (t, *J* = 8.5 Hz, 1H), 6.84 (dd, *J* = 2.4, 8.5 Hz), 6.79 (s, 1H), 6.77 (dd, *J* = 2.4, 11.7 Hz, 1H), 5.09 (s, 2H), 2.54 (t, *J* = 6.4 Hz, 2H), 1.74 (t, *J* = 6.7 Hz, 2H), 1.41 (s, 6H). ^13^C NMR (125 MHz, CDCl_3_): *δ* 160.7 (d, *J* = 6.0 Hz), 160.3, 159.7 (d, *J* = 231.0 Hz), 148.3, 147.0, 136.1, 131.1 (m), 128.8 (m), 128.5 (m), 127.6 (m), 118.0, 117.7 (d, *J* = 16.7 Hz), 113.7, 111.2, 102.9 (d, *J* = 26.6 Hz), 77.2, 70.6, 32.1, 27.1, 20.1. ^19^F NMR (470 MHz, CDCl_3_): δ -111.2 (t, J = 9.2 Hz). HRMS (ESI^+^) for C_23_H_22_ClFNO_2_^+^ [M+H]^+^ requires m/z = 398.1318, found 398.1316.

#### 5-(2,4-difluorophenyl)-2,2-dimethyl-3,4-dihydro-2H-pyrano[2,3-b]pyridine-7-carbonitrile (22a)

To a solution of **21a** (30.0 mg, 0.097 mmol) in dimethylformamide (3.0 mL) was added zinc cyanide (36.0 mg, 0.306 mmol) and tetrakis(triphenylphosphine)palladium(0) (30.0 mg, 0.026 mmol) in a microwave tube. The mixture was heated to 160 °C in a microwave synthesizer (CEM, Discover SP) for 30 min. The reaction was washed with water (20 mL) and extracted with ethyl acetate (20 mL x 3). The combined organic layers were dried over anhydrous MgSO_4_. The solvent was removed under reduced pressure and the residue was purified by silica flash column chromatography to give the product as a pale-yellow waxy solid (18.0 mg, 61.8% yield). ^1^H NMR (500 MHz, CDCl_3_): *δ* 7.21 (td, *J* = 6.3, 8.4 Hz, 1H), 7.16 (s, 1H), 7.02 (td, *J* = 2.2, 8.3 Hz, 1H), 6.95 (ddd, *J* = 2.4, 8.8, 9.9 Hz, 1H), 2.63 (m, 2H), 1.80 (t, *J* = 6.7 Hz, 2H), 1.44 (s, 6H). ^13^C NMR (125 MHz, CDCl_3_): *δ* 163.6 (dd, *J* = 11.6, 252.3 Hz), 161.5, 159.3 (dd, *J* = 12.0, 251.0 Hz), 146.0, 131.3 (dd, *J* = 4.6, 9.6 Hz), 129.9, 123.2, 121.0, 120.6 (d, *J* = 16.3 Hz), 117.0, 112.3 (d, *J* = 21.3 Hz), 104.8 (t, *J* = 25.5 Hz), 77.9, 31.6, 27.1, 20.9, 20.8. ^19^F NMR (470 MHz, CDCl_3_): δ -107.32 (dd, *J* = 6.5, 13.8 Hz), -109.04 (dd, *J* = 7.4, 17.4 Hz). HRMS (ESI^+^) for C_17_H_15_F_2_N_2_O^+^ [M+H]^+^ requires m/z = 301.1147, found 301.1149.

#### 5-(2-fluoro-4-methoxyphenyl)-2,2-dimethyl-3,4-dihydro-2H-pyrano[2,3-b]pyridine-7-carbonitrile (22b)

The procedure described for compound **22a** was applied to **21b** to give compound **22b** as a colorless waxy solid (18.0 mg, 52.3% yield). ^1^H NMR (500 MHz, CDCl_3_): *δ* 7.16 (s, 1H), 7.12 (t, *J* = 8.5 Hz, 1H), 6.80 (dd, *J* = 2.4, 8.5 Hz, 1H), 6.71 (dd, *J* = 2.4, 11.8 Hz, 1H), 3.85 (s, 3H), 2.67 (t, *J* = 6.3 Hz, 2H), 1.78 (t, *J* = 6.6 Hz, 2H), 1.43 (s, 6H). ^13^C NMR (125 MHz, CDCl_3_): *δ* 162.0 (d, *J* = 10.9 Hz), 161.4, 159.8 (d, *J* = 248.0 Hz), 146.9, 130.9 (d, *J* = 4.7 Hz), 129.6, 123.6, 121.1, 117.2, 116.4 (d, *J* = 16.4 Hz), 110.7, 102.1 (d, *J* = 25.5 Hz), 77.7, 55.9, 31.8, 27.2, 21.0. ^19^F NMR (470 MHz, CDCl_3_): δ -111.1 (t, *J* = 8.8 Hz). HRMS (ESI^+^) for C_18_H_18_FN_2_O_2_^+^ [M+H]^+^ requires m/z = 313.1347, found 313.1349.

#### 5-(4-(benzyloxy)-2-fluorophenyl)-2,2-dimethyl-3,4-dihydro-2H-pyrano[2,3-b]pyridine-7-carbonitrile (22c)

The procedure described for compound **22a** was applied to **21c** to give compound **22c** was obtained as a pale-yellow solid (29.0 mg, 29.7%). ^1^H NMR (500 MHz, CDCl_3_): ^1^H NMR (500 MHz, CDCl_3_): *δ* 7.40-7.45 (m, 4H), 7.35-7.38 (m, 1H), 7.17 (s, 1H), 7.12 (t, *J* = 8.5 Hz, 1H), 6.87 (dd, *J* = 2.4, 8.5 Hz), 6.80 (dd, *J* = 2.4, 11.8 Hz, 1H), 5.10 (s, 2H), 2.66 (t, *J* = 6.6 Hz, 2H), 1.79 (t, *J* = 6.7 Hz, 2H), 1.44 (s, 6H). ^13^C NMR (125 MHz, CDCl_3_): *δ* 161.4, 161.0 (d, *J* = 11.1 Hz), 159.8 (d, *J* = 248.1 Hz), 146.9, 136.0, 130.9 (d, *J* = 5.0 Hz), 129.6, 128.9, 128.5, 127.6, 123.5, 121.0, 117.2, 116.7 (d, *J* = 16.6 Hz), 111.5, 103.1 (d, *J* = 25.5 Hz), 77.7, 70.6, 31.8, 27.2, 21.0. ^19^F NMR (470 MHz, CDCl_3_): δ -110.9 (t, J = 9.4 Hz). HRMS (ESI^+^) for C_24_H_22_FN_2_O_2_ [M+H]^+^ requires m/z = 389.1660, found 389.1656.

#### 5-(2,4-difluorophenyl)-2,2-dimethyl-3,4-dihydro-2H-pyrano[2,3-b]pyridine-7-carboxamide (12)

To a solution of **22a** (18.0 mg, 0.06 mmol) in acetone (2.0 mL) was added a solution of sodium percarbonate (43.8 mg, 0.29 mmol) in water (1.0 mL) dropwise. The mixture was stirred at room temperature for 1h. After the reaction was completed, the mixture was diluted with water (20 mL) and extracted with ethyl acetate (20 mL x 3). The combined organic layers were dried over anhydrous MgSO_4_. The solvent was removed under reduced pressure and the residue was purified by silica flash column chromatography to give the product as a white solid (11.0 mg, 57.6% yield). ^1^H NMR (500 MHz, CDCl_3_): *δ* 7.71 (s, 2H), 7.22-7.27 (m, 1H), 6.99 (td, *J* = 2.4, 8.4 Hz, 1H), 6.92 (td, *J* = 2.4, 9.4 Hz, 1H), 5.51 (s, 1H), 2.60-2.64 (m, 2H), 1.81 (t, *J* = 6.7 Hz, 2H), 1.46 (s, 6H). ^13^C NMR (125 MHz, CDCl_3_): *δ* 166.4, 163.3 (dd, *J* = 11.9, 251.0 Hz), 159.6, 159.3 (dd, *J* = 12.1, 250.3 Hz), 146.4, 146.3, 131.6 (dd, *J* = 4.6, 9.2 Hz), 121.8 (dd, *J* = 3.0, 16.4 Hz), 119.3, 117.6, 112.0 (d, *J* = 21.5 Hz), 104.4 (t, *J* = 25.7 Hz), 32.0, 29.8, 20.8. HRMS (ESI^+^) for C_17_H_17_F_2_N_2_O_2_^+^ [M+H]^+^ requires m/z = 319.1253, found 319.1253.

#### 5-(2-fluoro-4-methoxyphenyl)-2,2-dimethyl-3,4-dihydro-2H-pyrano[2,3-b]pyridine-7-carboxamide (13)

The procedure described for compound **12** was applied to **22b** to give compound **13** as a white solid (25.0 mg, 78.9% yield). ^1^H NMR (500 MHz, CDCl_3_): *δ* 7.73 (s, 2H), 7.18 (t, *J* = 8.5 Hz, 1H), 6.79 (dd, *J* = 2.3, 8.5 Hz, 1H), 6.71 (dd, *J* = 2.4, 11.8 Hz, 1H), 5.54 (s, 1H), 3.85 (s, 3H), 2.66-2.70 (m, 2H), 1.81 (t, *J* = 6.7 Hz, 2H), 1.47 (s, 6H). ^13^C NMR (125 MHz, CDCl_3_): *δ* 166.6, 161.5 (d, *J* = 10.8 Hz), 159.6, 159.8 (d, *J* = 247.5 Hz), 147.4, 146.0, 131.2 (d, *J* = 5.2 Hz), 119.4, 117.9, 117.7, 110.4, 101.9 (d, *J* = 25.7 Hz), 77.2, 55.8, 32.1, 27.2, 20.8. ^19^F NMR (470 MHz, CDCl_3_): δ -111.3 (t, *J* = 10.7 Hz). HRMS (ESI^+^) for C_18_H_20_FN_2_O_3_^+^ [M+H]^+^ requires m/z = 331.1452, found 331.1454.

#### 5-(2-fluoro-4-hydroxyphenyl)-2,2-dimethyl-3,4-dihydro-2H-pyrano[2,3-b]pyridine-7-carbonitrile (23)

To a solution of 5-(4-(benzyloxy)-2-fluorophenyl)-2,2-dimethyl-3,4-dihydro-2H-pyrano[2,3-b]pyridine-7-carbonitrile (**22c**, 22.0 mg, 0.057 mmol) in ethyl acetate (1.3 mL) was added palladium on carbon (Pd/C) (10 wt. %, 3.0 mg). The mixture was stirred at room temperature for 1h under hydrogen. After the reaction was completed, Pd/C was filtered and the solvent was removed under reduced pressure. The residue was purified by silica flash column chromatography to give the product as white solid (6.0 mg, 35.5% yield). ^1^H NMR (500 MHz, CDCl_3_): *δ* 7.17 (s, 1H), 7.08 (t, *J* = 8.3 Hz, 1H), 6.74 (dd, *J* = 2.3, 8.3 Hz, 1H), 6.70 (dd, *J* = 2.3, 11.1 Hz, 1H), 5.52 (s, 1H), 2.66 (t, *J* = 6.8 Hz, 2H), 1.79 (t, *J* = 6.7 Hz, 2H), 1.44 (s, 6H). ^13^C NMR (125 MHz, CDCl_3_): *δ* 161.5, 159.8 (d, *J* = 248.8 Hz), 158.2 (d, *J* = 11.9 Hz), 147.0, 131.1 (d, *J* = 4.8 Hz), 129.5, 123.6, 121.2, 117.1, 116.6 (d, *J* = 16.6 Hz), 112.1, 103.8 (d, *J* = 25.1 Hz), 77.8, 31.8, 27.1, 20.9. ^19^F NMR (470 MHz, CDCl_3_): δ -111.2 (t, *J* = 9.0 Hz). HRMS (ESI^+^) for C_17_H_16_FN_2_O_2_ [M+H] requires m/z = 299.1190, found 299.1191.

#### 5-(2-fluoro-4-hydroxyphenyl)-2,2-dimethyl-3,4-dihydro-2H-pyrano[2,3-b]pyridine-7-carboxamide (24)

To a solution of 5-(2-fluoro-4-hydroxyphenyl)-2,2-dimethyl-3,4-dihydro-2H- pyrano[2,3-b]pyridine-7-carbonitrile (**23**, 6.0 mg, 0.02 mmol) in acetone (0.6 mL) was slowly added sodium bicarbonate (9.5 mg, 0.06 mmol) in water (0.3 mL). The resulting mixture was stirred at room temperature for 1h. After the reaction was completed, the reaction was quenched with saturated aqueous NH_4_Cl (1.0 mL) and extracted with ethyl acetate (5.0 mL x 3). The combined organic layers were dried over anhydrous MgSO_4_. The solvent was removed under reduced pressure and the residue was purified by silica flash column chromatography to give the product as white solid (4.5 mg, 70.8% yield). ^1^H NMR (500 MHz, CDCl_3_): *δ* 7.87 (s, 1H), 7.64 (s, 1H), 7.39 (s, 1H), 7.02 (t, *J* = 8.4 Hz, 1H), 6.77 (dd, *J* = 2.3, 8.4 Hz, 1H), 6.70 (dd, *J* = 2.3, 11.4 Hz, 1H), 5.62 (s, 1H), 2.68 (m, 2H), 1.80 (t, *J* = 6.7 Hz, 2H), 1.46 (s, 6H). ^13^C NMR (125 MHz, CDCl_3_): *δ* 167.3, 159.8 (d, *J* = 247.2 Hz), 159.7, 158.7 (d, *J* = 11.9 Hz), 147.6,145.3, 131.1 (d, *J* = 5.0 Hz), 120.0, 118.2, 117.1 (d, *J* = 16.6 Hz), 112.1 (d, *J* = 1.9 Hz), 103.7 (d, *J* = 25.1 Hz), 32.0, 27.2, 20.9. ^19^F NMR (470 MHz, CDCl_3_): δ -111.8 (t, *J* = 10.3 Hz). HRMS (ESI^+^) for C_17_H_18_FN_2_O_3_ [M+H]^+^ requires m/z = 317.1296, found 317.1296.

### Radiochemistry

#### 5-(2-fluoro-4-[^11^C]methoxyphenyl)-2,2-dimethyl-3,4-dihydro-2H-pyrano- [2,3-b]pyridine-7-carboxamide ([^11^C]**13**)

[^11^C]CH_3_I was prepared from the cyclotron-generated [^11^C]CO_2_, which was produced via the ^14^N(p,α)^11^C reaction on nitrogen with 2.5% oxygen and 16 MeV protons (GE Healthcare, PETtrace). Briefly, [^11^C]CO_2_ was trapped on molecular sieves in a TRACERlab FX-CH_3_I synthesizer (GE Healthcare) and reduced to [^11^C]CH_4_ in the presence of hydrogen at 350 °C. The resulting [^11^C]CH_4_ passed through an oven containing I_2_ to afford [^11^C]CH_3_I via a radical reaction. [^11^C]CH_3_I was then transferred under helium gas to a 5 mL V-vial containing precursor **24** (0.4 ± 0.1 mg), an aqueous 0.5N NaOH (3 μL) and anhydrous DMF (350 μL). After the transfer was completed, the mixture was heated at 80 °C for 3 min. The reaction was then quenched by adding 1.0 mL of water and purified using a semi-preparative HPLC system equipped with a Waters XBridge C18 column (250 × 10 mm, 5 μ), a UV detector (wavelength = 254 nm) and a radioactivity detector. The product was eluted with a mobile phase of acetonitrile/water/Et_3_N (50/50/0.1%) at a flow rate of 5 mL/min. The fractions containing [^11^C]**13** (t_R_ = 8.6 min) were collected into a large dilution flask, which was pre-loaded with 23 mL of sterile water for injection, USP. The diluted solution was loaded onto a C18 light cartridge (Waters; pre-activated with 8 mL of EtOH followed by 16 mL of water) and the cartridge was washed with 10 mL of sterile water to remove traces of salts, residual acetonitrile and Et_3_N. [^11^C]**13** was then released from the cartridge via 0.6 mL of dehydrated ethyl alcohol (USP) followed by 5.4 mL of 0.9% sodium chloride solution (USP) into a product collection vessel. The formulated [^11^C]**13** solution was filtered through a vented sterilizing filter (Millipore-GV 0.22μ, EMD Millipore) into a 10 mL vented sterile vial for injection. The synthesis time was ca. 45 min from end-of-bombardment. The chemical and radiochemical purities of [^11^C]**13** were determined by a HPLC system (UltiMate 3000) equipped with an analytical column (Waters, XBridge, C18, 3.5 μ, 4.6 × 150 mm), a UV detector (λ = 254 nm) and a radioactivity detector. The mobile phase of acetonitrile/water/Et_3_N (45/55/0.1%) was used and the flow rate was 1 mL/min. The identity of [^11^C]**13** was confirmed by the co-injection with unlabeled compound **13**.

### Pharmacology

The negative allosteric modulatory activity was determined following a standard protocol by Eurofins Discovery. Briefly, 20 μL of 10k CHO cells/well in CP24™ were seeded into white walled, 384-well microplates and incubated at 37°C/5% CO_2_ overnight. On day of testing, media is exchanged for 10 μL of HBSS/10 mM HEPES. Intermediate dilution of compounds was performed to generate 4x stocks in HBSS/10 mM HEPES and 5 μL of 4x sample is added to the cell plate. Cells were incubated at 37°C for 15 minutes. Then, 5 μL of 4x Forskolin and 4x EC_80_ of the challenge agonist glutamate were added and cells were incubated for 30 min at 37°C. The concentration of Forskolin was 15 μM and the concentration of glutamate was 8.9 μM. Assay signal was generated through incubation with 5 μL of cAMP XS+ Ab reagent and 20 μL cAMP XS+ ED/CL lysis cocktail for one hour followed by incubation with 20 μL cAMP XS+ EA reagent for two hours at room temperature. Plates were read following signal generation with a PerkinElmer Envision^TM^ instrument for chemiluminescent signal detection. The signal is normalized to EC_80_ response (0%) and basal signal (100%). The NAM activity was analyzed using CBIS data analysis suite (ChemInnovation, CA). Percentage inhibition was calculated using the formula of % Inhibition = 100% x (mean RLU of test sample-mean RLU of EC_80_ control)/ (mean RLU of forskolin positive control - mean RLU of EC_80_ control). The assay was run in duplicate.

### Molecular modeling

The mGluR2 receptor model was built in YASARA^41^ from 17 initial models based on the crystal structures of the human metabotropic glutamate receptor 5 (PDBID:4OO9),^53^ human metabotropic glutamate receptor 1 (PDBID:4OR2),^54^ metabotropic glutamate receptor 5 apo form (PDBID:6N52),^55^ and an mGluR2 structure (PDBID: 5KZN).^56^ The model was further validated by several structural analysis tools from SAVES containing VERIFY3D,^57^ ERRA,^58^ QMEAN,^59^ and ModFOLD^60^ (Supporting Information, Section 2). The key interacting residues were predicted by Partial Order Optimum Likelihood (POOL),^61^ which include the previously reported interacting residues of Phe623, Arg635, Phe643, His723, and Asn735.^49, 62^ Compounds **12** and **13** were optimized and converted into PDB format in Avogadro 1.2 before docking.^63^ Molecular docking was performed into the model structure using Extra precision Induced Fit Docking in Glide.^64–66^

### Physiochemical Properties

#### Partition coefficient (LogD_7.4_)

The LogD_7.4_ was measured by mixing a test compound (0.1 mg) with *n*-octanol (1.0 mL) and PBS buffer (1.0 mL) at pH 7.4 in an Eppendorf tube.^67^ The tube was vortexed for 1 min before shaken at 37 °C overnight. The amount of the test compound in each phase was determined from the area under the peak at a wavelength of 254 nm in the HPLC system (UltiMate 3000). The compound was eluted with acetonitrile/water/Et_3_N (45/55/0.1%) at a flow rate of 1.0 mL/min with a Waters XBridge C18 column (250 × 10 mm, 5μ). The LogD_7.4_ was calculated by Log([compound in octanol]/[compound in PBS]). The assay was repeated at least three times for each compound.

#### Rat plasma stability

The rat plasma stability was determined by our previously described method.^22, 68^ Briefly, the test compound (2.5 μL, 1 mM DMSO stock solution) was mixed with an aliquot of rat serum (100 μL, Abcam, Inc.) in an Eppendorf tube. The tube was vortexed and incubated at 37 °C for 0 min and 60 min, separately, before the addition of 250 μL ice-cold acetonitrile. The resulting mixture was centrifuged at 10, 000g for 20 min and the supernatant was collected for analysis on the HPLC system (UltiMate 3000). The same analytical conditions were used as those in the LogD_7.4_ assay. The plasma stability value was expressed as (peak area at 60 min)/(peak area at 0 min) x 100%. The assay was repeated at least three times for each compound. Compound **24** was used as internal standard.

#### Rat liver microsome stability

The rat liver microsome stability was measured by our previously described method. ^22, 69^ Briefly, 1.5 μL of 1 mM compound solution in DMSO was added to an Eppendorf tube containing 432 μL of PBS buffer. The tube was kept at 37 °C for 10 min before a 13 μL aliquot of the Sprague-Dawley rat liver microsome (Sigma-Aldrich, No. M9066) was added. The tube was vortexed before shaken at 37 °C for 5 min. The NADPH (50 μL, 10 mM in PBS solution) was added and the resulting mixture was incubated at 37 °C for 0 min and 60 min, separately, before the addition of 250 μL of ice-cold acetonitrile. The mixture was centrifuged at 10, 000g for 20 min and supernatant was collected for analysis on the HPLC system (UltiMate 3000). The same analytical conditions were employed as those in the LogD_7.4_ assay. The liver microsome stability value was expressed as (peak area at 60 min)/(peak area at 0 min) x 100%. The assay was repeated at least three times for each compound. Compound **24** was used as internal standard and *N*-(4-chloro-3-methoxyphenyl)pyridine-2-carboxamide (ML128) was employed as positive control.

#### Pgp-Glo^TM^ assay

The Pgp-Glo™ assay was performed by following our previously described method^22^ and using the manufacturer’s instructions (Promega, Co. USA). Briefly, 25 μg of Pgp membrane (Promega, Cat. # V3601) was added to a 96-well plate (Thermo Lab systems, Cat. # 9502887) containing untreated samples, Na_3_VO_4_ (100 μM), Verapamil (100 μM), and tested compounds (100 μM). The Pgp ATPase reaction was activated by adding a solution of MgATP in the assay buffer (5 mM). After a brief mixing, the 96-well plate was placed in a 37 °C incubator for 40 min. The assay was then treated with 50 μL ATP detection solution and incubated at room temperature for 20 min to develop luminescent signal. The luminescence was read on an *in vivo* imaging system (IVIS^®^ Spectrum, PerkinElmer, USA). The change in luminescence relative to the Na_3_VO_4_ samples represents the Pgp ATPase activity with a unit of photon per second (p/s). The assay was repeated at least three times for each compound.

### PET imaging studies in rats

PET imaging experiments and data analysis of [^11^C]**13** in rats were performed by our previously described methods.^22^ Briefly, the imaging studies were carried out in Triumph II Preclinical Imaging System (Trifoil Imaging, LLC, Northridge, CA). Six normal Sprague Dawley rats (male, 285-421 g) were used which resulted in eight imaging studies comprising four baseline studies, two pretreatment studies with VU6001966 (**9**), and two blocking experiments with MNI-137 (**26**). For the imaging studies, rats were anesthetized with isoflurane (1.0-1.5%) and oxygen (1-1.5 L/min) and the vital signs, such as heart rate and breathing, were monitored. The data acquisition for 60 min started from the injection of [^11^C]**13** (63.0-87.3 MBq, iv.) through the tail vein using a catheter. The blocking agent **9** (0.5 mg/kg) was dissolved in a solution of 10% ethanol and 5% Tween-80 in 85% saline (0.1 mg/mL) while **26** (0.2 mg/kg, iv.) was formulated into a solution of 10% DMSO and 5% Tween-20 in 85% PBS (0.25 mg/mL). The blocking agents were administered 1 or 20 min before the tracer injection.

After each PET acquisition, a CT scan was performed to provide anatomical information and data for attenuation correction. The list mode PET data were reconstructed to twenty-four dynamic volumetric images (9x20s, 7x1min, 6x5min, 2×10min) via the maximum-likelihood expectation-maximization (MLEM) algorithm with 30 iterations. The ROIs, i.e., striatum, frontal cortex, cingulate cortex, hippocampus, hypothalamus, thalamus, and cerebellum were drawn onto coronal PET slices according to the rat brain atlas. The time activity curves for these ROIs were generated by PMOD 3.2 (PMOD Technologies Ltd., Zurich, Switzerland).

### PET imaging studies in a nonhuman primate

PET imaging experiments, arterial blood sampling, and data analysis of [^11^C]**13** in a cynomolgus monkey (Macaca fascicularis) (5.0 kg, female) were done by our previously described methods.^22^

#### PET imaging

The PET scans were performed in a Discovery MI (GE Healthcare) PET/CT scanner. Prior to each study, the monkey was sedated with ketamine/xylazine (10/0.5 mg/kg IM) and maintained under anesthesia with a flow of isoflurane (1-2%) in oxygen. A CT scan was done before each PET acquisition to verify anatomical location and get data for attenuation correction. The PET data acquisition started immediately at the start of a 3-minute tracer infusion and lasted for 120 min. Radiotracer activity injected at baseline and blocking studies was 190.55 MBq (A_m_ = 288.5 GBq/µmol) and 239.39 MBq (A_m_ = 135.6 GBq/µmol). The blocking agent, **9** (1.0 mg/kg, iv.) was administered 20 min before tracer injection. After the PET scan, the acquired PET data were reconstructed via a 3D time-of-flight iterative reconstruction algorithm with 3 iterations and 34 subsets. The data were also corrected for photon attenuation and scatter, radioactive decay, system dead time, detector inhomogeneity and random coincident events. The list mode PET data were framed to fifty-four dynamic volumetric images (6x10, 8x15, 6x30, 8x60, 8x120 and 18x300s) with voxel dimensions of 256 x 256 x 89 and voxel sizes of 1.17 x 1.17 x 2.8 mm^3^.

#### Arterial blood sampling and analysis

Prior to radiotracer injection, a 3-mL arterial blood sample was drawn to determine the plasma protein binding of [^11^C]**13**. Briefly, the blood sample was centrifuged and an aliquot of the supernatant was spiked with [^11^C]**13** in PBS to 22.2 MBq/mL. The resulting solution was inculcated for 10-15 min before centrifugation with the Centrifree Ultrafiltration Devices (Millipore Sigma). Aliquots of the ultrafiltrate (C_free_) and the plasma mixture (C_total_) were measured for radioactive concentration in a Wallac Wizard 2480 gamma counter. This process was performed in triplicate to determine the plasma free fraction (*f_p_*) of [^11^C]**13**.

Upon PET data acquisition, twenty three arterial blood samples were drawn by sampling every 30 seconds for the first 5 minutes followed by a decreased frequency of every 15 minutes till the end of the scan. The plasma samples were obtained as supernatant of the centrifugated whole-blood samples. The metabolism of [^11^C]**13** was evaluated using selected plasma samples from 5, 10, 15, 30, 60, 90, and 120 minutes. The amount of the intact [^11^C]**13** in plasma samples were measured by the previously described automated column switching radioHPLC system.^70–71^ Briefly, the plasma sample was trapped on a capture column (Waters Oasis HLB 30 μm) with a mobile phase of water: acetonitrile (99:1) at 1.8 mL/min (Waters 515 pump). After 4 minutes, the sample was transferred to an analytical column (Waters XBridge BEH C18, 130 Å, 3.5 μm, 4.6 mm x 100 mm) by backflushing the catch column with a mobile phase of acetonitrile: 0.1M ammonium formate in water (45:55) at 1 mL/min (Waters 515 pump) with 0.1% of TFA (pH 2.5). The eluent from the analytical column was collected in 1-minute intervals and the radioactivity was measured to determine the parent fraction in plasma (%PP) with a Wallac Wizard 2480 gamma counter. The radioactivity concentration (C(t)) measured from the well counter was expressed as kBq/cc. Therefore, the radioactivity time courses using standardized uptake value (SUV) was calculated as SUV(t) = C(t)/(ID/BW), where ID standards for injected dose in MBq and BW means body weight in kg. The time courses of %PP(t) were fitted with a sum of two decaying exponentials plus a constant. The resulting model fit and the (C_total_(t)) in plasma were multiplied to derive the metabolite-corrected arterial input function for kinetic modeling.^24, 72^

#### Image processing and analyses

All PET data were processed with an in-house developed MATLAB software that uses FSL.^73^ The PET images were first co-registered to the structural T1- weighted magnetization-prepared rapid gradient-echo (MEMPRAGE) images, which were aligned into an (magnetic resonance) MR monkey template space.^74^ The resulting transformation was then applied to PET images. Regional TACs were extracted from the native PET image space for specific ROIs.

The extracted TACs were modeled via the reversible one- (1T) and two- (2T) tissue compartment model configurations with the metabolite-corrected arterial plasma input function. The 2T model was assessed in its irreversible (k_4_ = 0) and reversible configurations. A fixed vascular contribution of the WB radioactivity to the PET signal was set to 5%. The kinetic parameters were estimated using the nonlinear weighted least-squares fitting and the frame durations were chosen for the weights. Regional total volume of distributions (*V_T_*) were calculated from the estimated microparameters following the consensus nomenclature reported by Innis *et al*.^75^ The stability of *V_T_* estimates was assessed by progressively truncating the PET data in 10 min increment from the full duration of 120 min to 60 min. Additionally, the Logan graphical analysis technique was also assessed to generate *V_T_* estimates with different cutoff time t*.^76^

## ASSOCIATED CONTENT

### Supporting Information

The Supporting Information is available free of charge on the ACS Publications website at DOI: Molecular formular strings (CSV)

Preparation and validation of mGluR2 NAM model, POOL prediction of key residues, semipreparative HPLC purification and analytical HPLC characterization of [^11^C]**13**, and ^1^H, ^13^C NMR spectra for synthesized compounds (PDF)

The PDB coordinates of the mGluR2 NAM model (PDB)

## AUTHOR INFORMATION

### Author Contributions

The manuscript was written through contributions of all authors.

### Notes

The authors declare no conflict interest.

## ACKNOWLEDGEMENT

This project was funded by NIH grants [R01EB021708, R01NS100164, 1S10RR023452-01 and 1S10OD025234-01] for the imaging instrumentation and characterization of the organic compounds. This projected was supported by the NIH grants [S10OD018035 and P41EB022544] for the blood counting and metabolite analysis equipment.

## ABBREVIATIONS USED

CNS: central nervous system
mGluR2: metabotropic glutamate receptor 2
PET: positron emission tomography
PAM: positive allosteric modulator
NAM: negative allosteric modulator
BBB: blood-brain barrier
7-TM: seven transmembrane
Pgp-BCRP: P-glycoprotein and the breast cancer resistance protein, P-glycoprotein
CHO: chinese hamster ovary
DMF: dimethylformamide
DMA: dimethylacetamide
EOS: end of synthesis
LCMS: Liquid chromatography-mass spectrometry
NHP: non-human primate
ROI: region of interest
TAC: time-activity curve
*f_p_*: plasma free fraction
WB: whole-blood
PL: plasma
AIC: Akaike information criteria
NMT: NIMH macaque template
SUV: standardized uptake value
*V_T_*: regional total volume of distribution
MEMPRAGE: magnetization-prepared rapid gradient-echo
USP: United States Pharmacopeia
MLEM: maximum-likelihood expectation-maximization.

