## Supporting information for "Synthesis and characterization of 5-(2-fluoro-4-[^11^C]methoxyphenyl)-2,2-dimethyl-3,4-dihydro-2*H*-pyrano[2,3-*b*]pyridine-7-carboxamide as a PET imaging ligand for metabotropic glutamate receptor 2"

**Table of Contents**

1. Preparation of mGluR2 homology model for NAMs……………………….……………S3

2. Structural evaluation of mGluR2 model…………………………………………….…S4

   2.1 MODFOLD results…………………………………………………………….….S4

   2.2 Structure Analysis Verification Server (SAVES) results………………………....S5

   2.3 SWISS Model-QMEAN results…………………………………………………...S6

   2.4 Ramachandran Plot…………………………………………………………...….S7

3. Prediction of Binding Site……………………………………………………………...S8

4. Purification and confirmation of [$^{11}$C]**13**………………..……………………..……..S9

5. Prediction of the metabolism sites of **13** with SMARTCyp …..................…………..…S11

6. NMR spectra……………………………………………………………………….....S12

7. References……………………………………………………………………...….S48

**Supporting Information:**

### 1. Preparation of mGluR2 homology model for NAMs:

The target sequence having 872 residues used for building the model for mGluR2 was listed below:

MGSLLALLALLLLWGAVAEGPAKKVLTLEGDLVLGGLFPVHQKGGPAEDCGPVNEHRGIQ

RLEAMLFALDRINRDPHLLPGVRLGAHILDSCSKDTHALEQALDFVRASLSRGADGSRHI

CPDGSYATHGDAPTAITGVIGGSYSDVSIQVANLLRLFQIPQISYASTSAKLSDKSRYDY

FARTVPPDFFQAKAMAEILRFFNWTYVSTVASEGDYGETGIEAFELEARARNICVATSEK

VGRAMSRAAFEGVVRALLQKPSARVAVLFTRSEDARELLAASQRLNASFTWVASDGWGAL

ESVVAGSEGAAEGAITIELASYPISDFASYFQSLDPWNNSRNPWFREFWEQRFRCSFRQR

DCAAHSLRAVPFEQESKIMFVVNAVYAMAHALHNMHRALCPNTTRLCDAMRPVNGRRLYK

DFVLNVKFDAPFRPADTHNEVRFDRFGDGIGRYNIFTYLRAGSGRYRYQKVGYWAEGLTL

DTSLIPWASPSAGPLPASRCSEPCLQNEVKSVQPGEVCCWLCIPCQPYEYRLDEFTCADC

GLGYWPNASLTGCFELPQEYIRWGDAWAVGPVTIACLGALATLFVLGVFVRHNATPVVKA

SGRELCYILLGGVFLCYCMTFIFIAKPSTAVCTLRRLGLGTAFSVCYSALLTKTNRIARI

FGGAREGAQRPRFISPASQVAICLALISGQLLIVVAWLVVEAPGTGKETAPERREVVTLR

CNHRDASMLGSLAYNVLLIALCTLYAFKTRKCPENFNEAKFIGFTMYTTCIIWLAFLPIF

YVTSSDYRVQTTTMCVSVSLSGSVVLGCLFAPKLHIILFQPQKNVVSHRAPTSRFGSAAA

RASSSLGQGSGSQFVPTVCNGREVVDSTTSSL

A hybrid model was generated in YASARA[1] from the above sequence and the template structures with the PDB IDs, 4OO9,[2] 4OR2,[3] 6N52[4] and 5KZN[5]. These structures were obtained after a BLAST search against the PDB of the above mGluR2 sequence.[6] YASARA generated 17 models initially from these structures and finally a hybrid model was generated using the best parts from these 17 initial models, to increase the accuracy beyond each contributor. Figure S1 shows the hybrid model generated in YASARA with initial model in blue and hybridized parts in a different color. The resulting hybrid model obtained the following quality Z-scores (Table S1).

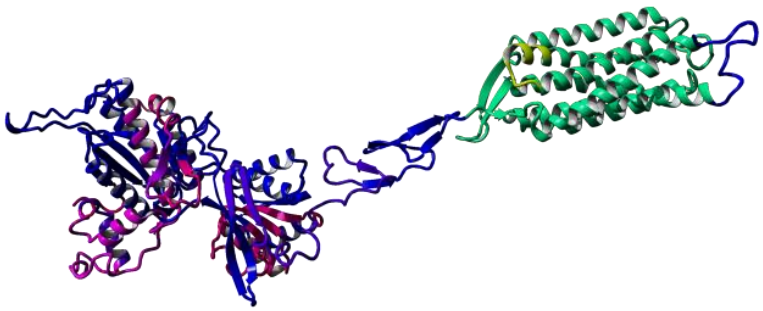

**Figure S1**: The figure shows the initial model in blue, and all hybridized parts in different colors

**Table S1**: Z-Scores for the hybrid model generated on YASARA

| Check type | Quality Z-score | Comment |
|---|---|---|
| Dihedrals | 0.043 | Good |
| Packing 1D | -0.082 | Good |
| Packing 3D | -1.216 | Satisfactory |
| Overall | -0.591 | Good |

### 2. Structural evaluation of mGluR2 model

This hybrid model was further validated by the following methods.

#### 2.1 MODFOLD results

The model generated was validated using ModFOLD.[7] The confidence and P-value for this model is HIGH: 1.001 E-3 with the global model quality score of 0.4433, indicating it a complete and confident model for mGluR2. The p-value represents the probability of each model being incorrect. The p-value for this model is 0.001001, meaning there is only a 1/1001 chance of this model being incorrect.

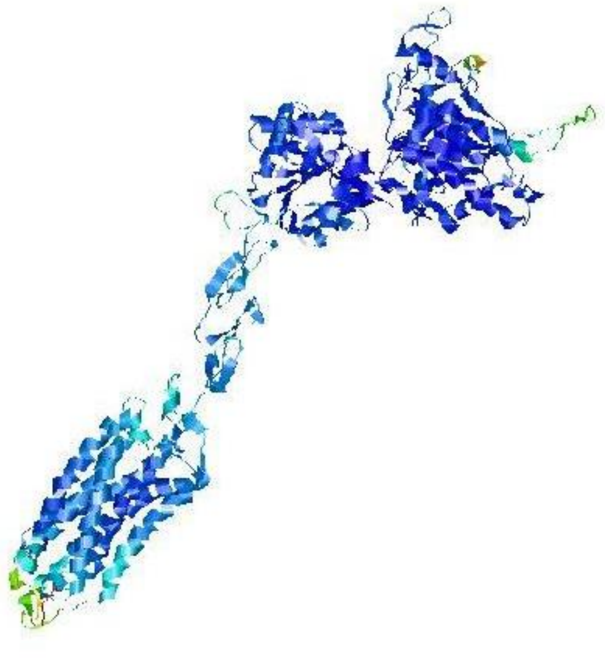

**Figure S2**: This image was generated by ModFOLD based on residue accuracy prediction for the model. Blue is for high accuracy through green, yellow, orange to red which is for low accuracy.

**2.2 Structure Analysis Verification Server (SAVES) results**

The second server used to validate this model was SAVES[8-10] and its components, VERIFY 3D and ERRAT. VERIFY 3D scores as a function of sequence number for the model. As Figure S3 shows, VERIFY 3D assigned a 3D-1D score of > 0.2 for at least 87.23% of the amino acids. This implies that the model is compatible with its sequence.

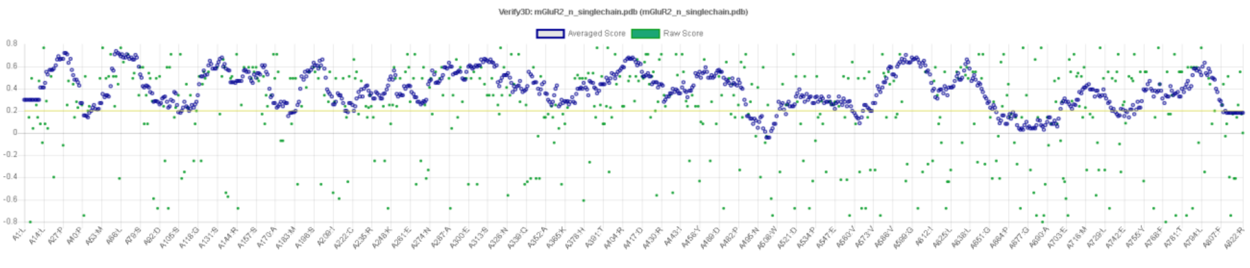


**Figure S3**: Verify 3D scores for the hybrid model

ERRAT scores as a function of sequence number for the hybrid model generated by YASARA. As shown in Figure S4, green indicates a good score, yellow represents regions that can be rejected at 95% confidence and red symbolizes regions that can be rejected at 99% confidence. As Figure S4 shows, the hybrid model contains significantly low red colored regions. The quality factor for this model is 95.9. Therefore, it is a good model according to ERRAT.

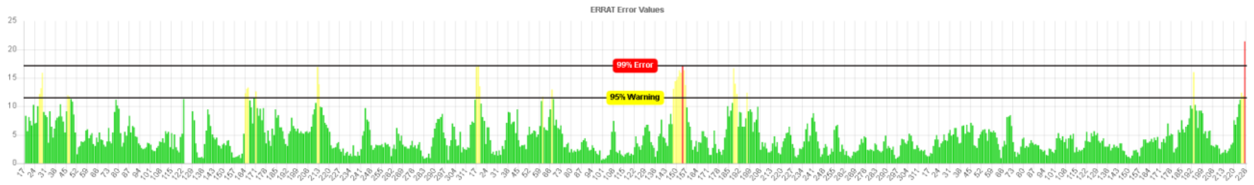


**Figure S4**: ERRAT scores for the hybrid model

**2.3 SWISS Model-QMEAN results**

QMEAN is a composite scoring function which derives both global and local absolute quality estimates based on one single model.[9] The QMEAN score for this hybrid model is -1.43. Below is an image showing the sequence of the protein colored by local quality.

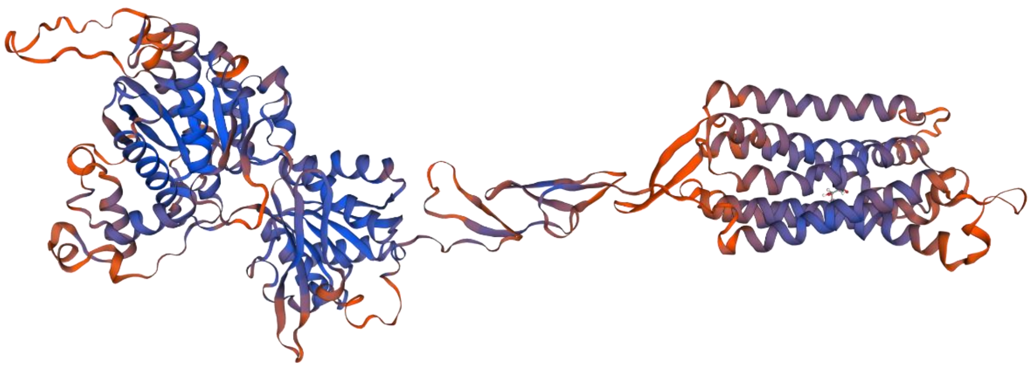

**Figure S5**: Image generated by QMEAN showing the local quality of the hybrid model. Blue indicates better quality regions; orange indicates lower quality regions

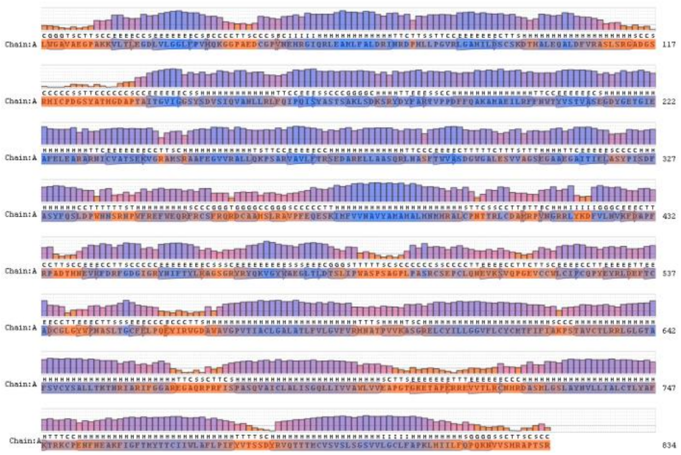

**Figure S6**: Image showing the local quality of the structure as a function of sequence number, generated by QMEAN.

### 2.4 Ramachandran Plot:

The hybrid model generated by YASARA was further evaluated via Ramachandran plot. As Figure S7 shows 89.6% (643) of the residues lie in the favored regions and 10.2% (73) lie in the additionally allowed regions. There are 0.1% (1) residues in the generously allowed regions and no residues in disallowed regions. This is further evidence of a quality model structure.

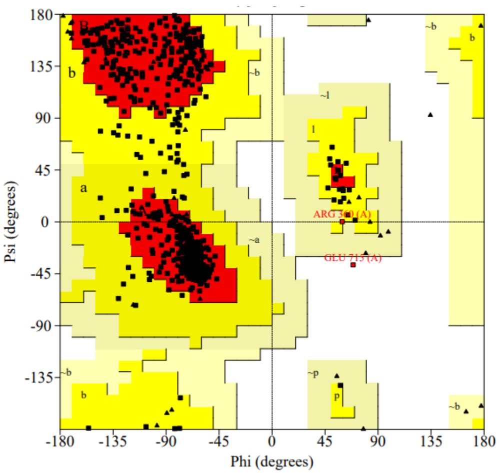


**Figure S7:** A Ramachandran plot for the hybrid model. Plot generated with the SAVES server.

### 3. Prediction of Binding Site

Partial Order Optimum Likelihood (POOL)[11] was used to predict the key binding residues in the allosteric binding site. The identified top residues are listed below and the ones that are reported previously are highlighted in bold.[12-13]

Cys560, Cys606, Leu609, Cys616, Tyr617, **Phe623**, **Arg635**, Arg636, Gly638, Leu639, Gly640, Thr641, **Phe643**, Val645, Cys646, Tyr647, Leu650, Lys653, Cys683, **His723**, Tyr734, **Asn735**, Ile739, Cys742, Tyr781, Tyr787, Cys795, Val798, Ser801, Lys813.

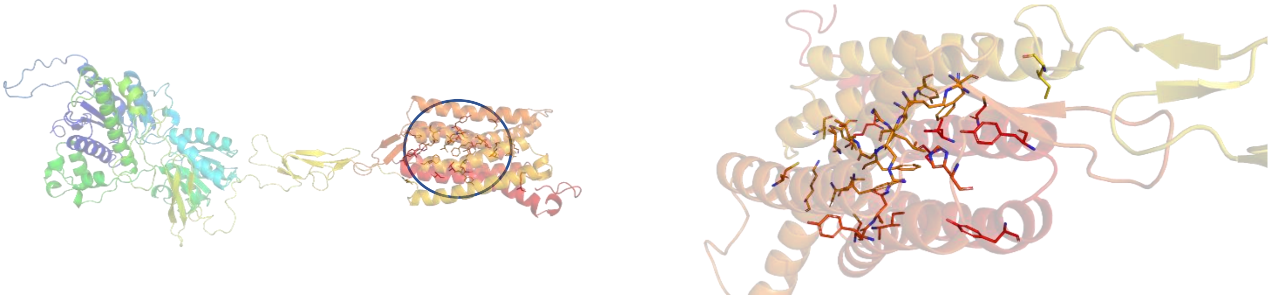

**Figure S8**: Position of the allosteric binding site for NAMs.

### 4. Purification and confirmation of [$^{11}$C]13

Figure S9 shows the semi-preparative HPLC spectra for the purification of [$^{11}$C]**13** from the reaction mixture. The HPLC radioactivity trace is shown at the top and the UV trace is shown at the bottom. The retention time of [$^{11}$C]**13** was 8.62 min under the following HPLC conditions: Column: Waters XBridge, C18, 250 × 10 mm, 5 μ; Wavelength of 254 nM; Mobile phase: acetonitrile/water/$Et_3N$ (50/50/0.1%) at a flow rate of 5 mL/min.

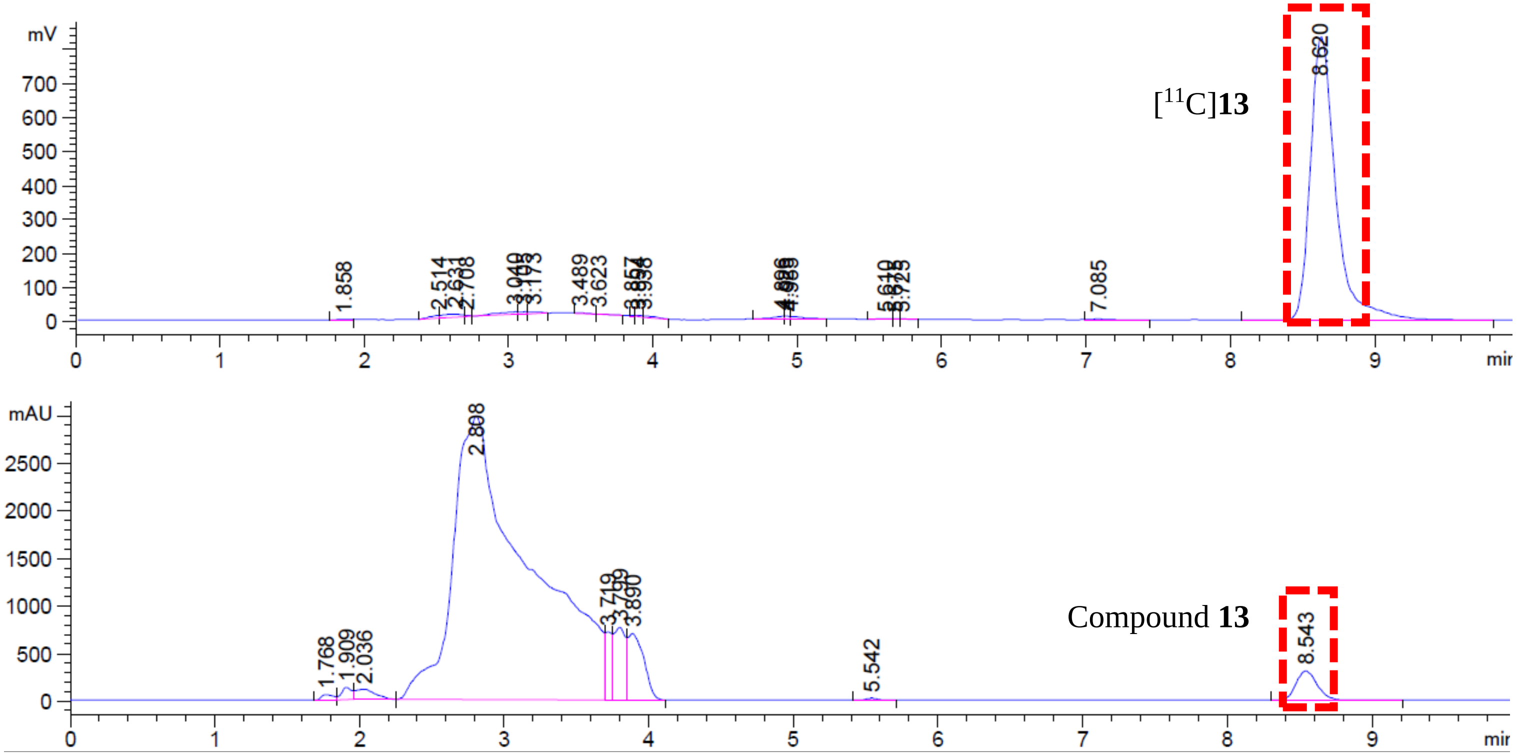


**Figure S9**. Purification of [$^{11}$C]**13** from reaction mixture via semi-preparative HPLC

The identity of the purified and formulated [$^{11}$C]**13** was confirmed by co-injecting with the unlabeled compound **13** in an analyticl HPLC system (Figure S10). The radioHPLC trace of [$^{11}$C]**13** spectrum is shown in black (top) and the UV Trace of reference **13** is shown in blue (bottom). The retention time of [$^{11}$C]**13** was 8.11 min under the following HPLC conditions: Column: Waters, XBridge, C18, 4.6 × 150 mm 3.5 μ; Wavelength of 254 nM; Mobile phase: acetonitrile/water/$Et_3N$ (45/55/0.1%) at a flow rate of 1 mL/min.

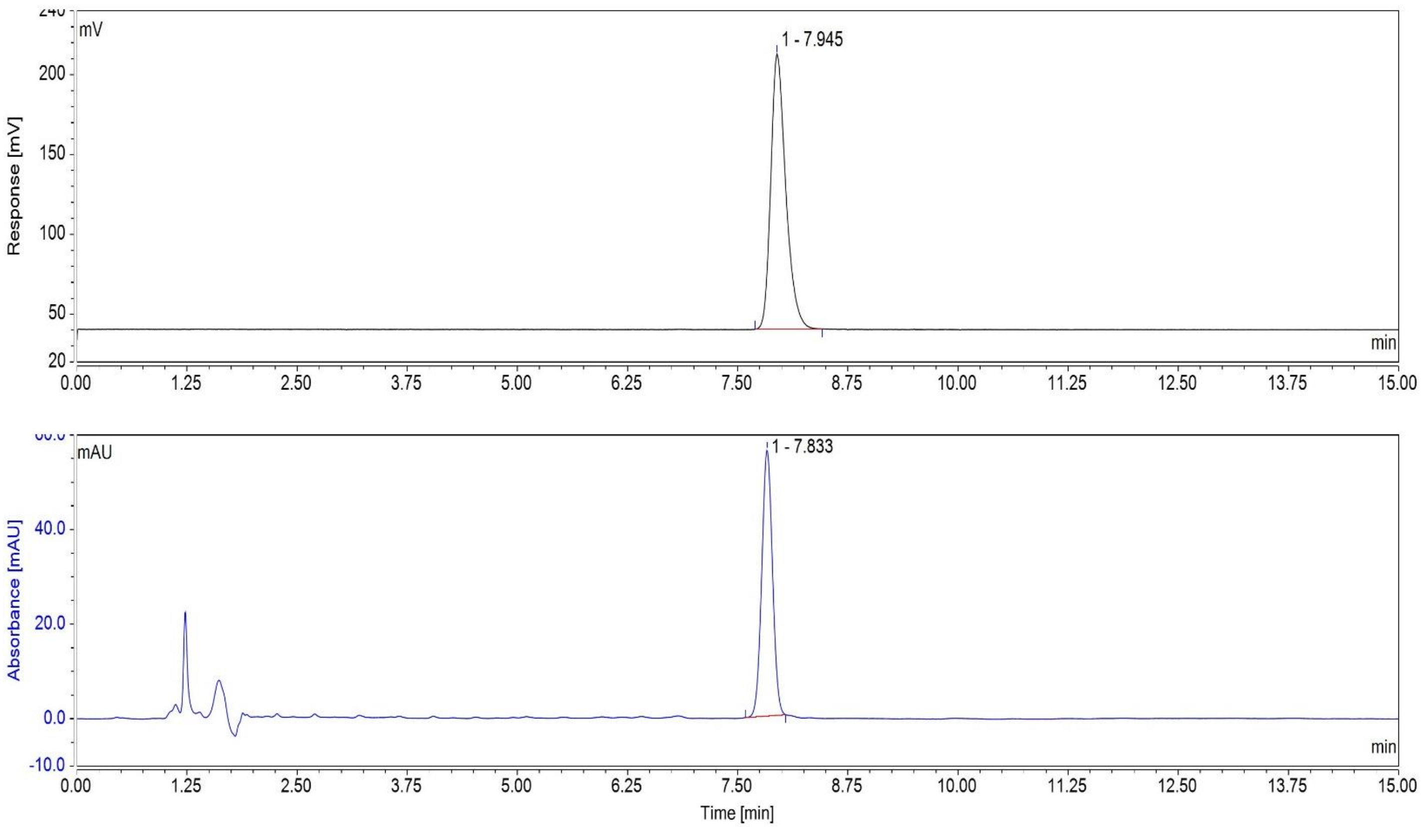


**Figure S14**. Analytical HPLC Spectra for formulated [$^{11}$C]**13**

### 5. Prediction of the metabolism sites of 13 with SMARTCyp[14]

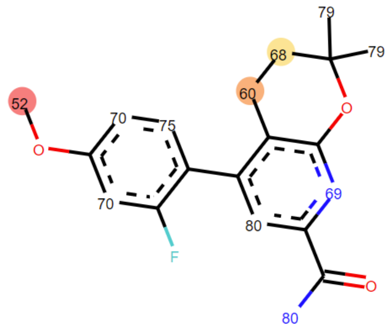


| 3A4 Ranking | Atom | 3A4 Score | Energy | 2DSASA | Span2end | Relative Span | Similarity |
|---|---|---|---|---|---|---|---|
| 1 | C.22 | 51.5 | 62.2 | 66.4 | 0 | 1.0 | 1.0 |
| 2 | C.18 | 59.6 | 66.4 | 25.1 | 3 | 0.7 | 0.3 |
| 3 | C.17 | 68.2 | 75.9 | 28.4 | 2 | 0.8 | 0.3 |
| 4 | N.6 | 68.6 | 75.6 | 10.9 | 2 | 0.8 | 0.3 |
| 5 | C.9 | 70.3 | 77.2 | 27.8 | 3 | 0.7 | 0.7 |
| 6 | C.11 | 70.3 | 77.2 | 26.2 | 3 | 0.7 | 0.3 |
| 7 | C.8 | 74.8 | 80.8 | 23.3 | 4 | 0.6 | 0.7 |
| 8 | C.19 | 79.3 | 89.6 | 58.5 | 0 | 1.0 | 0.3 |
| 9 | N.24 | 80.3 | 89.6 | 50.6 | 1 | 0.9 | 0.7 |
| 10 | C.2 | 80.5 | 86.3 | 17.3 | 4 | 0.6 | 0.3 |

### 6. NMR spectra

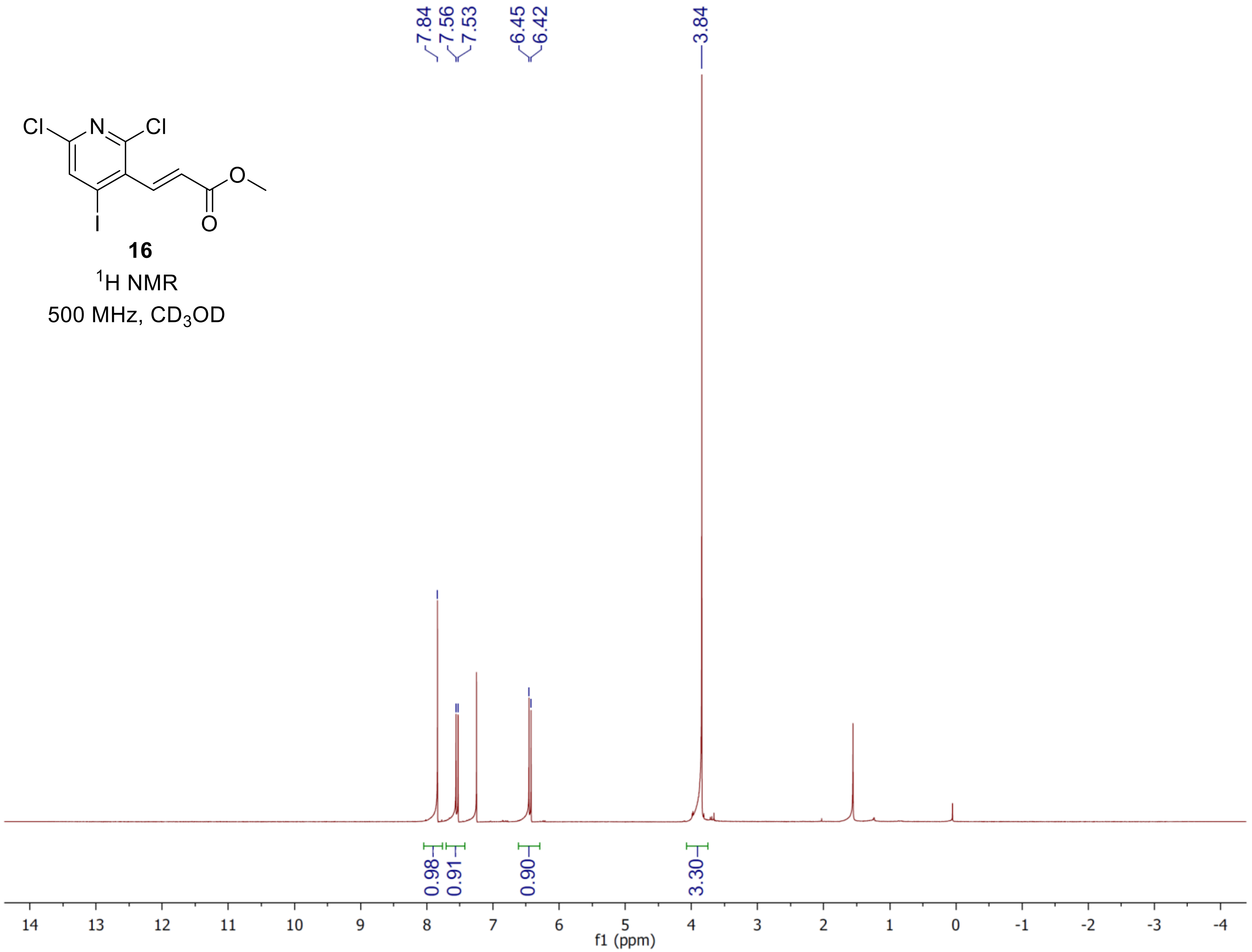

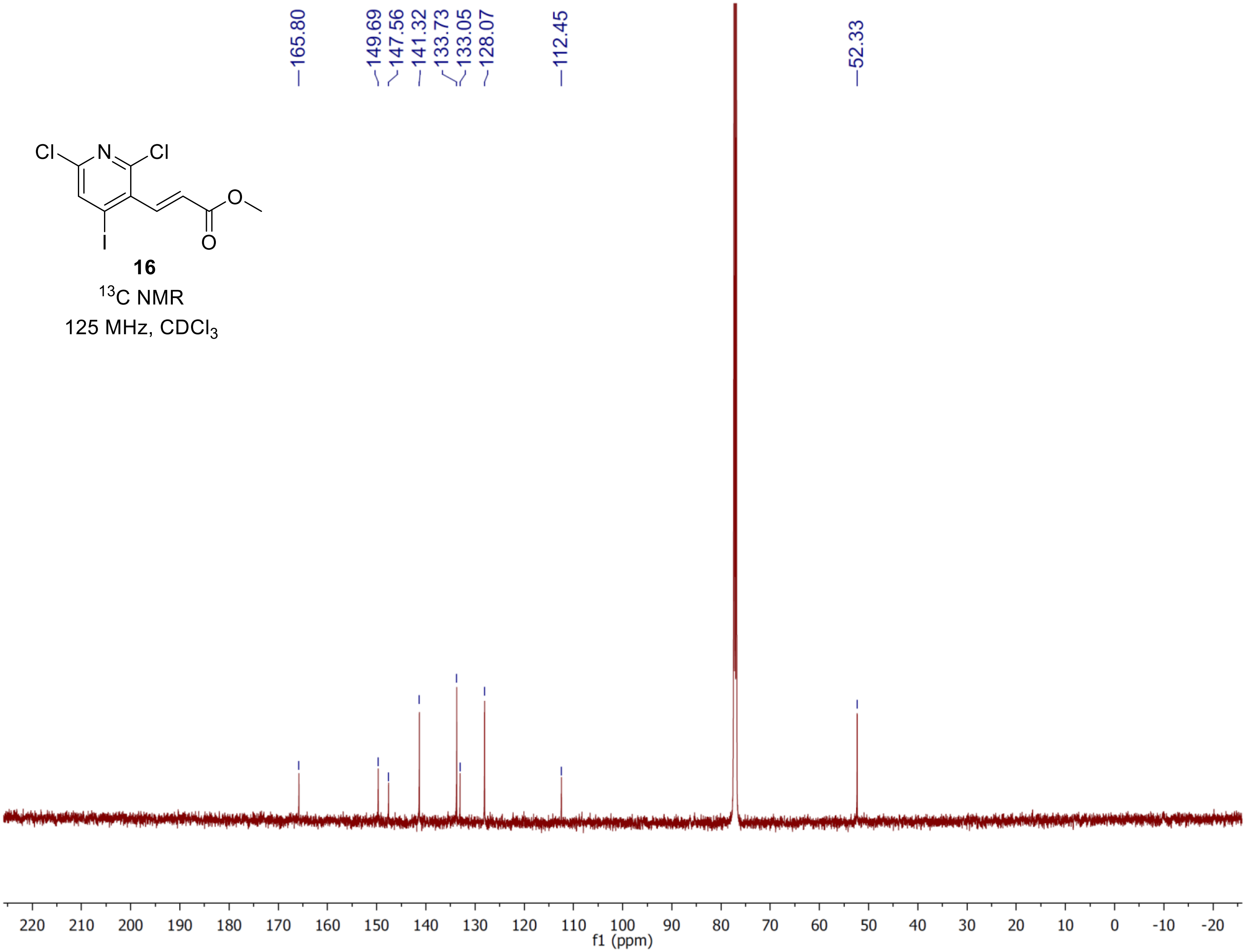


16

$^{13}$C NMR

125 MHz, $CDCl_3$

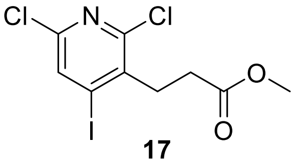

**17**

$^1$H NMR

500 MHz, $CDCl_3$

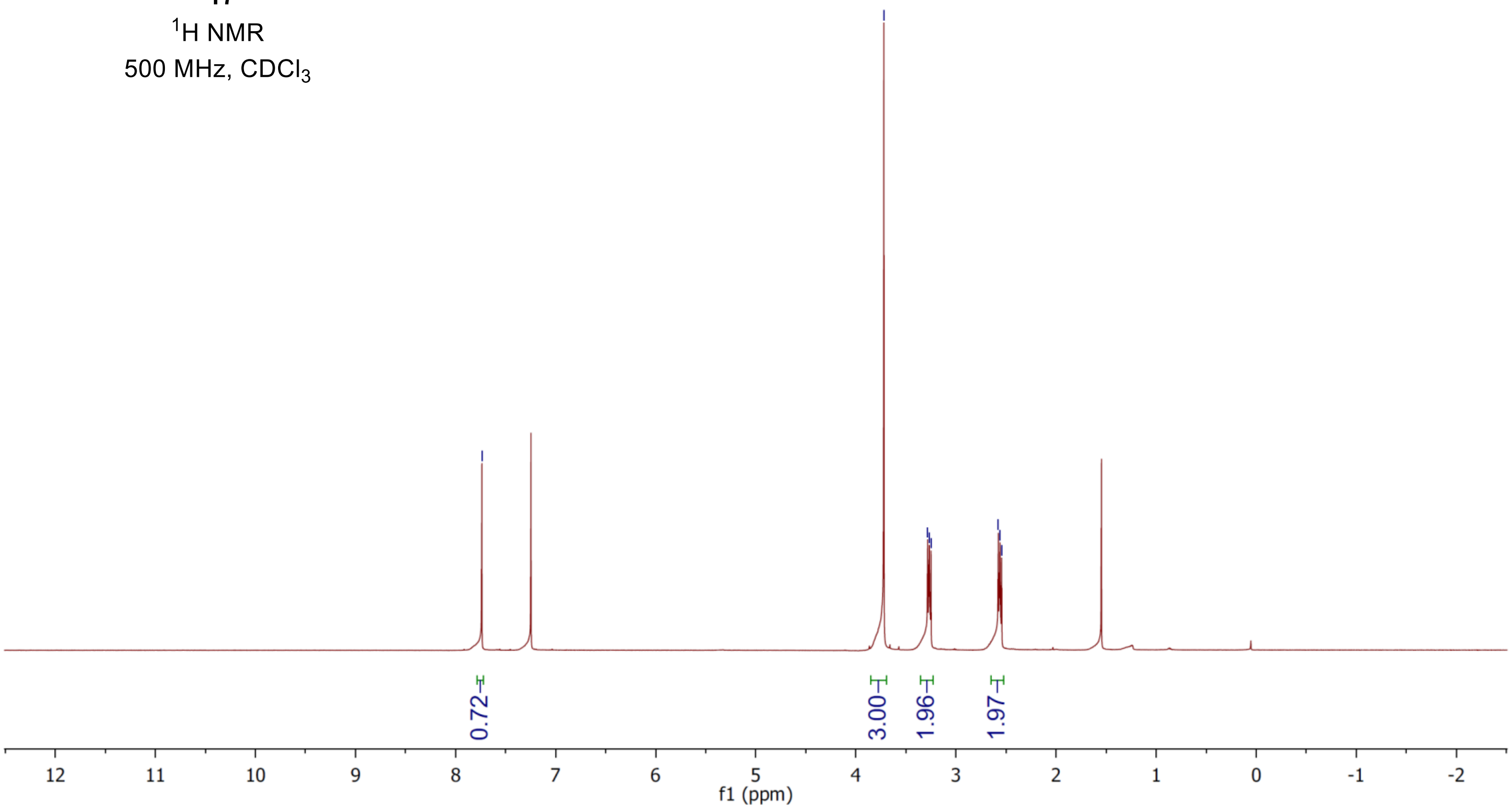

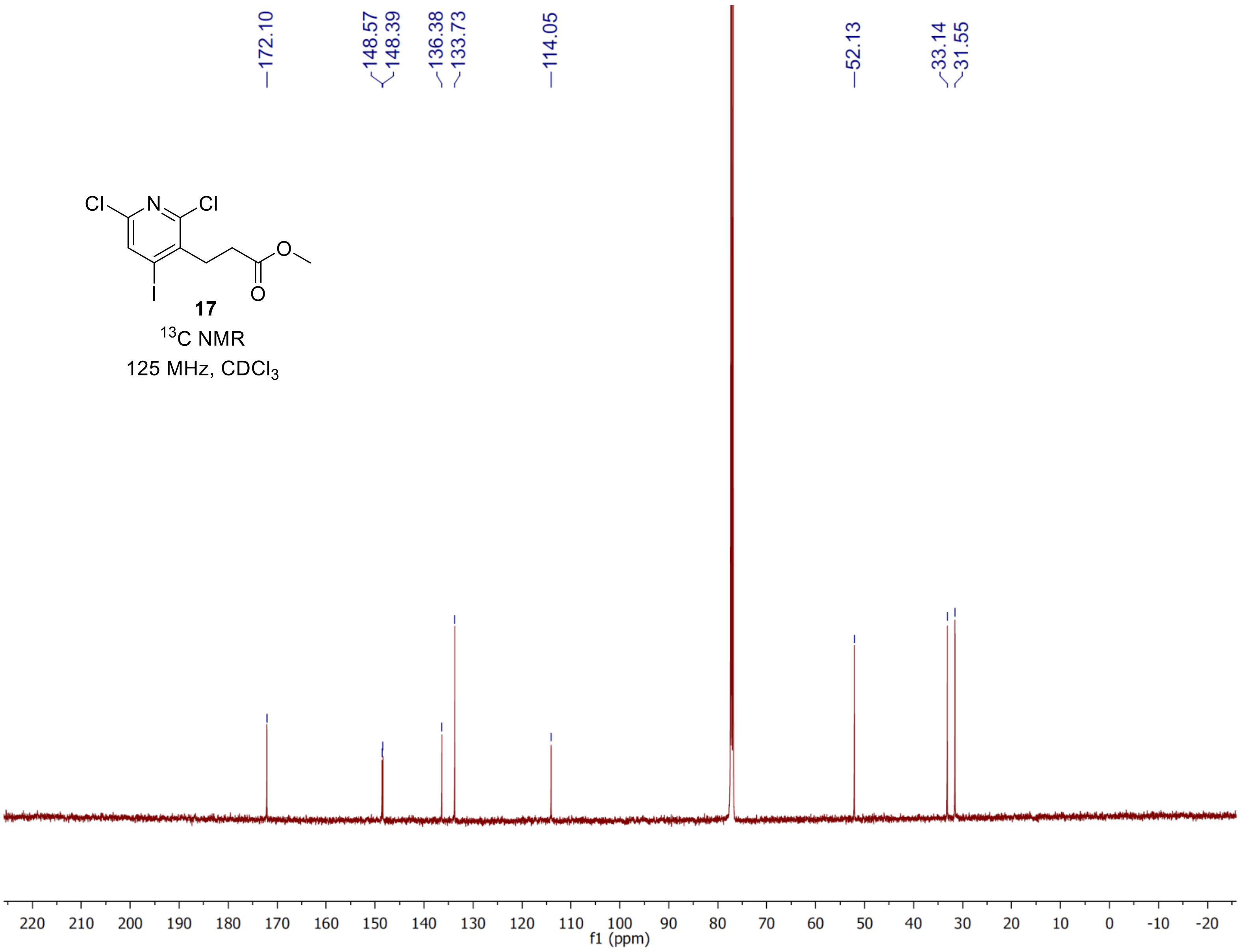
172.10
148.57
148.39
136.38
133.73
114.05
52.13
33.14
31.55
Cl
N
Cl
O
I
O
17
$^{13}$C NMR
125 MHz, $CDCl_3$
220 210 200 190 180 170 160 150 140 130 120 110 100 90 80 70 60 50 40 30 20 10 0 -10 -20
f1 (ppm)

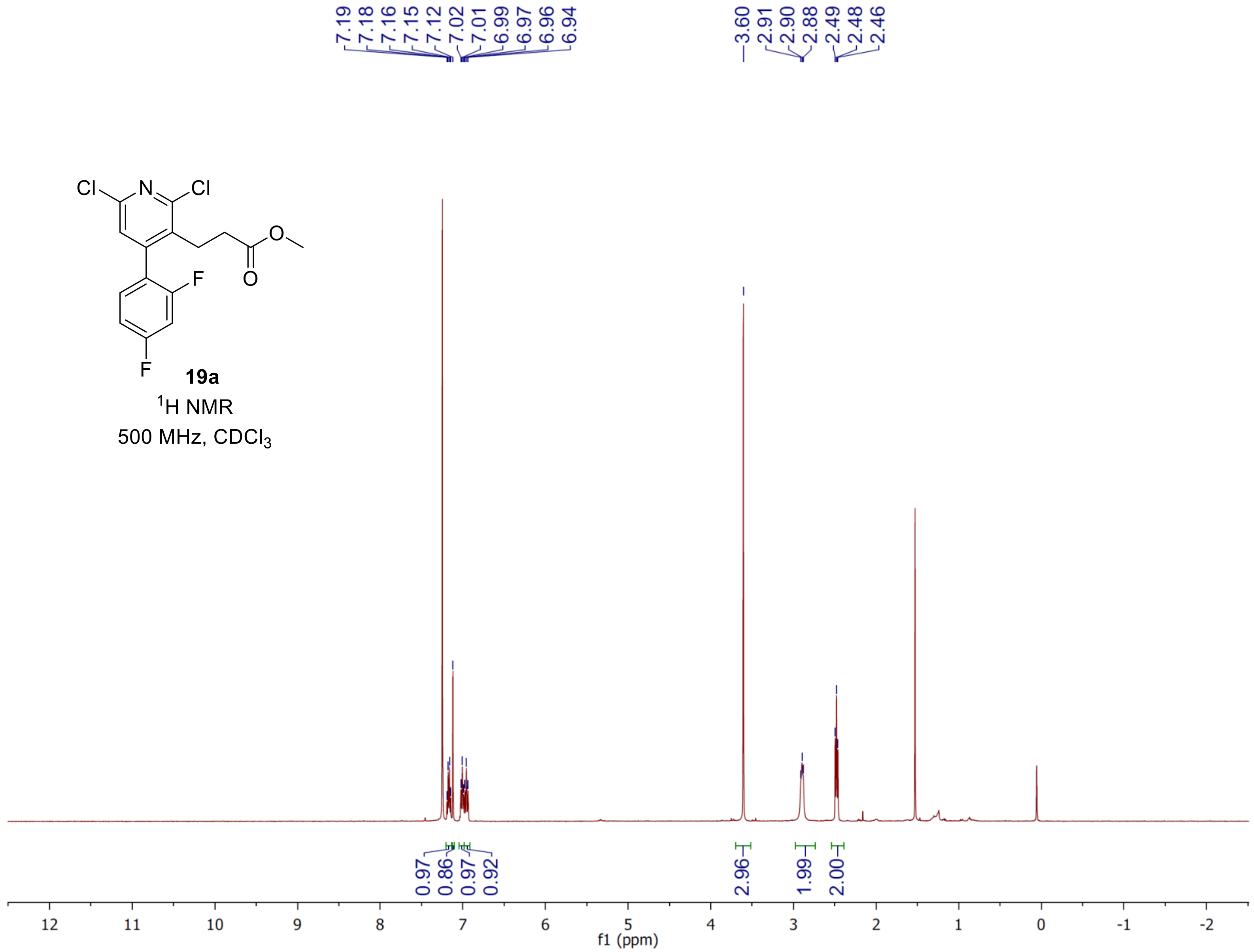

7.19
7.18
7.16
7.15
7.12
7.02
7.01
6.99
6.97
6.96
6.94
3.60
2.91
2.90
2.88
2.49
2.48
2.46
Cl
N
Cl
O
O
F
F
19a
1H NMR
500 MHz, CDCl3
0.97
0.86
0.97
0.92
2.96
1.99
2.00
12
11
10
9
8
7
6
5
4
3
2
1
0
-1
-2
f1 (ppm)

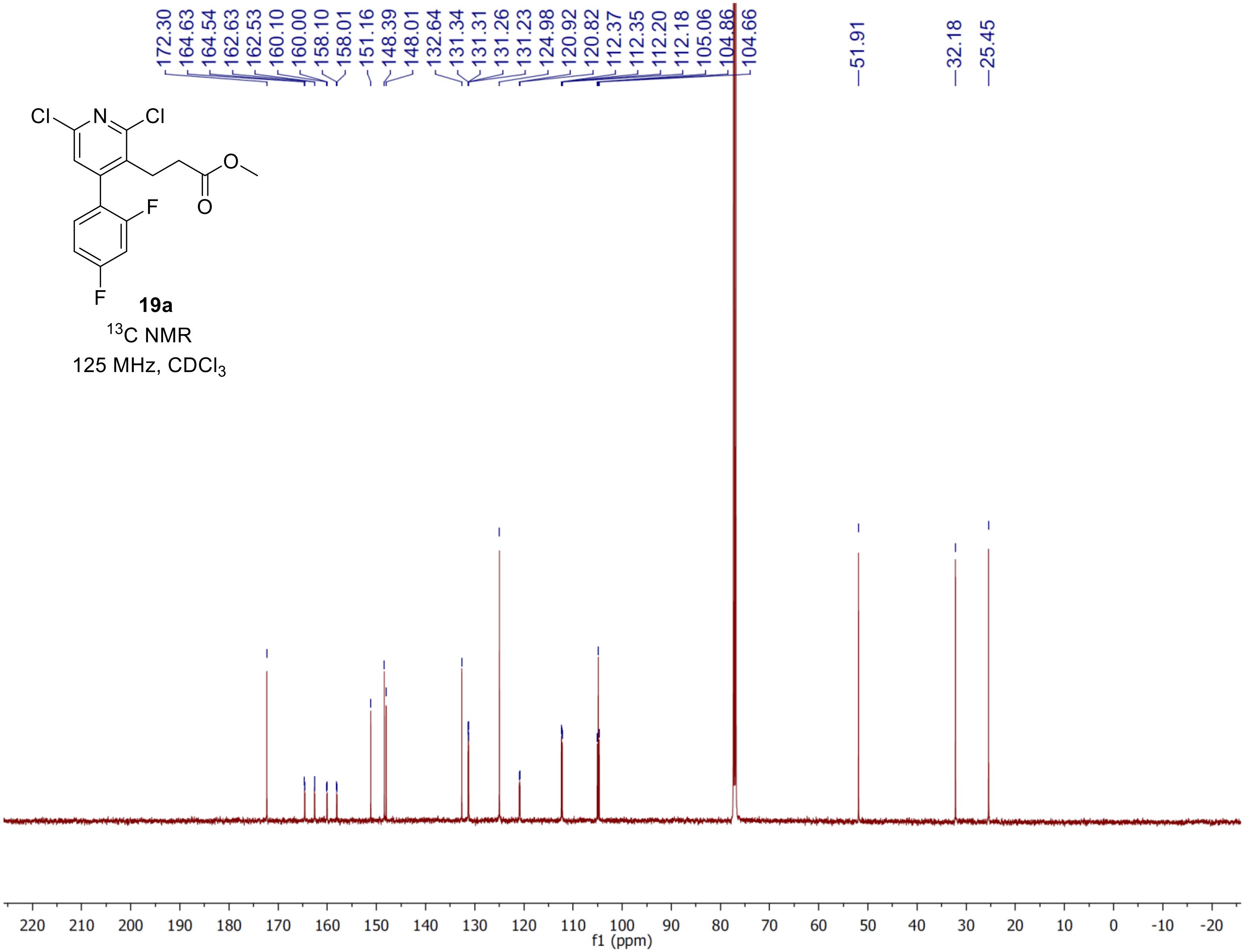
19a
$^{13}$C NMR
125 MHz, $CDCl_3$
172.30
164.63
164.54
162.63
162.53
160.10
160.00
158.10
158.01
151.16
148.39
148.01
132.64
131.34
131.31
131.26
131.23
124.98
120.92
120.82
112.37
112.35
112.20
112.18
105.06
104.86
104.66
51.91
32.18
25.45
f1 (ppm)

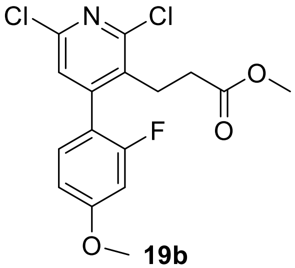


**19b**

$^{1}$H NMR

500 MHz, $CDCl_3$

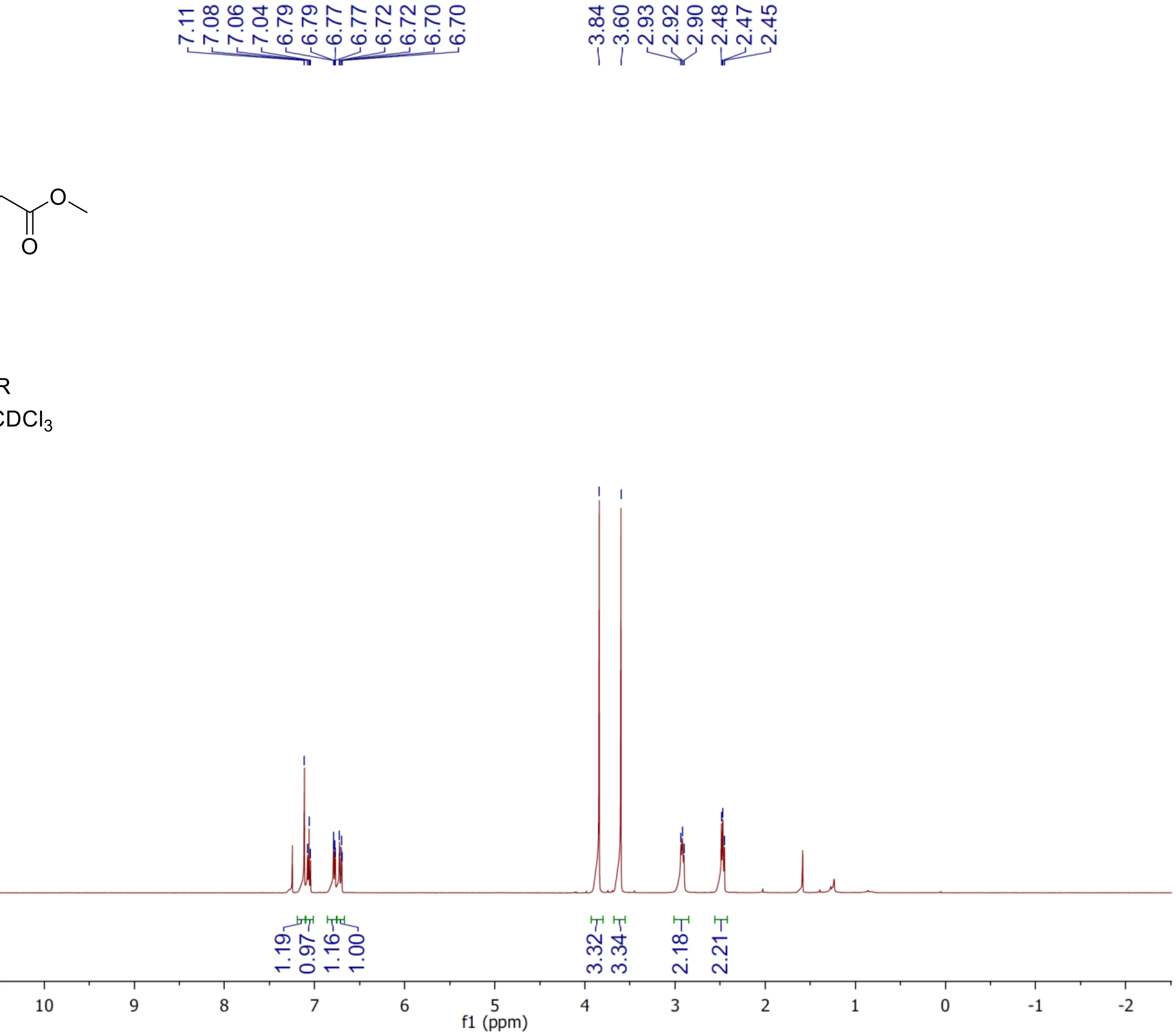

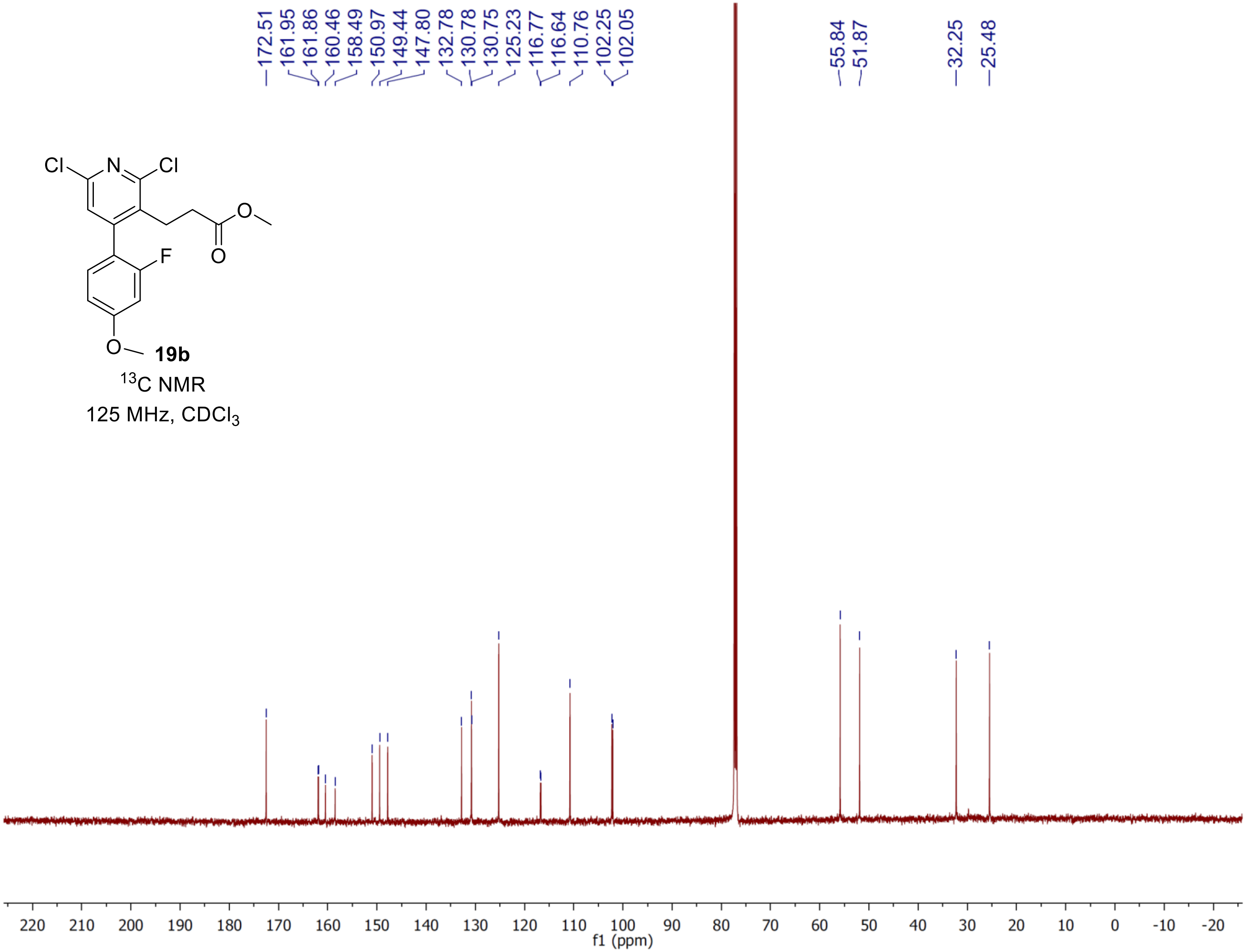

172.51
161.95
161.86
160.46
158.49
150.97
149.44
147.80
132.78
130.78
130.75
125.23
116.77
116.64
110.76
102.25
102.05
55.84
51.87
32.25
25.48
Cl
N
Cl
O
O
F
O
19b
$^{13}$C NMR
125 MHz, $CDCl_3$
220 210 200 190 180 170 160 150 140 130 120 110 100 90 80 70 60 50 40 30 20 10 0 -10 -20
f1 (ppm)

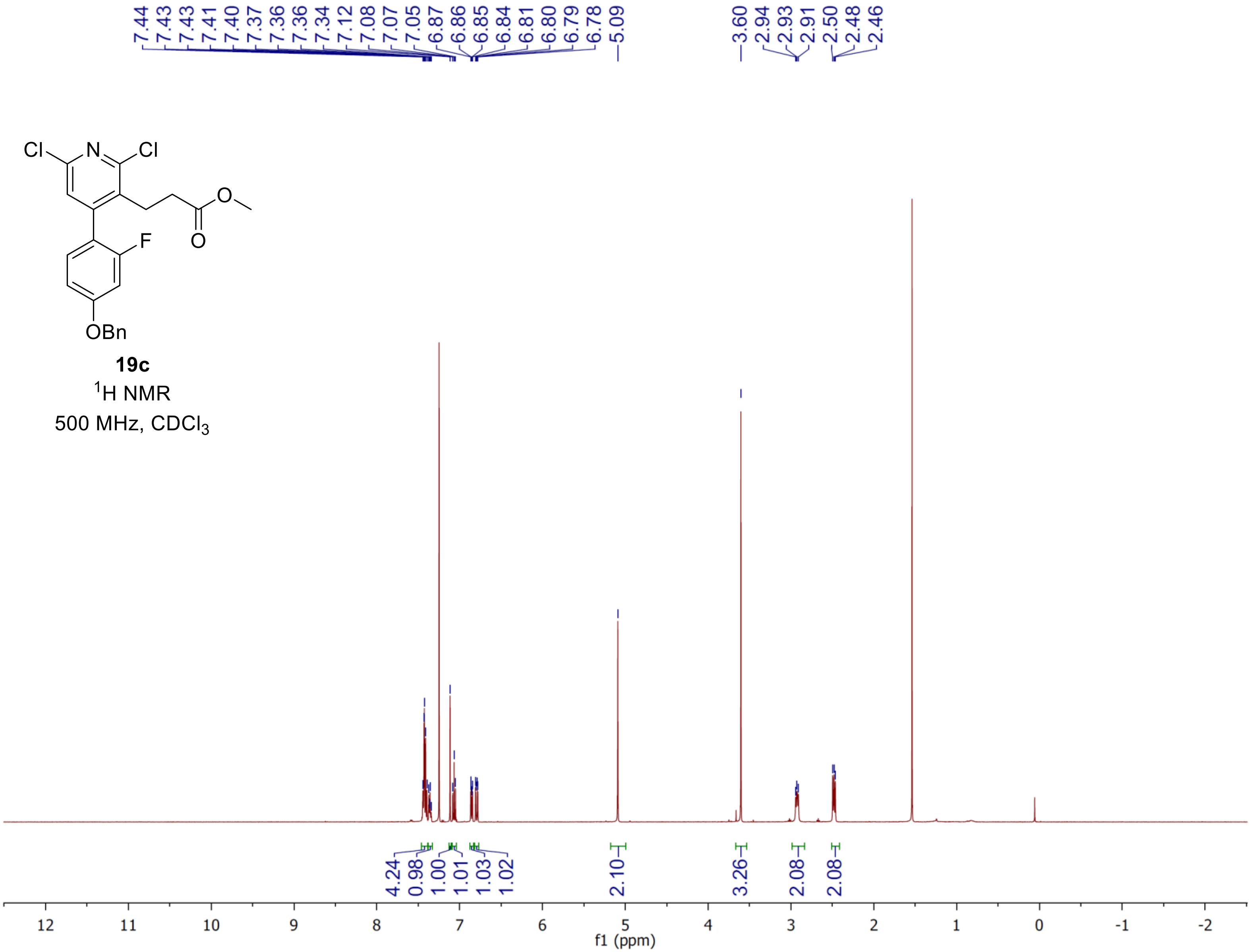
7.44 7.43 7.43 7.41 7.40 7.37 7.36 7.36 7.34 7.12 7.08 7.07 7.05 6.87 6.86 6.85 6.84 6.81 6.80 6.79 6.78 5.09
3.60 2.94 2.93 2.91 2.50 2.48 2.46
Cl N Cl
O
O
F
OBn
19c
1H NMR
500 MHz, CDCl3
4.24 0.98 1.00 1.01 1.03 1.02
2.10
3.26
2.08
2.08
12 11 10 9 8 7 6 5 4 3 2 1 0 -1 -2
f1 (ppm)

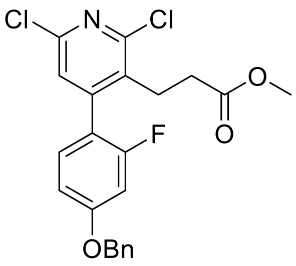

**19c**

$^{13}C$ NMR

125 MHz, $CDCl_3$

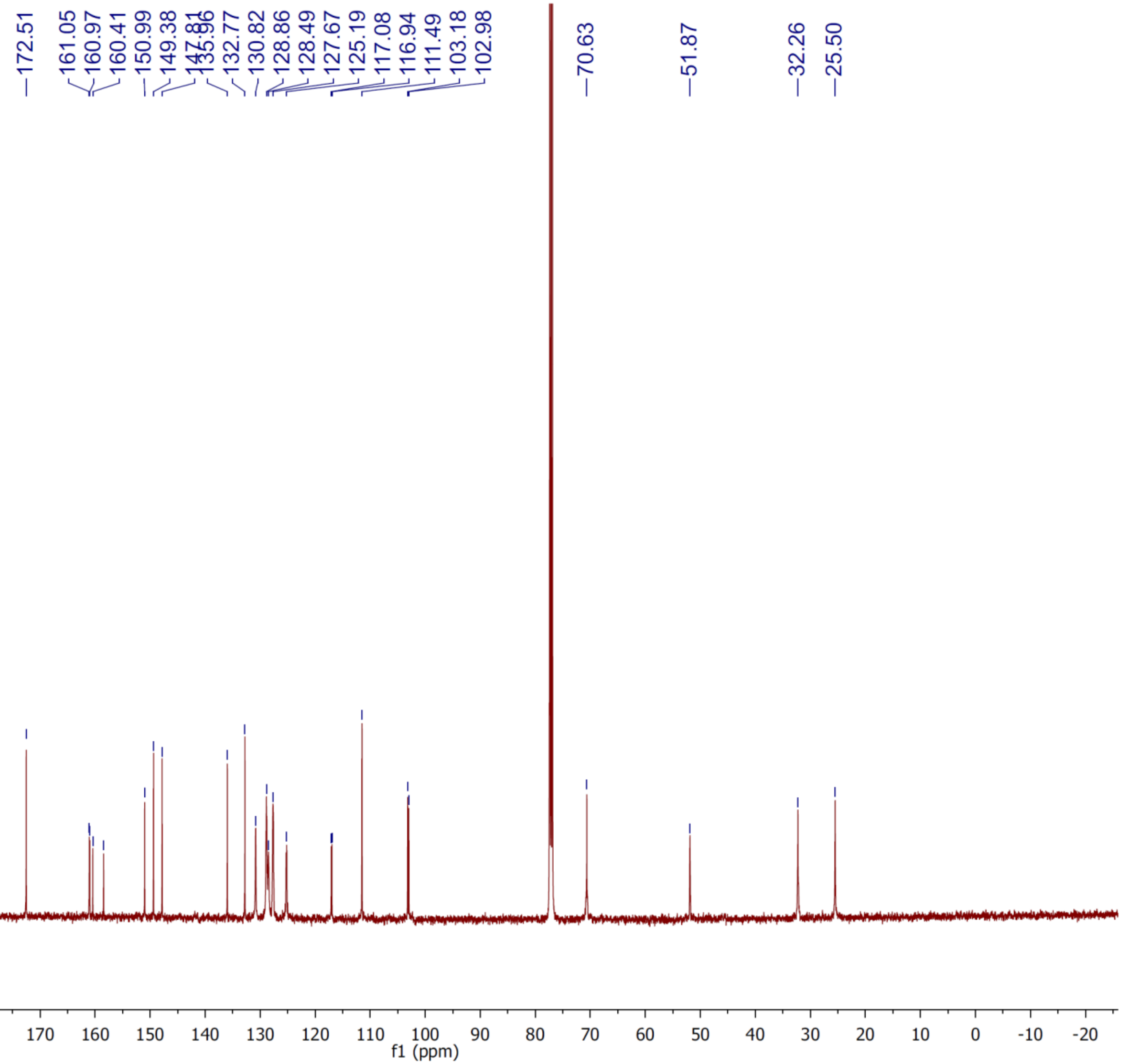

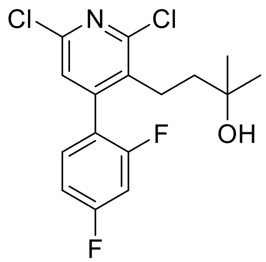


**20a**

$^{1}H$ NMR

500 MHz, $CDCl_3$

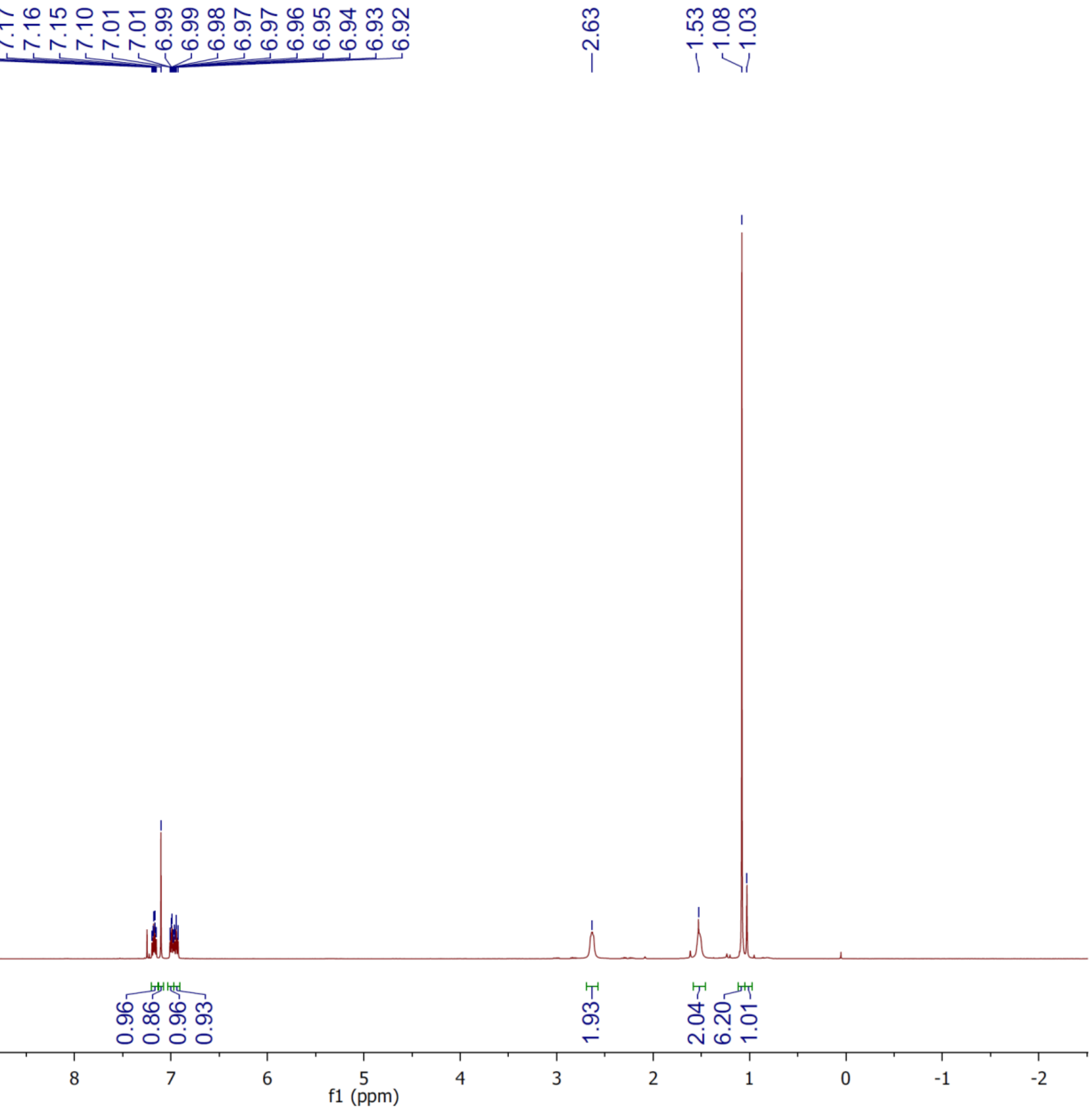

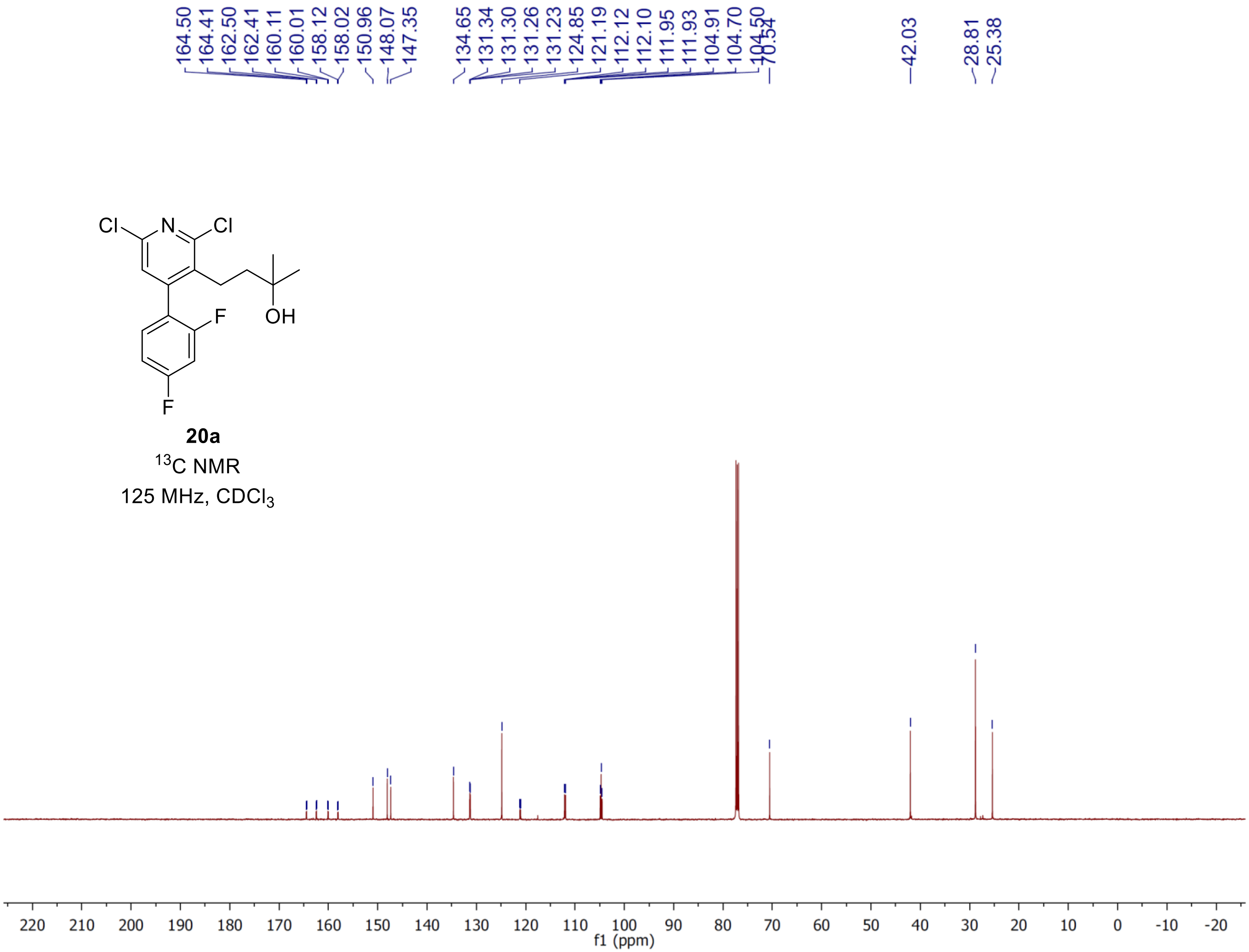
164.50
164.41
162.50
162.41
160.11
160.01
158.12
158.02
150.96
148.07
147.35
134.65
131.34
131.30
131.26
131.23
124.85
121.19
112.12
112.10
111.95
111.93
104.91
104.70
104.50
70.54
42.03
28.81
25.38
Cl
N
Cl
F
OH
F
20a
$^{13}$C NMR
125 MHz, $CDCl_3$
f1 (ppm)

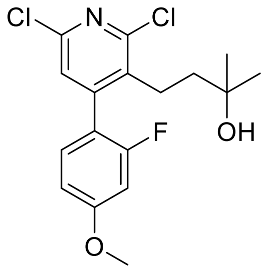


**20b**

$^{1}H$ NMR

500 MHz, $CDCl_3$

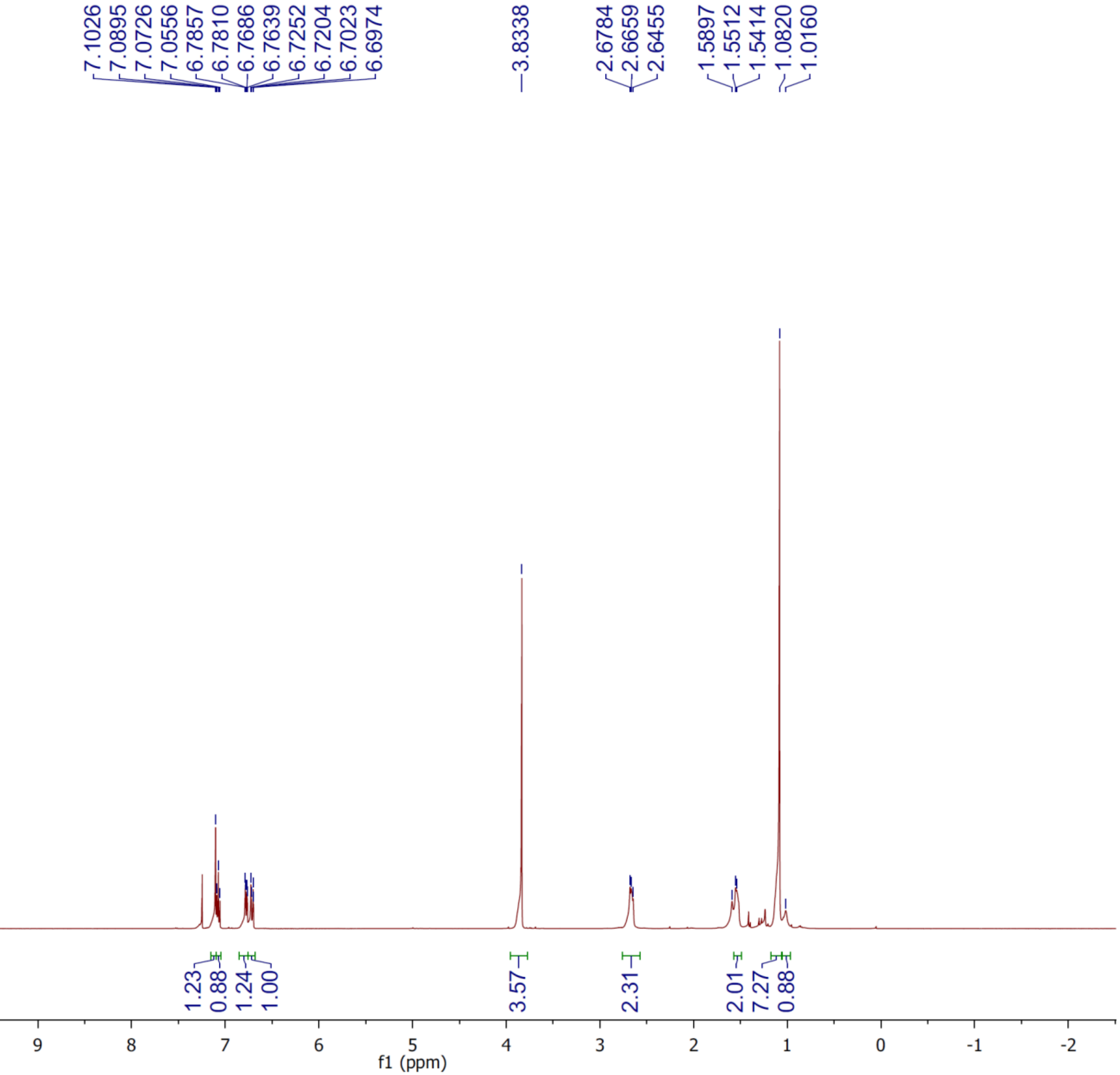

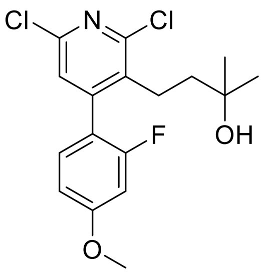


**20b**

$^{13}C$ NMR

125 MHz, $CDCl_3$

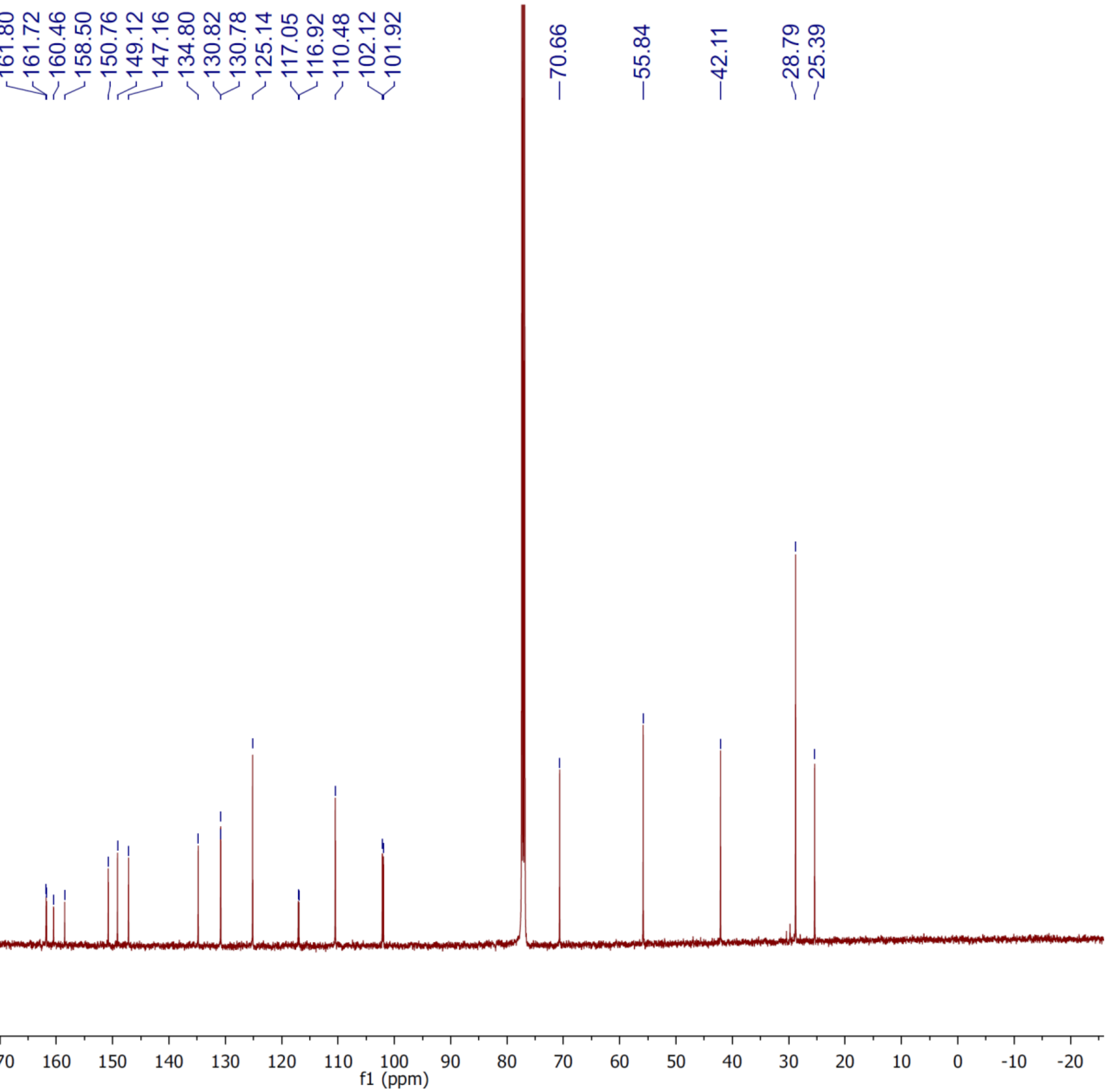

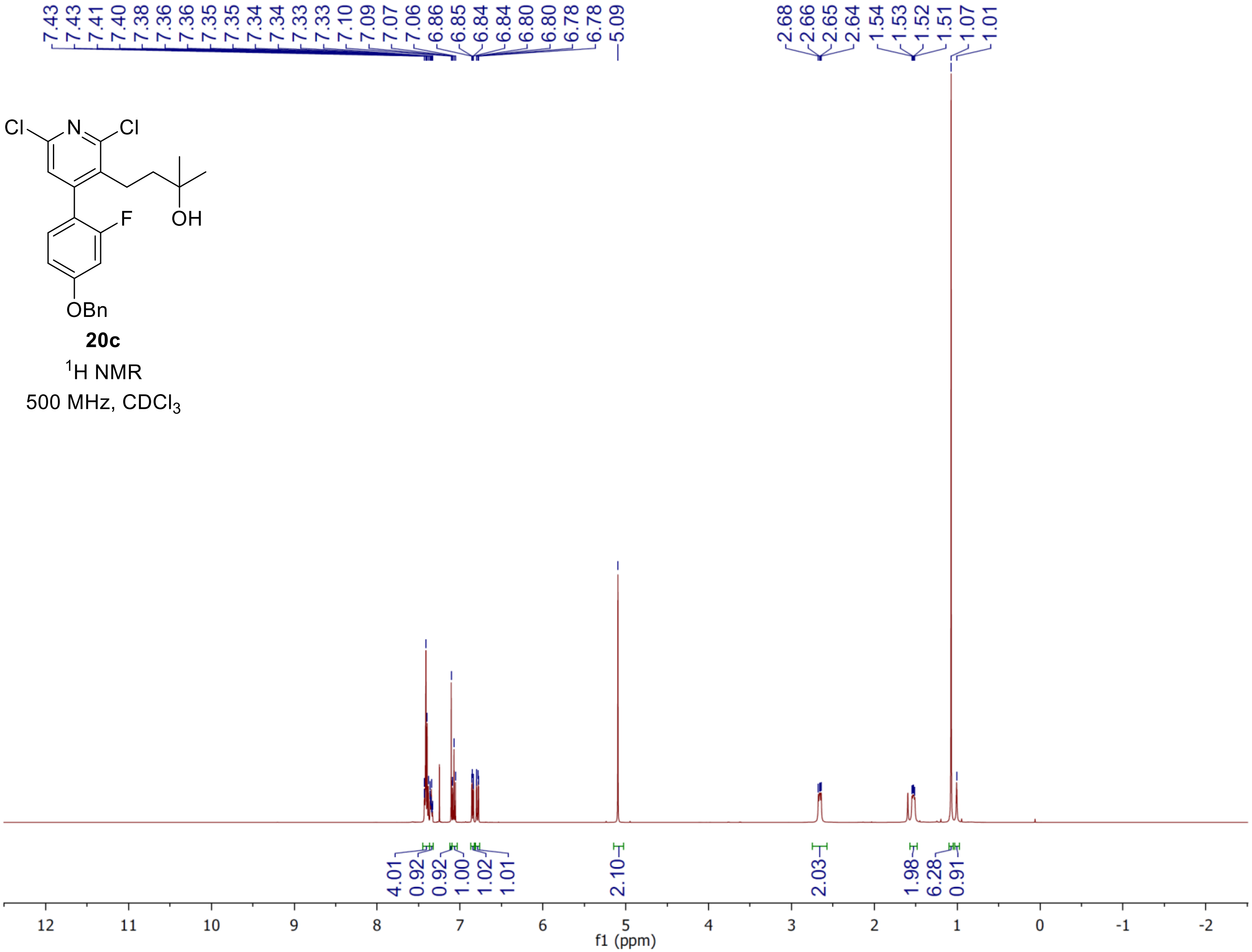
7.43 7.43 7.41 7.40 7.38 7.36 7.36 7.35 7.35 7.34 7.34 7.33 7.33 7.10 7.09 7.07 7.06 6.86 6.85 6.84 6.84 6.80 6.80 6.78 6.78 5.09
2.68 2.66 2.65 2.64 1.54 1.53 1.52 1.51 1.07 1.01
Cl N Cl
F OH
OBn
20c
$^1$H NMR
500 MHz, $CDCl_3$
4.01 0.92 0.92 1.00 1.02 1.01 2.10 2.03 1.98 6.28 0.91
12 11 10 9 8 7 6 5 4 3 2 1 0 -1 -2
f1 (ppm)

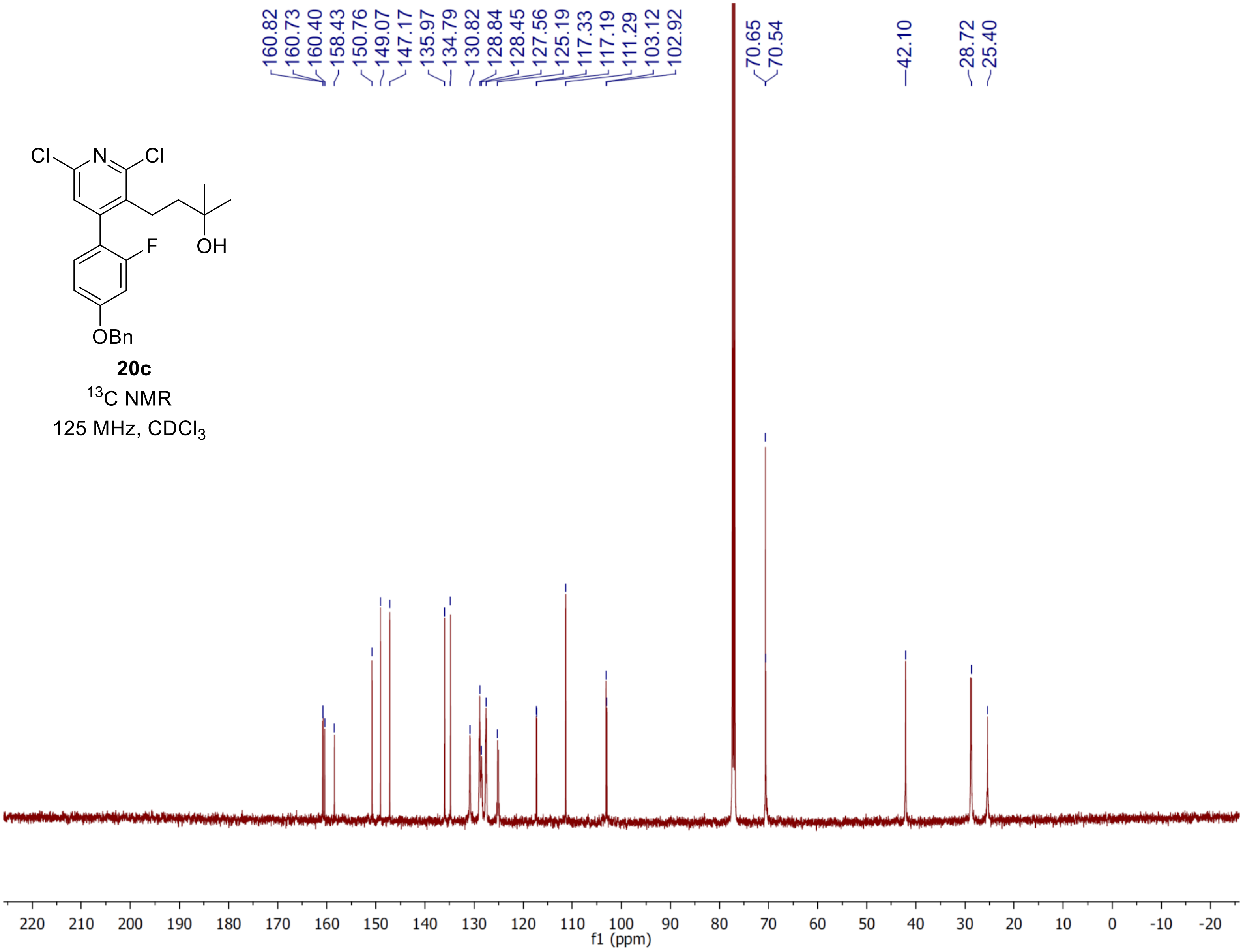


**20c**

$^{13}C$ NMR

125 MHz, $CDCl_3$

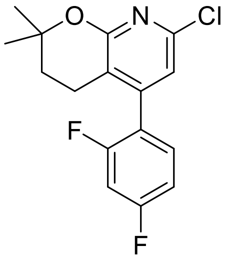

**21a**

$^{1}H$ NMR

500 MHz, $CDCl_3$

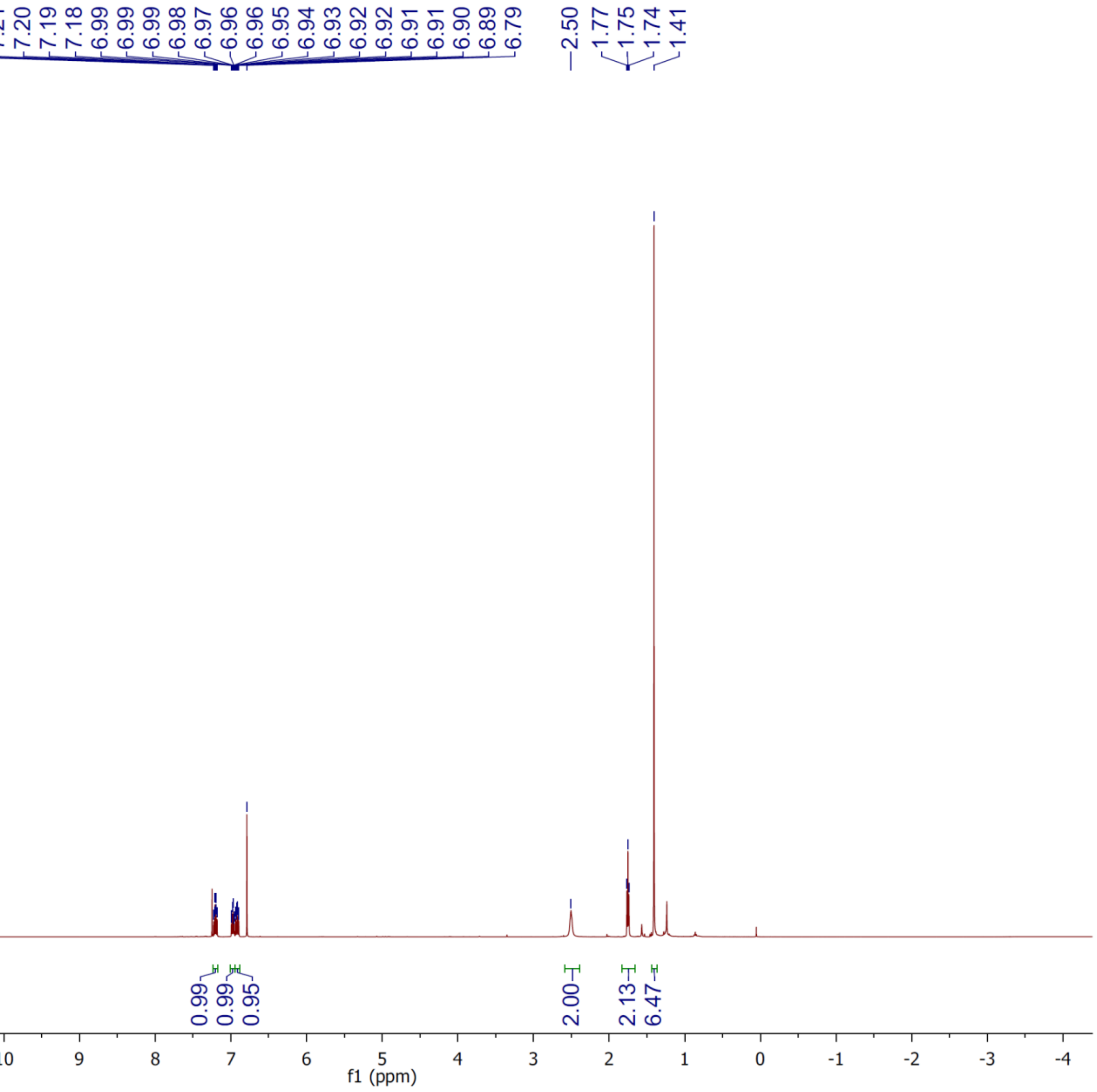

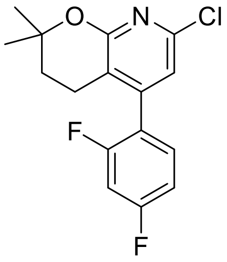


**21a**

$^{13}$C NMR

125 MHz, $CDCl_3$

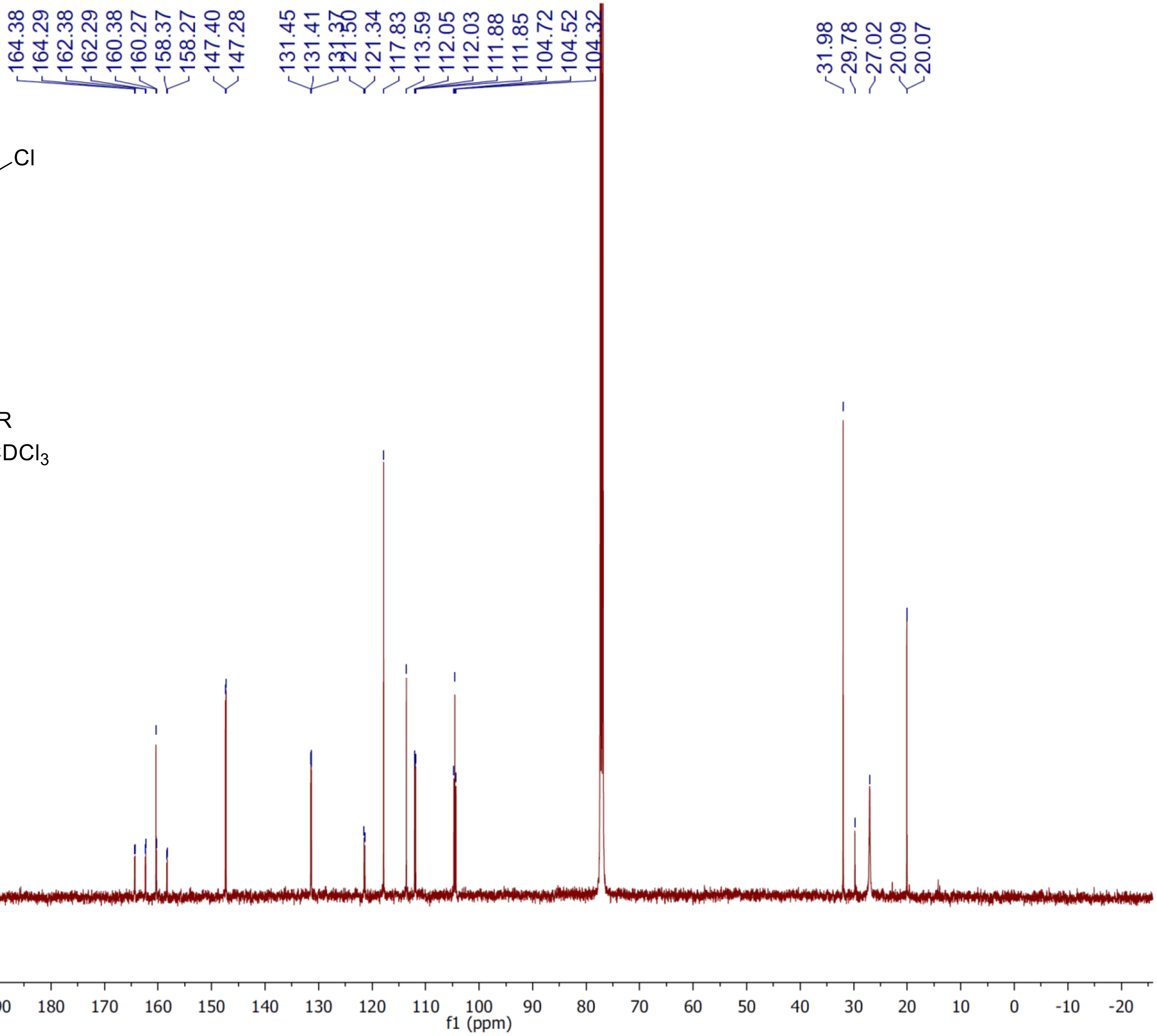

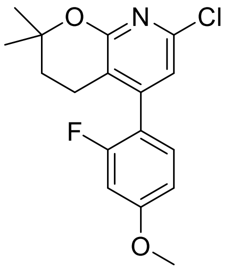

**21b**

$^{1}$H NMR

500 MHz, $CDCl_3$

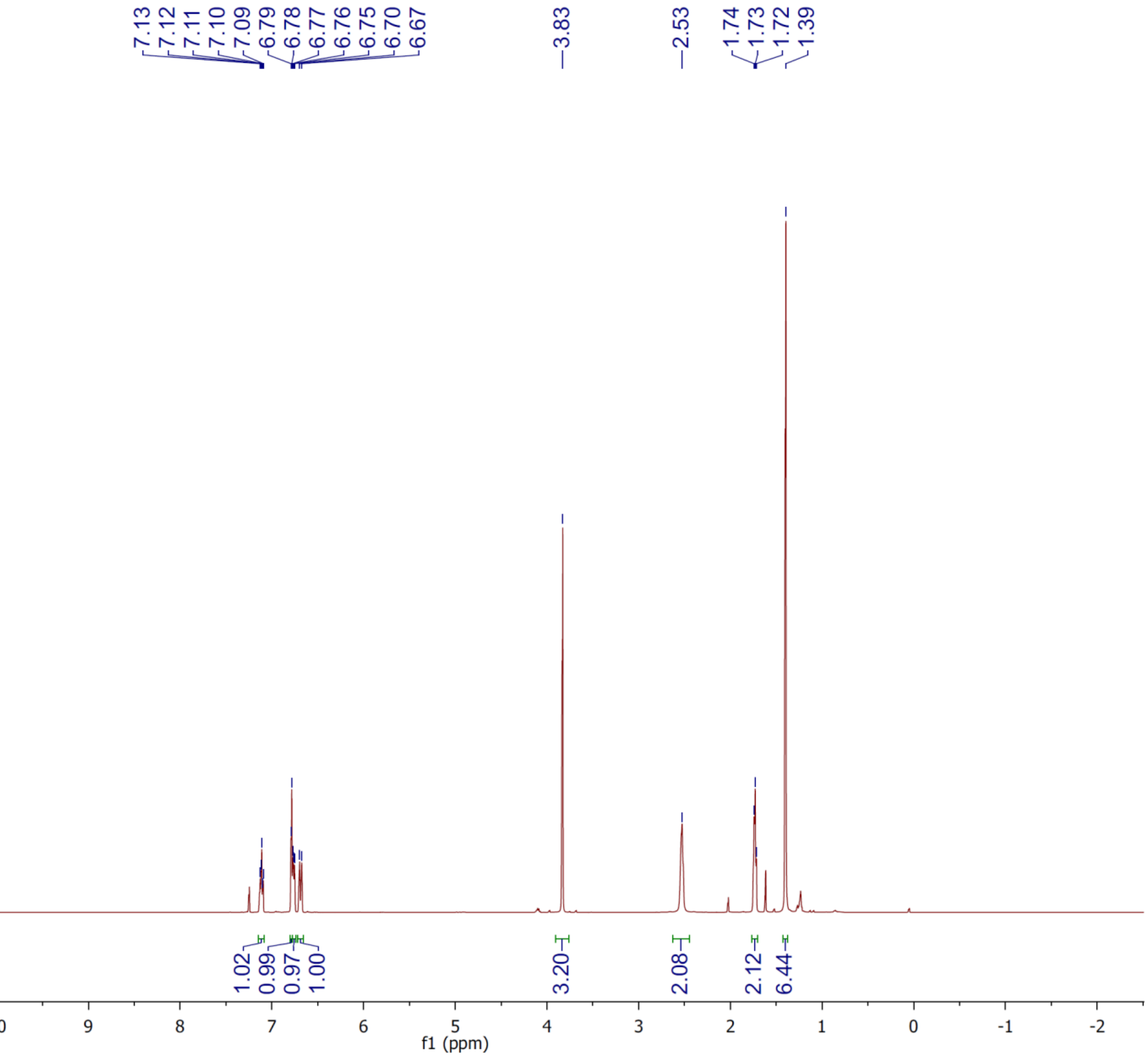

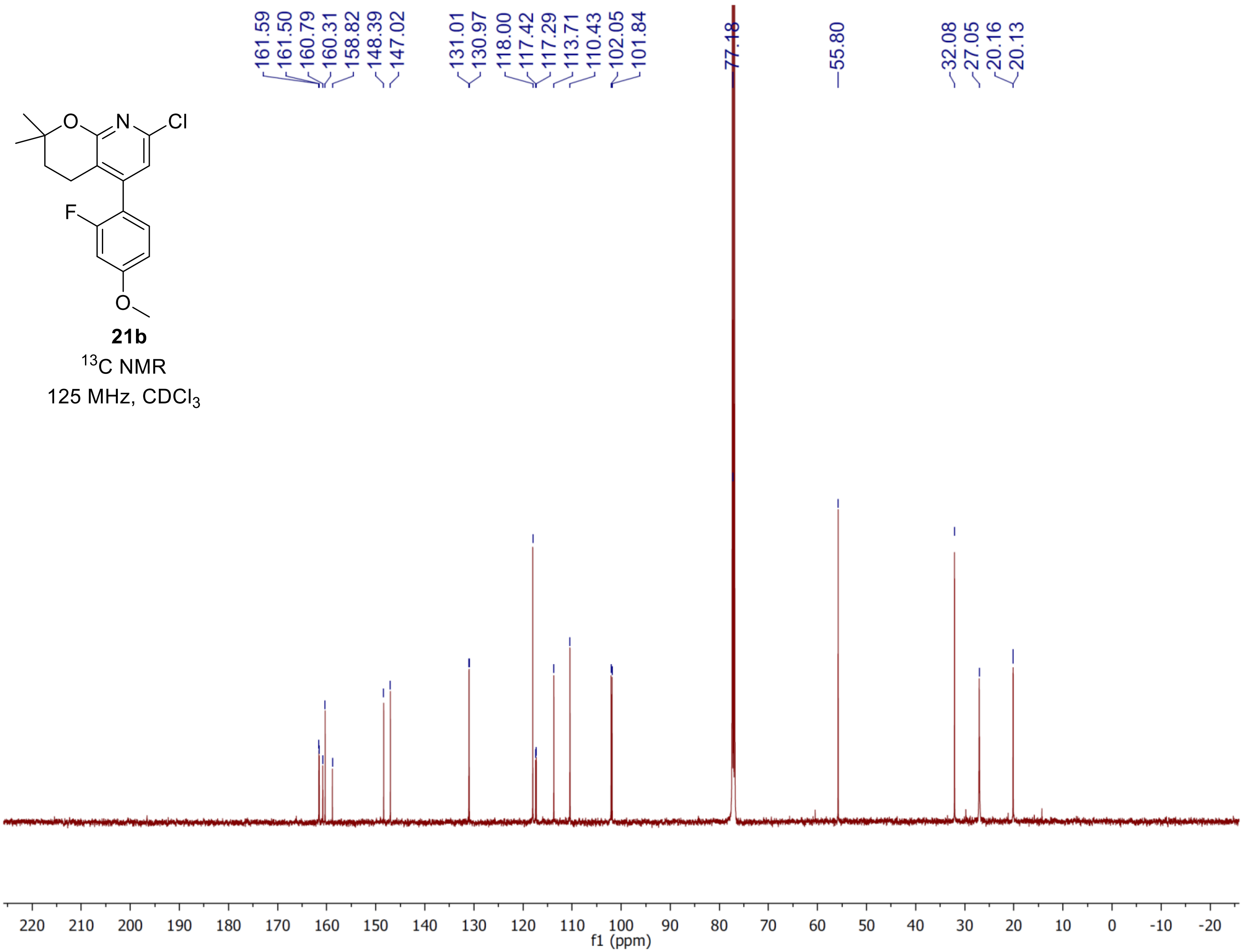
21b
$^{13}$C NMR
125 MHz, CDCl$_3$
161.59
161.50
160.79
160.31
158.82
148.39
147.02
131.01
130.97
118.00
117.42
117.29
113.71
110.43
102.05
101.84
77.18
55.80
32.08
27.05
20.16
20.13
f1 (ppm)

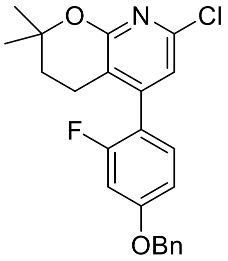

**21c**

$^{1}$H NMR

500 MHz, $CDCl_3$

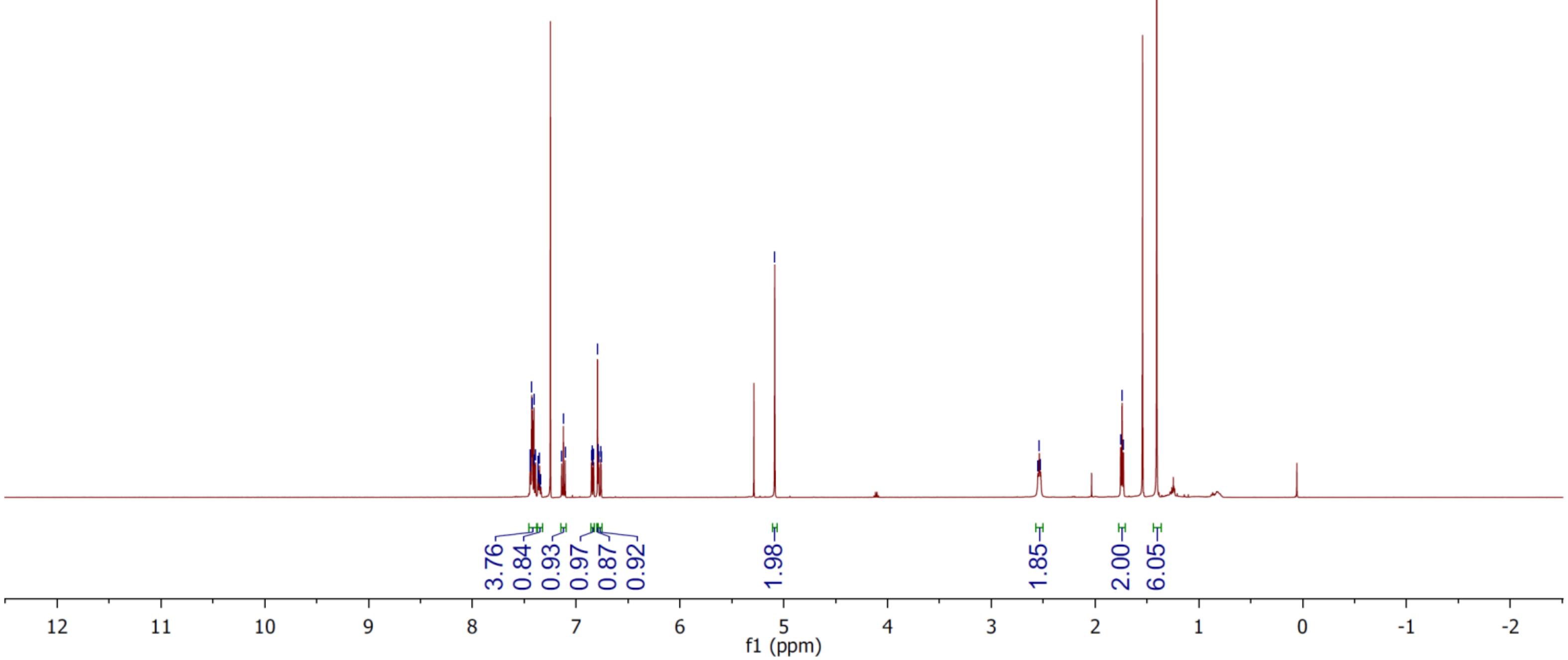

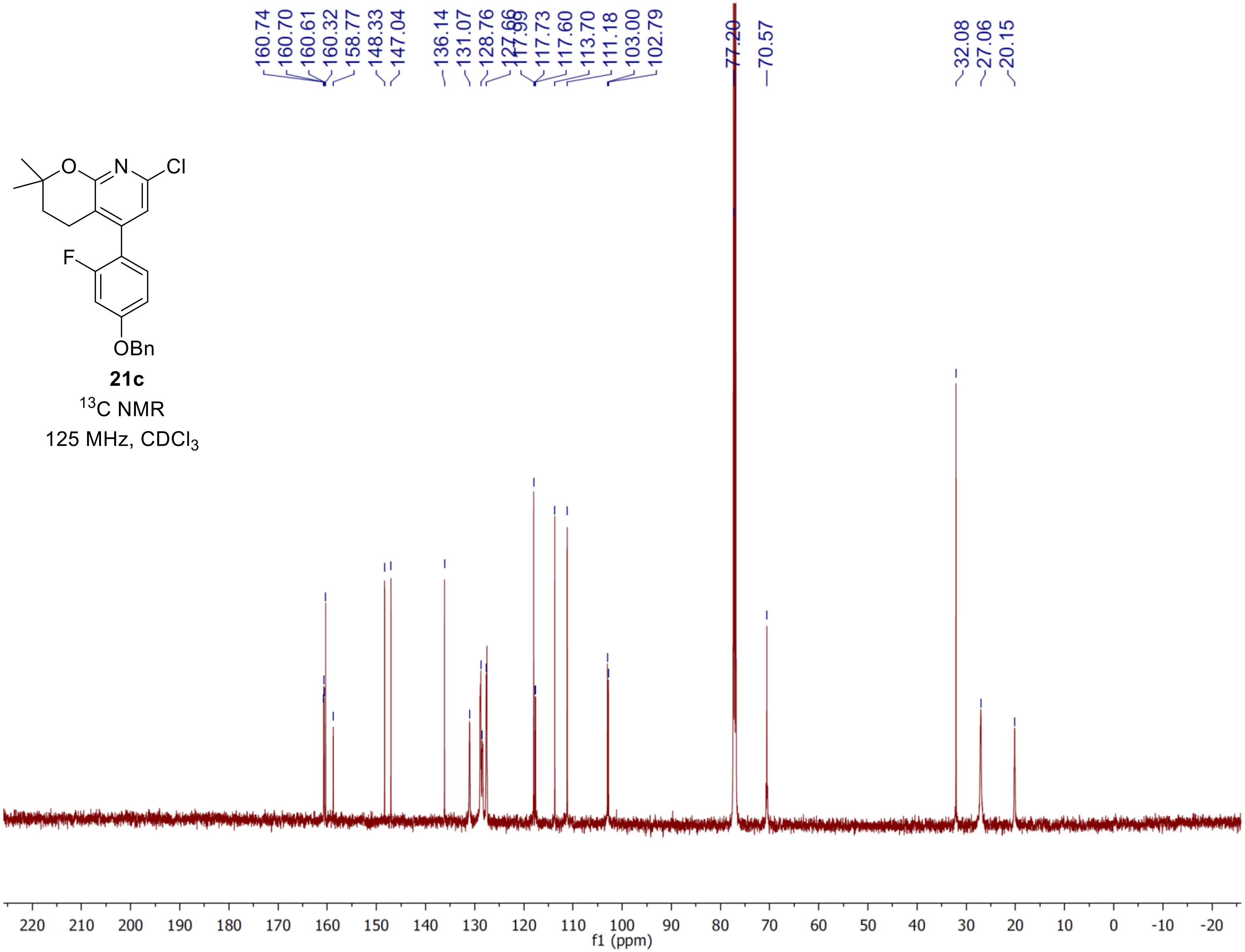
160.74
160.70
160.61
160.32
158.77
148.33
147.04
136.14
131.07
128.76
117.99
117.73
117.60
113.70
111.18
103.00
102.79
77.20
70.57
32.08
27.06
20.15
O
N
Cl
F
OBn
21c
$^{13}$C NMR
125 MHz, $CDCl_3$
f1 (ppm)

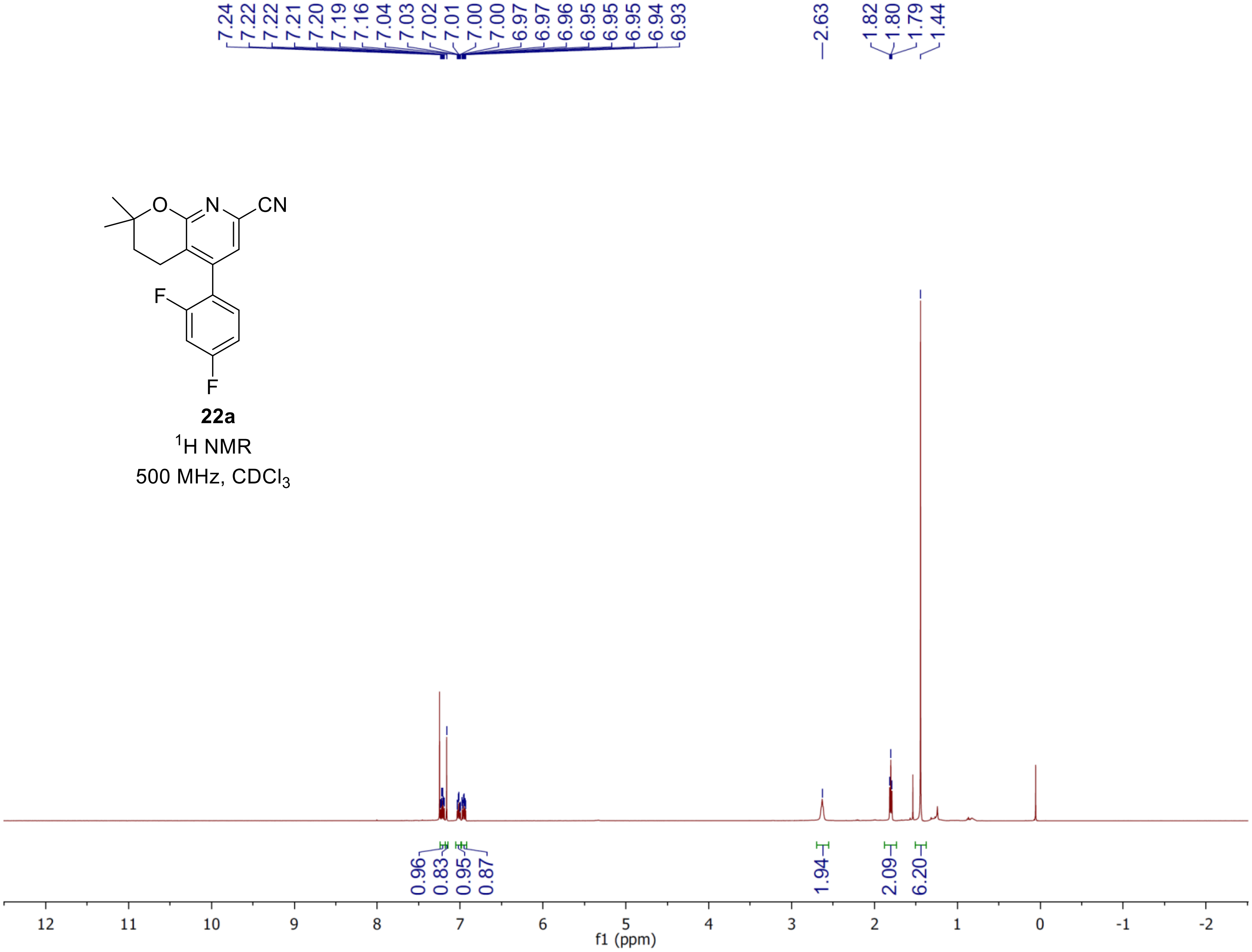


**22a**

$^{1}H$ NMR

500 MHz, $CDCl_3$

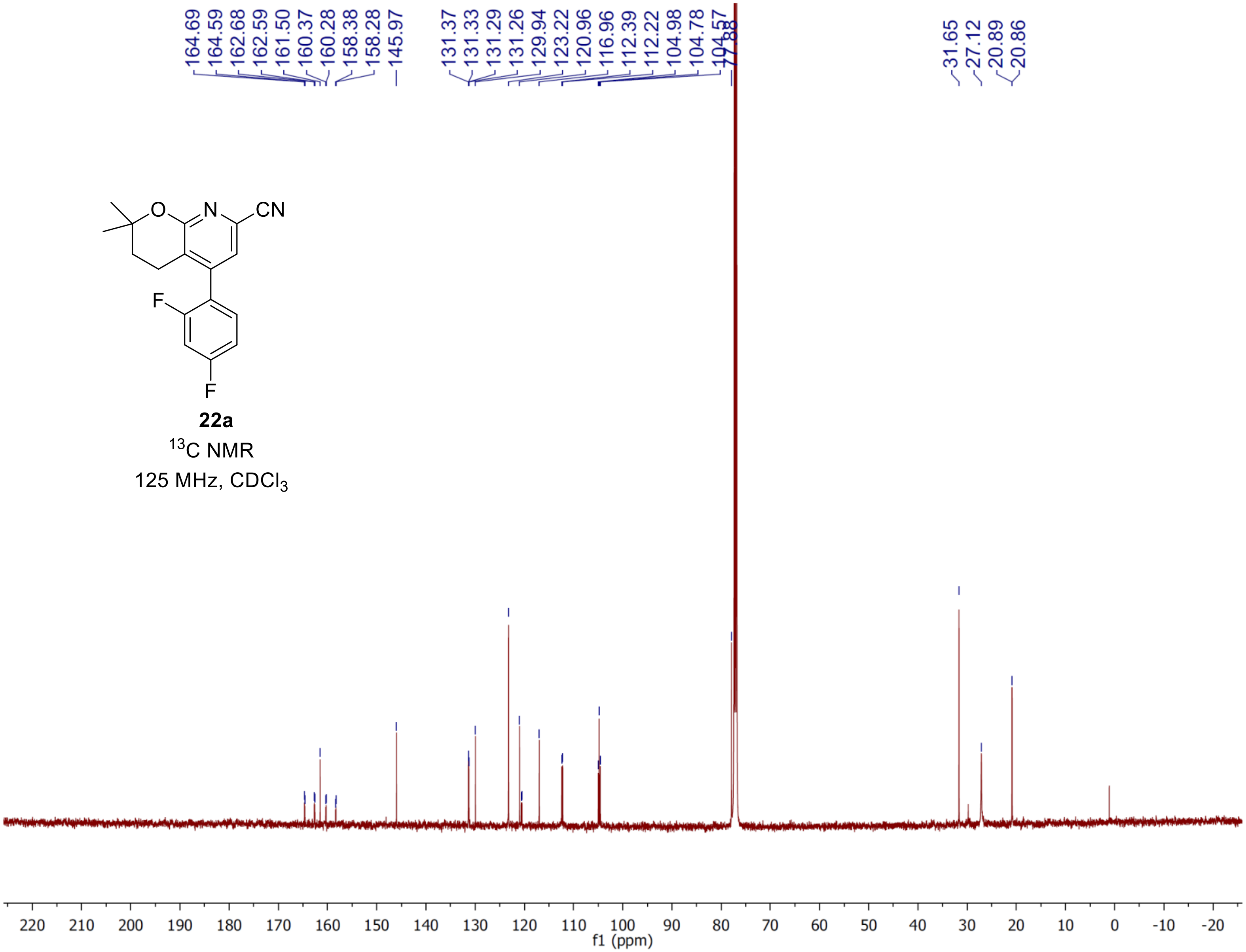


**22a**

$^{13}C$ NMR

125 MHz, $CDCl_3$

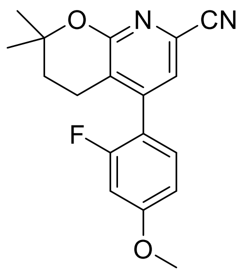

**22b**

$^{1}$H NMR

500 MHz, $CDCl_3$

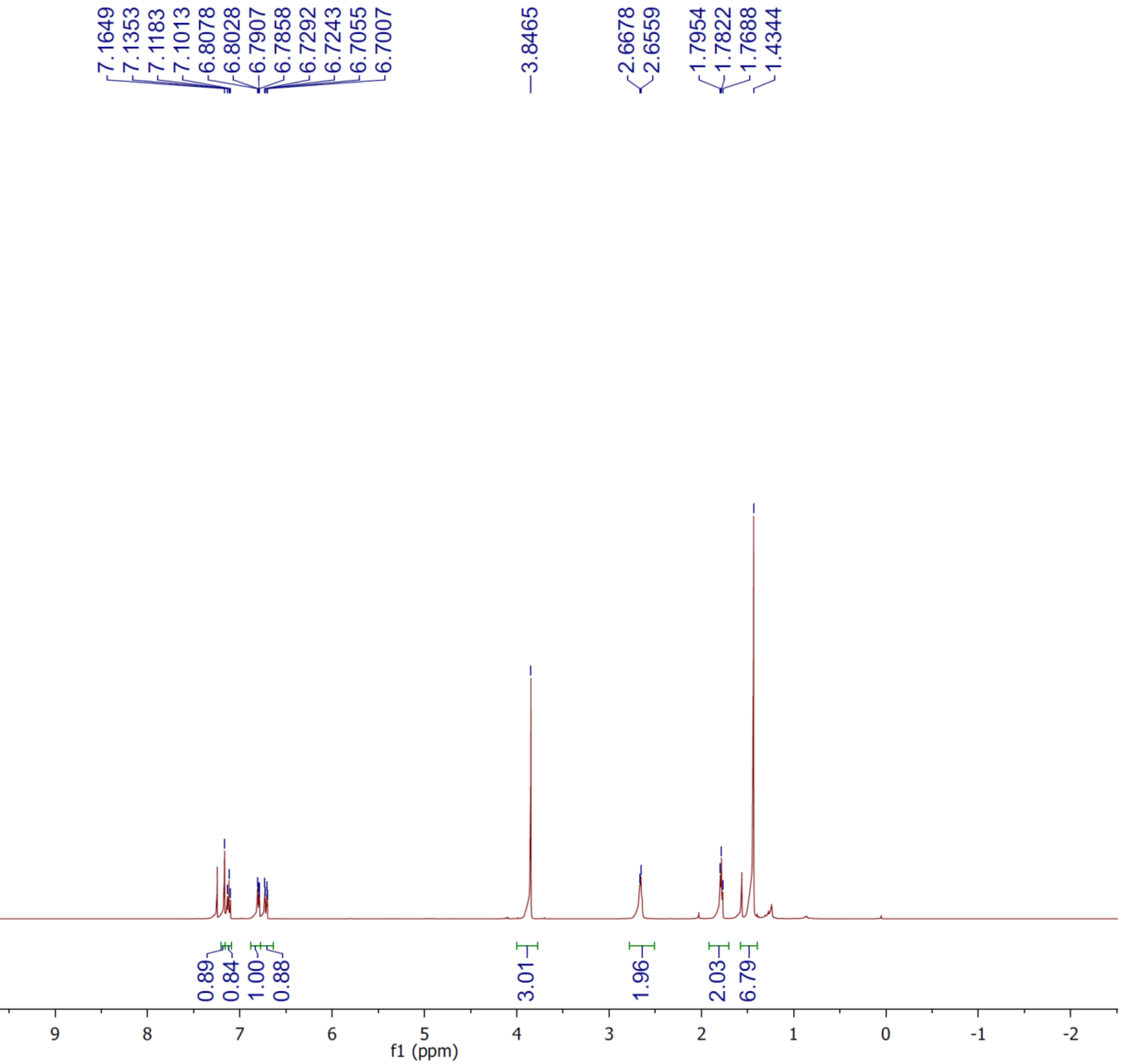

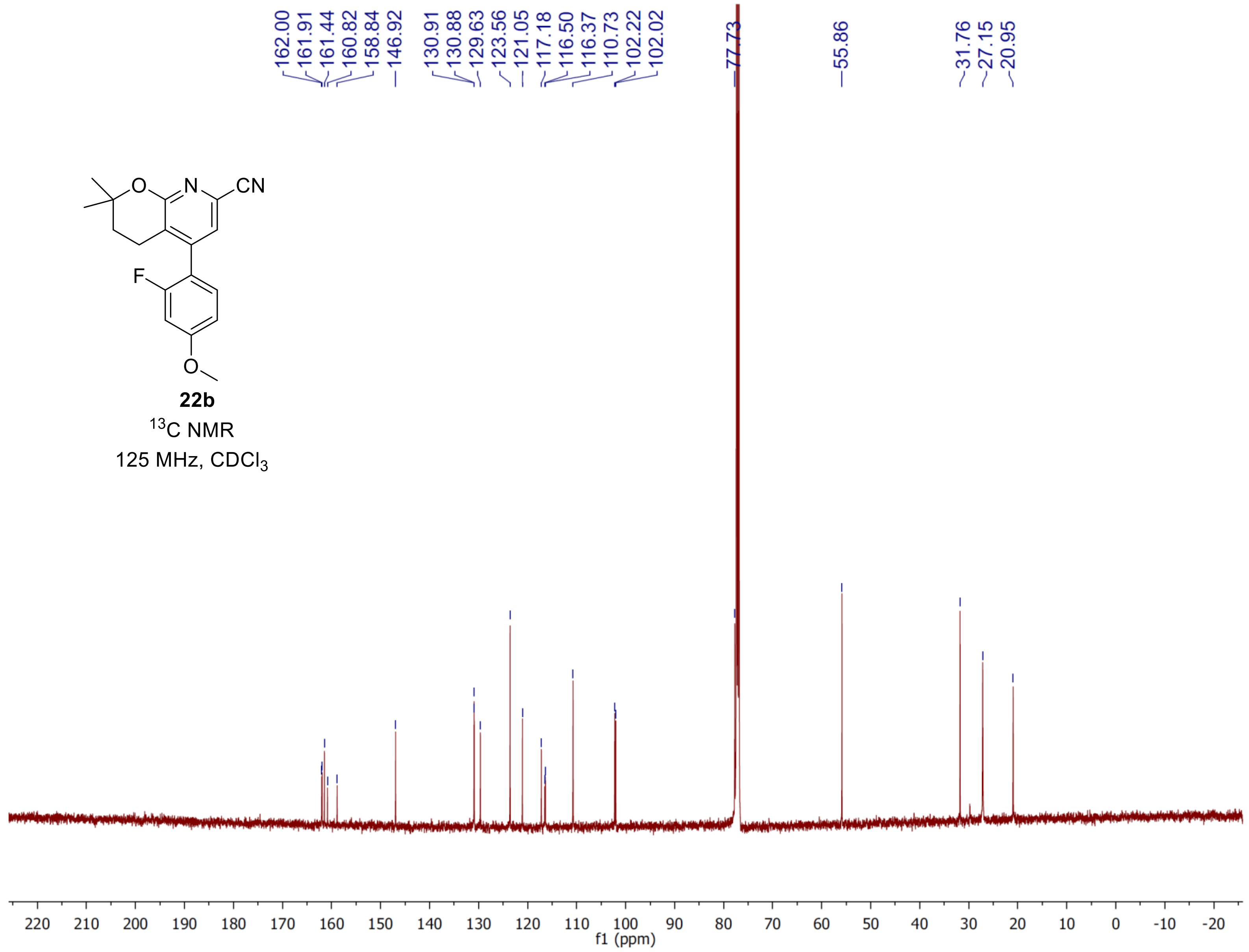
162.00
161.91
161.44
160.82
158.84
146.92
130.91
130.88
129.63
123.56
121.05
117.18
116.50
116.37
110.73
102.22
102.02
77.73
55.86
31.76
27.15
20.95
CN
N
O
F
O
22b
$^{13}$C NMR
125 MHz, $CDCl_3$
220 210 200 190 180 170 160 150 140 130 120 110 100 90 80 70 60 50 40 30 20 10 0 -10 -20
f1 (ppm)

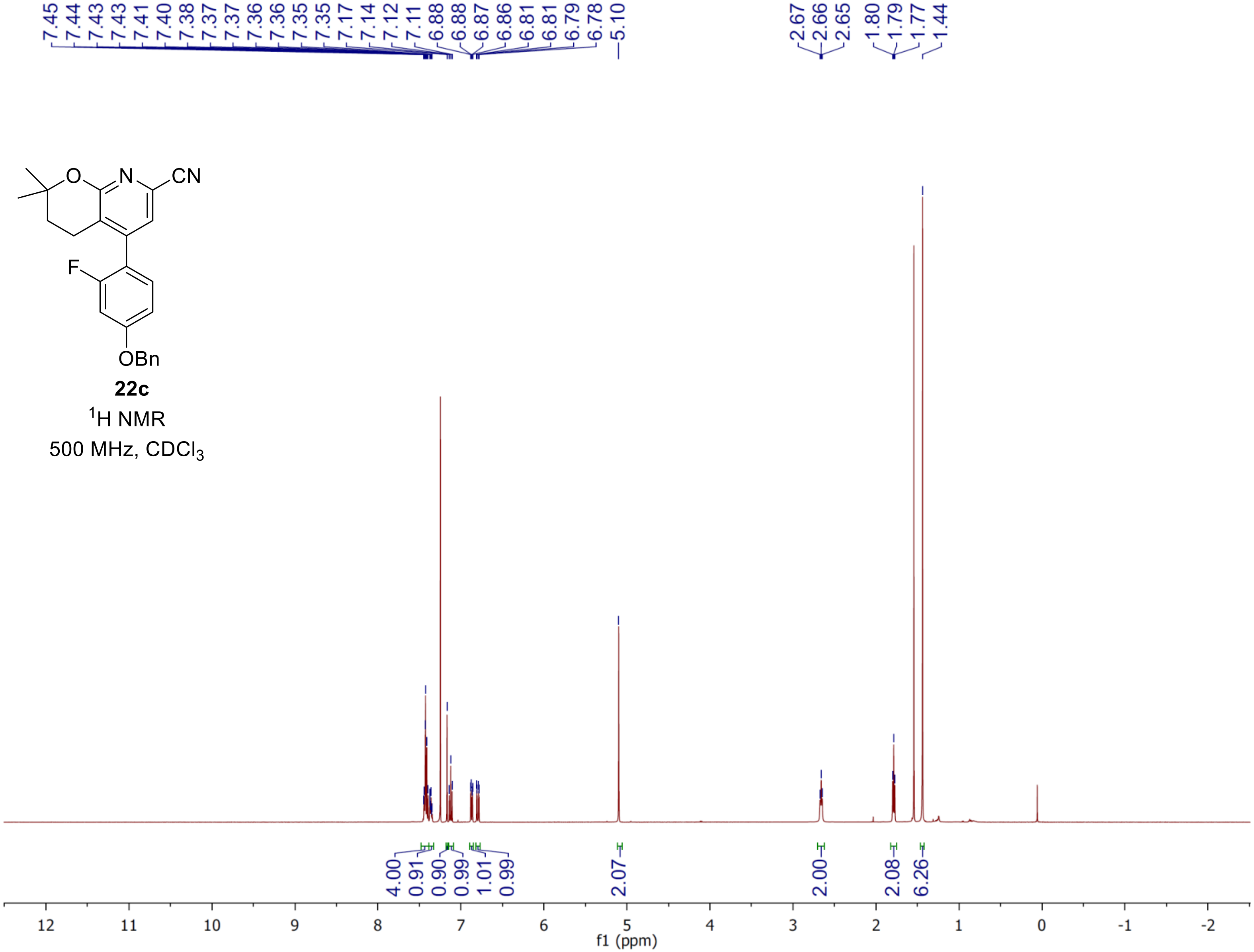


**22c**

$^{1}$H NMR

500 MHz, $CDCl_3$

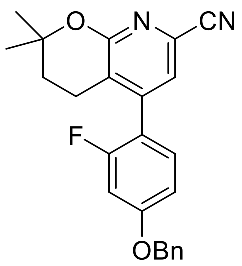

**22c**

$^{13}C$ NMR

125 MHz, $CDCl_3$

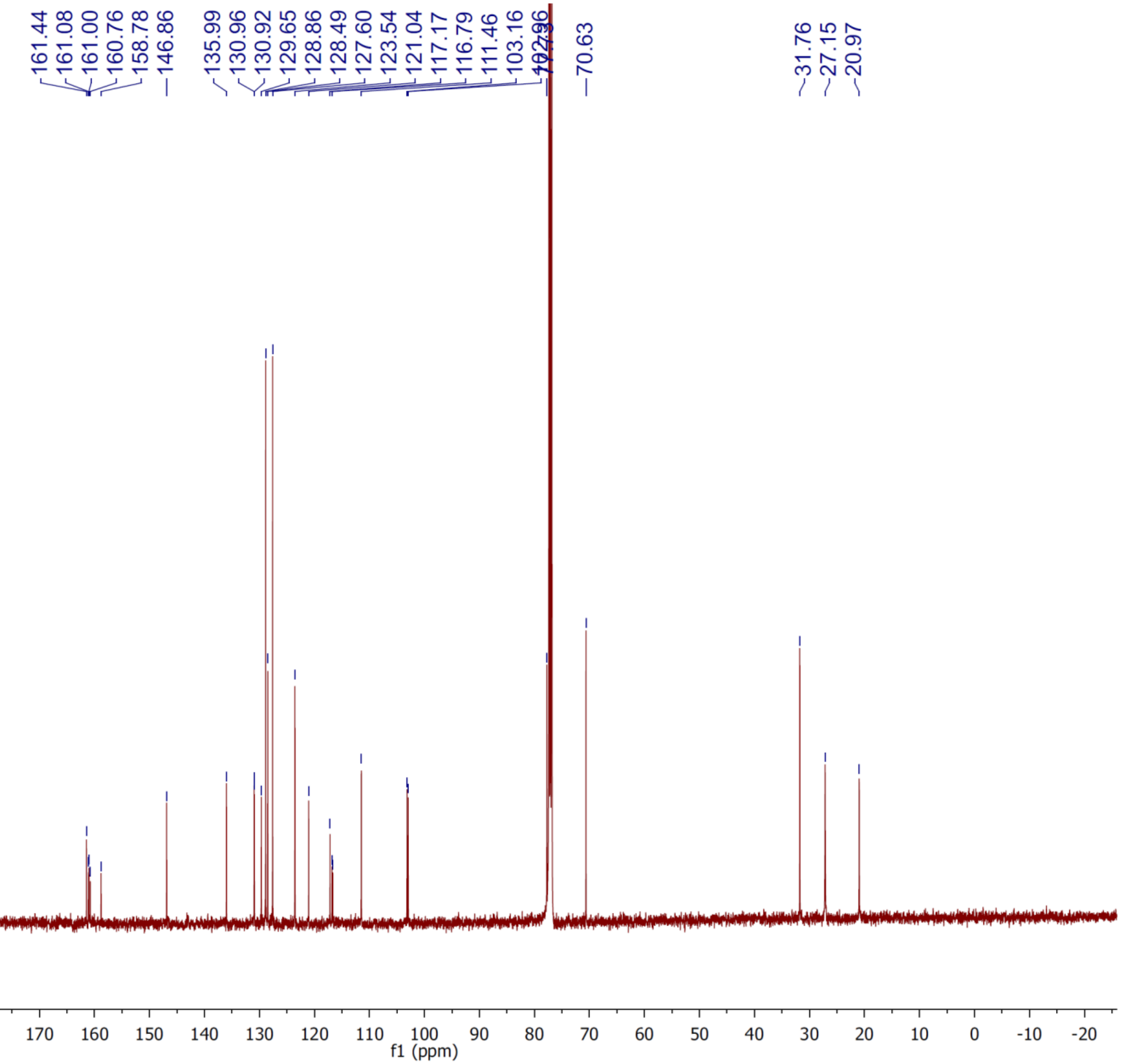

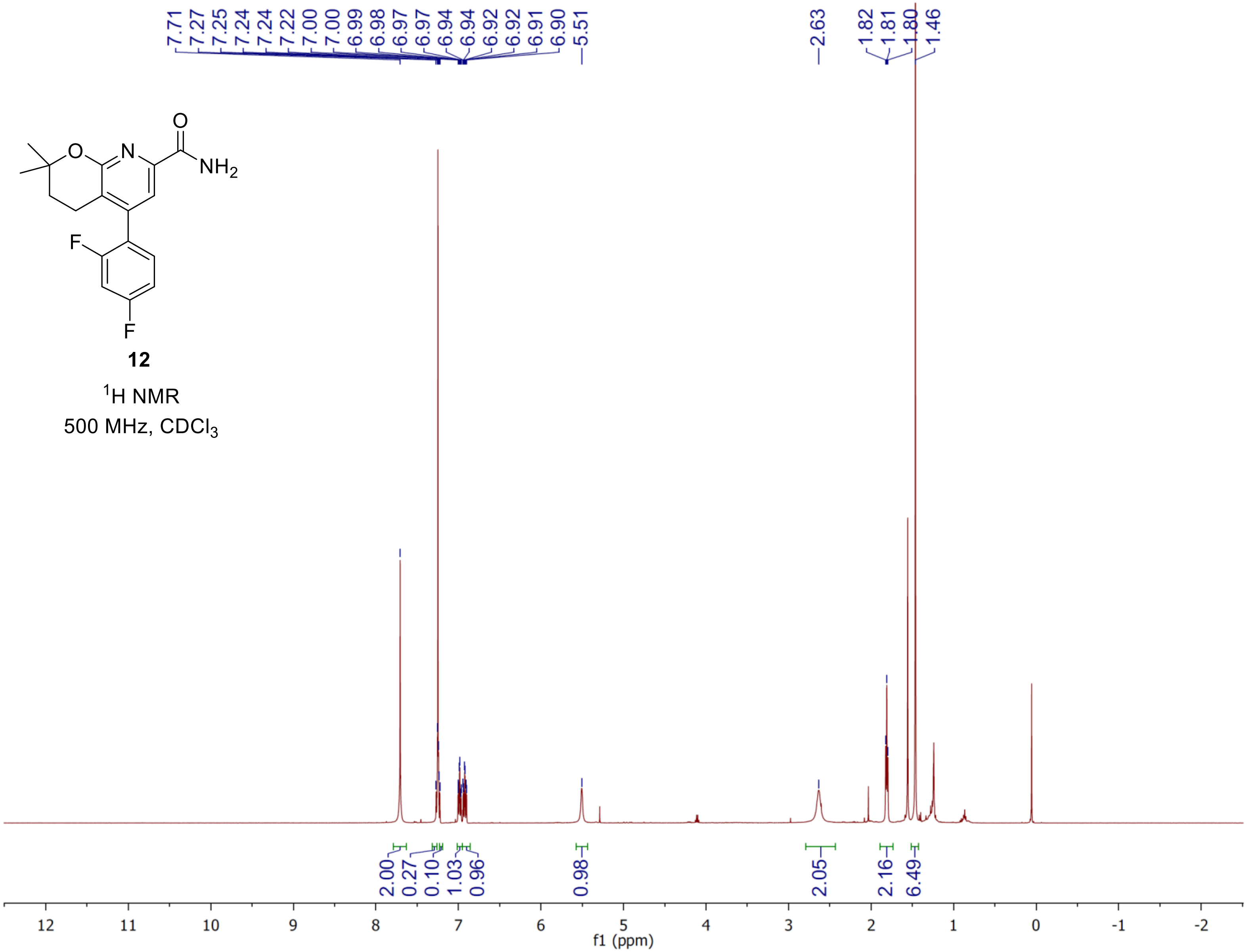

O
O
N
NH2
F
F
12
1H NMR
500 MHz, CDCl3
7.71
7.27
7.25
7.24
7.24
7.22
7.00
7.00
6.99
6.98
6.97
6.97
6.94
6.94
6.92
6.92
6.91
6.90
5.51
2.63
1.82
1.81
1.80
1.46
2.00
0.27
0.10
1.03
0.96
0.98
2.05
2.16
6.49
f1 (ppm)

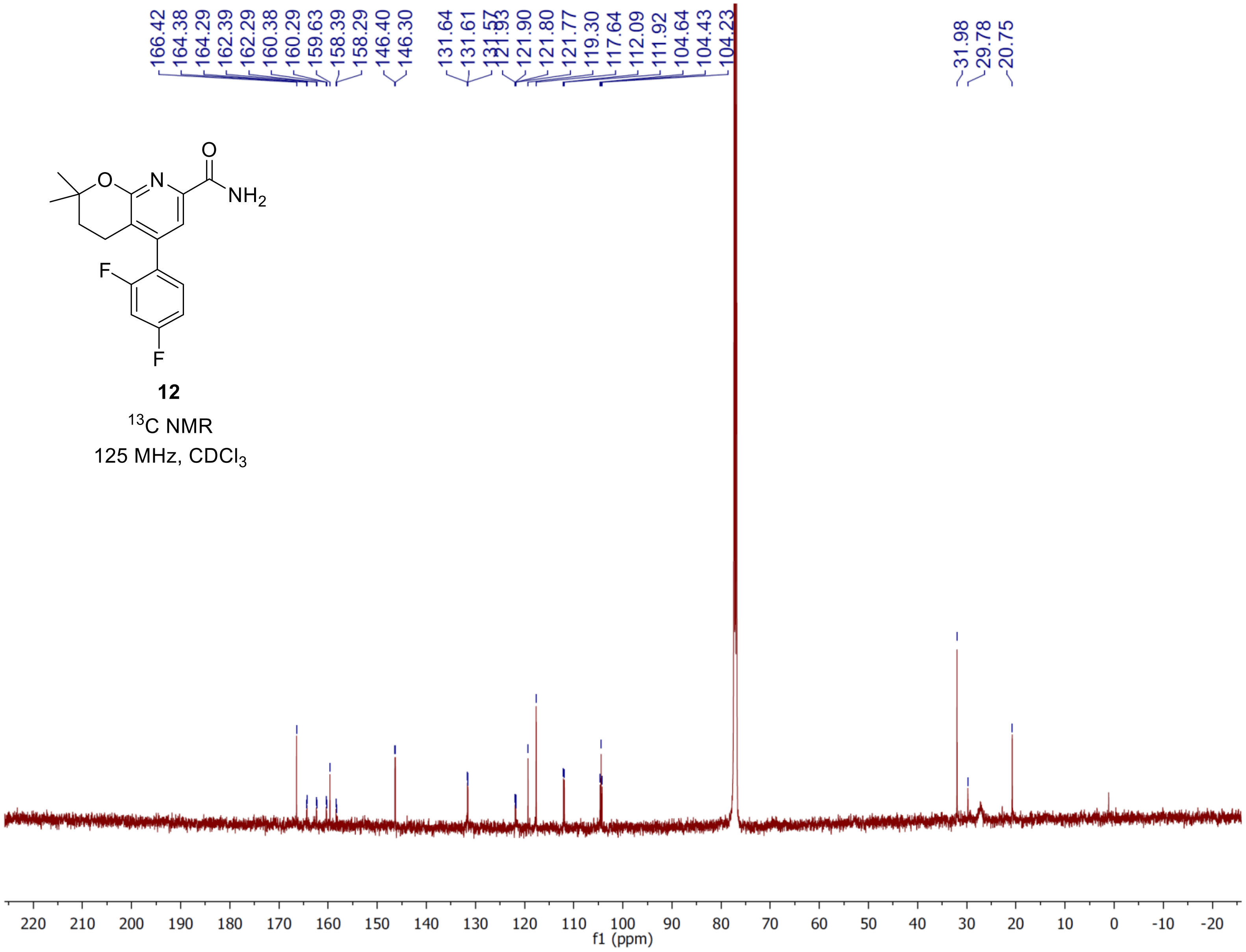
166.42
164.38
164.29
162.39
162.29
160.38
160.29
159.63
158.39
158.29
146.40
146.30
131.64
131.61
131.57
121.93
121.90
121.80
121.77
119.30
117.64
112.09
111.92
104.64
104.43
104.23
31.98
29.78
20.75
O
N
O
NH2
F
F
12
13C NMR
125 MHz, CDCl3
f1 (ppm)

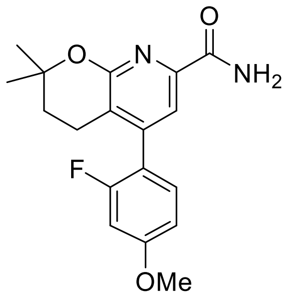


**13**

$^{1}$H NMR

500 MHz, $CDCl_3$

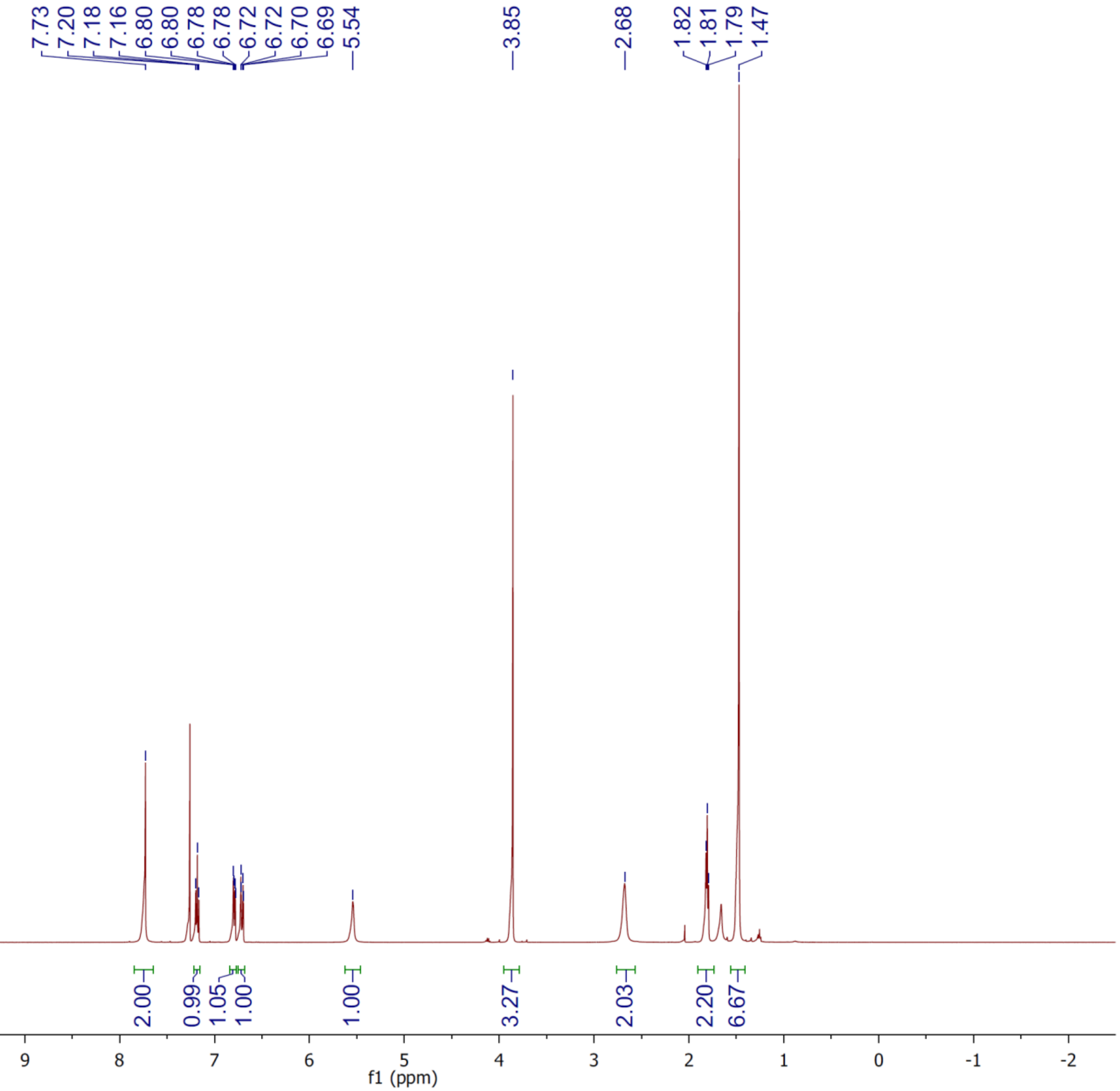

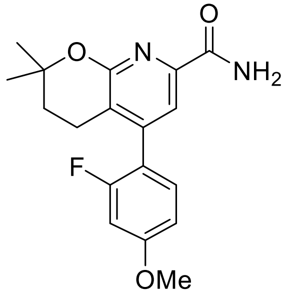

13

$^{13}C$ NMR

125 MHz, $CDCl_3$

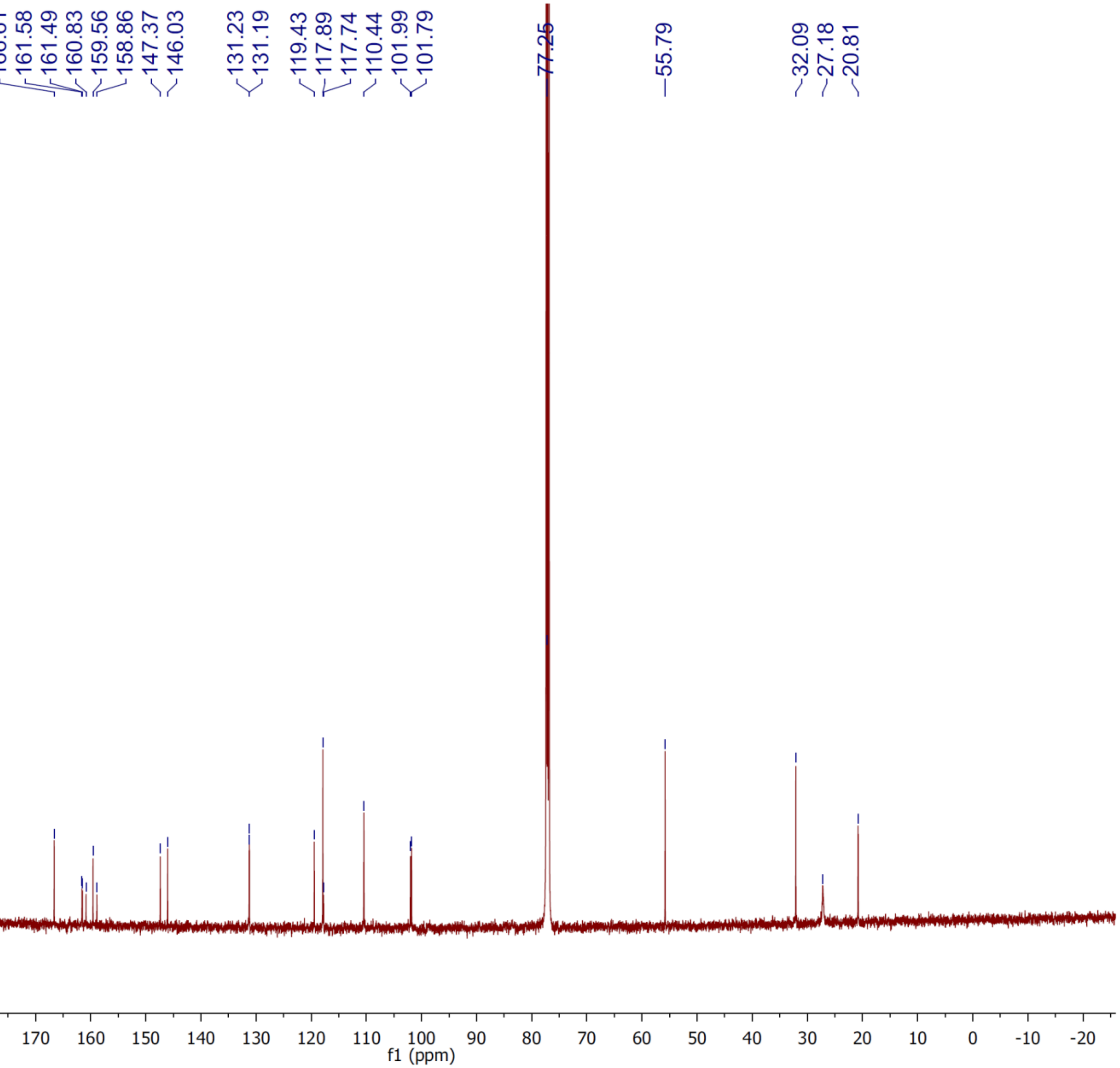

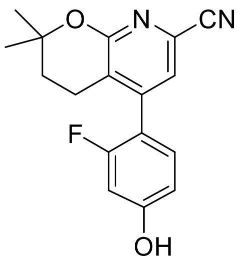

**23**

$^{1}$H NMR

500 MHz, $CDCl_3$

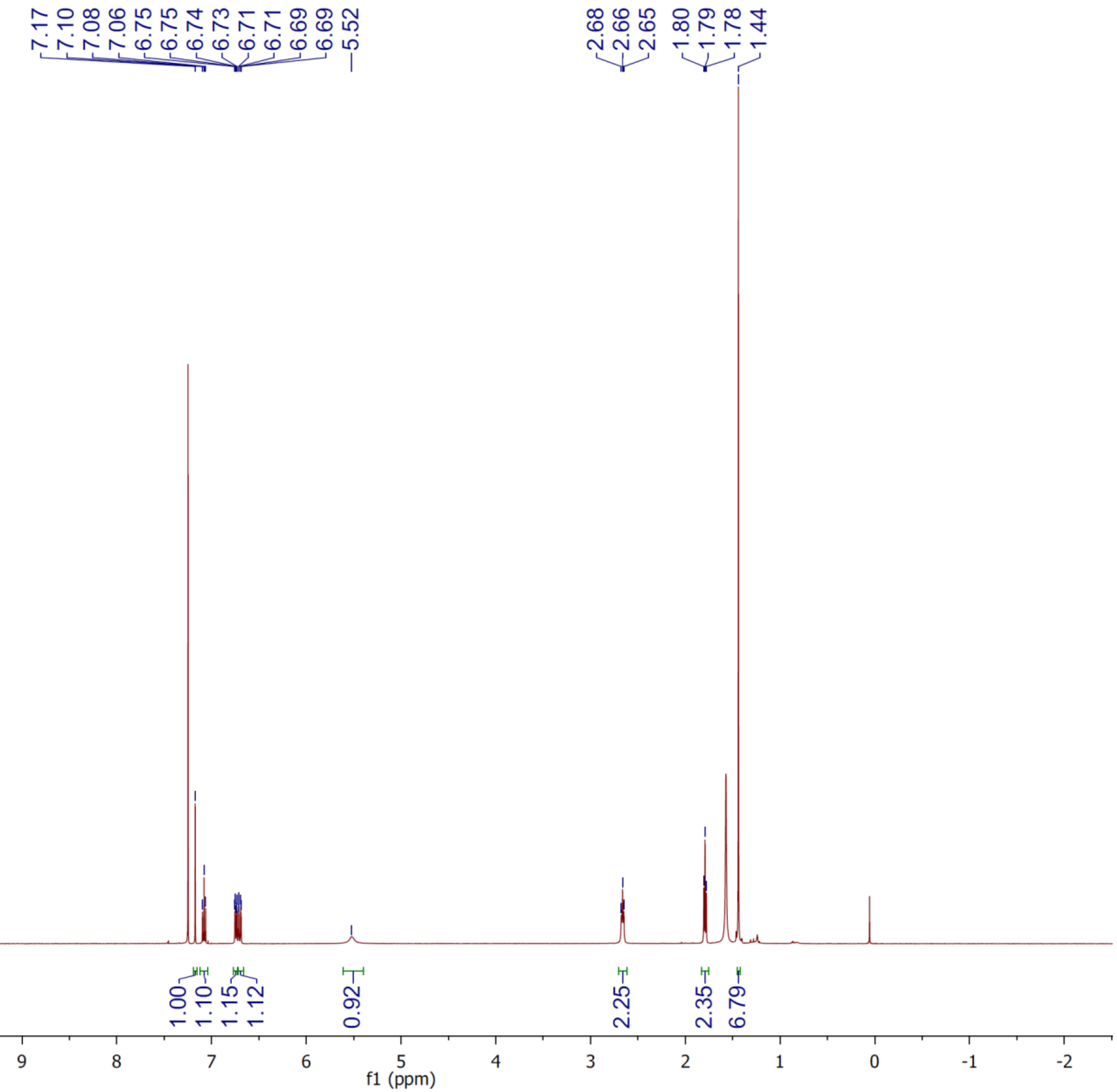

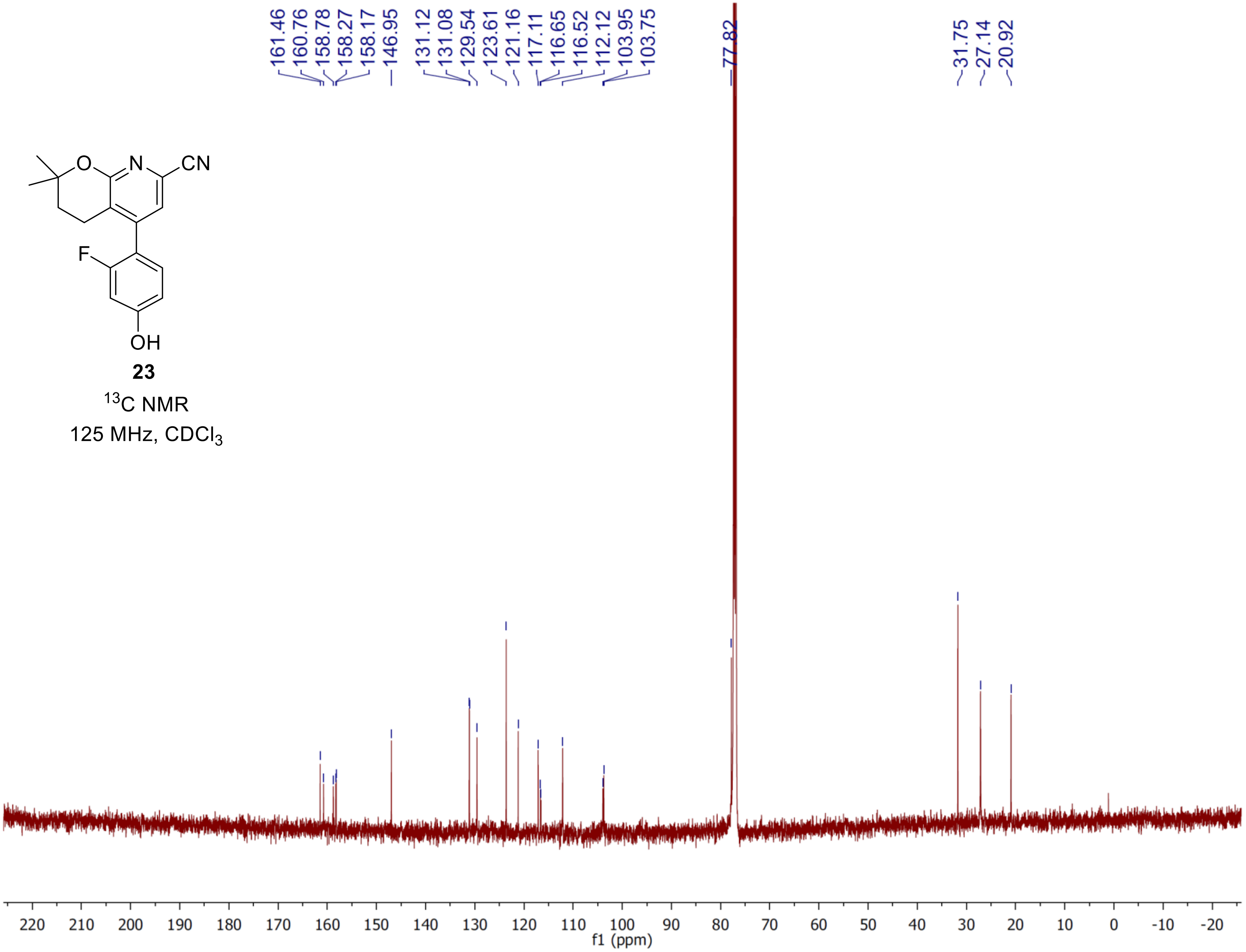

161.46
160.76
158.78
158.27
158.17
146.95
131.12
131.08
129.54
123.61
121.16
117.11
116.65
116.52
112.12
103.95
103.75
77.82
31.75
27.14
20.92
O
N
CN
F
OH
23
$^{13}C$ NMR
125 MHz, $CDCl_3$
220 210 200 190 180 170 160 150 140 130 120 110 100 90 80 70 60 50 40 30 20 10 0 -10 -20
f1 (ppm)

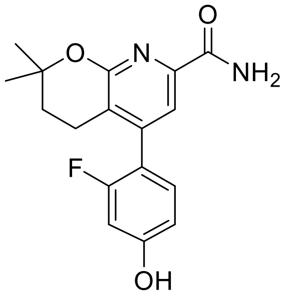


**24**

$^1$H NMR

500 MHz, $CDCl_3$

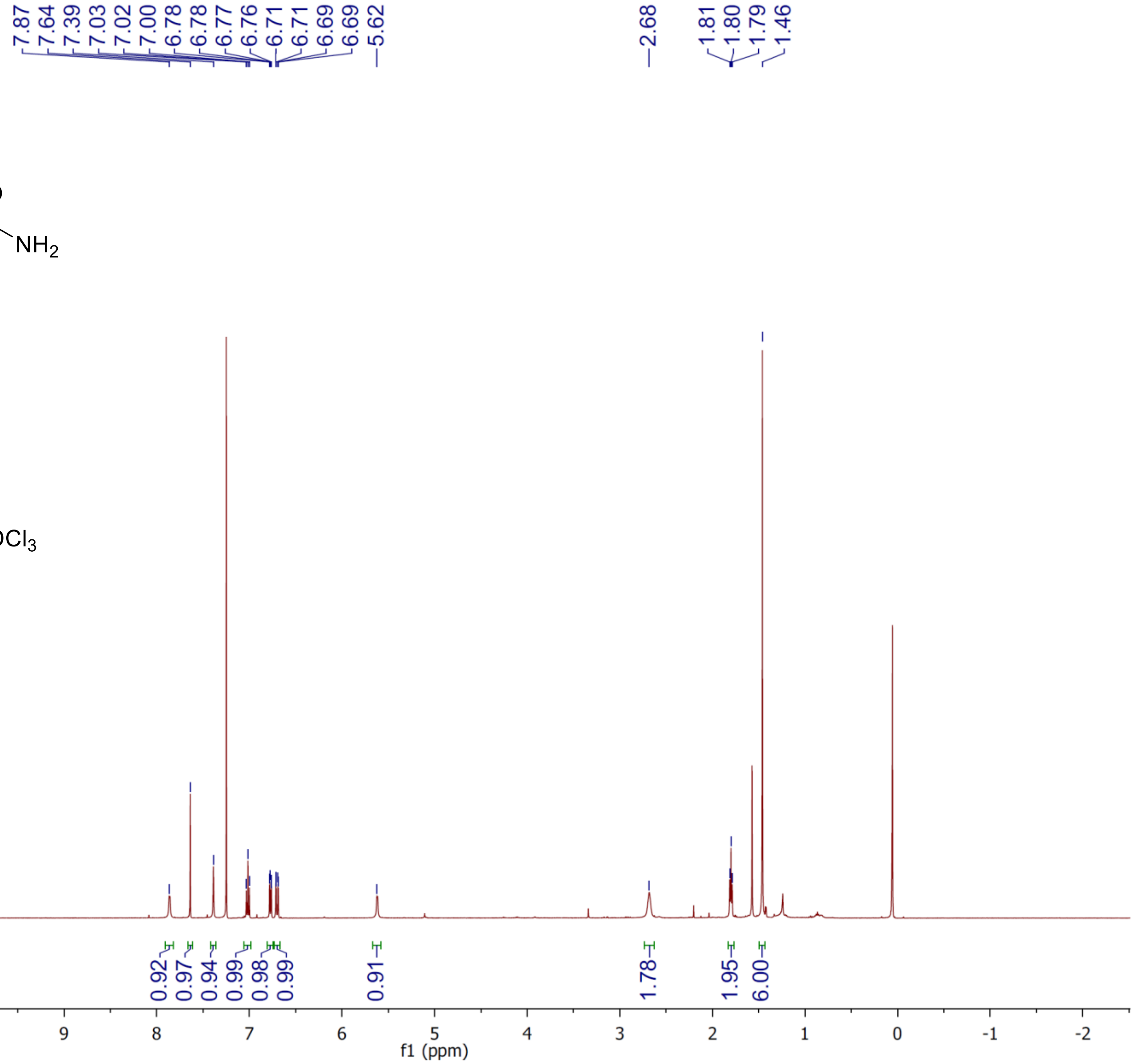

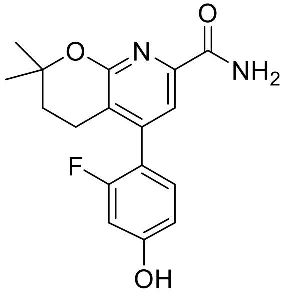

**24**

$^{13}C$ NMR

125 MHz, $CDCl_3$

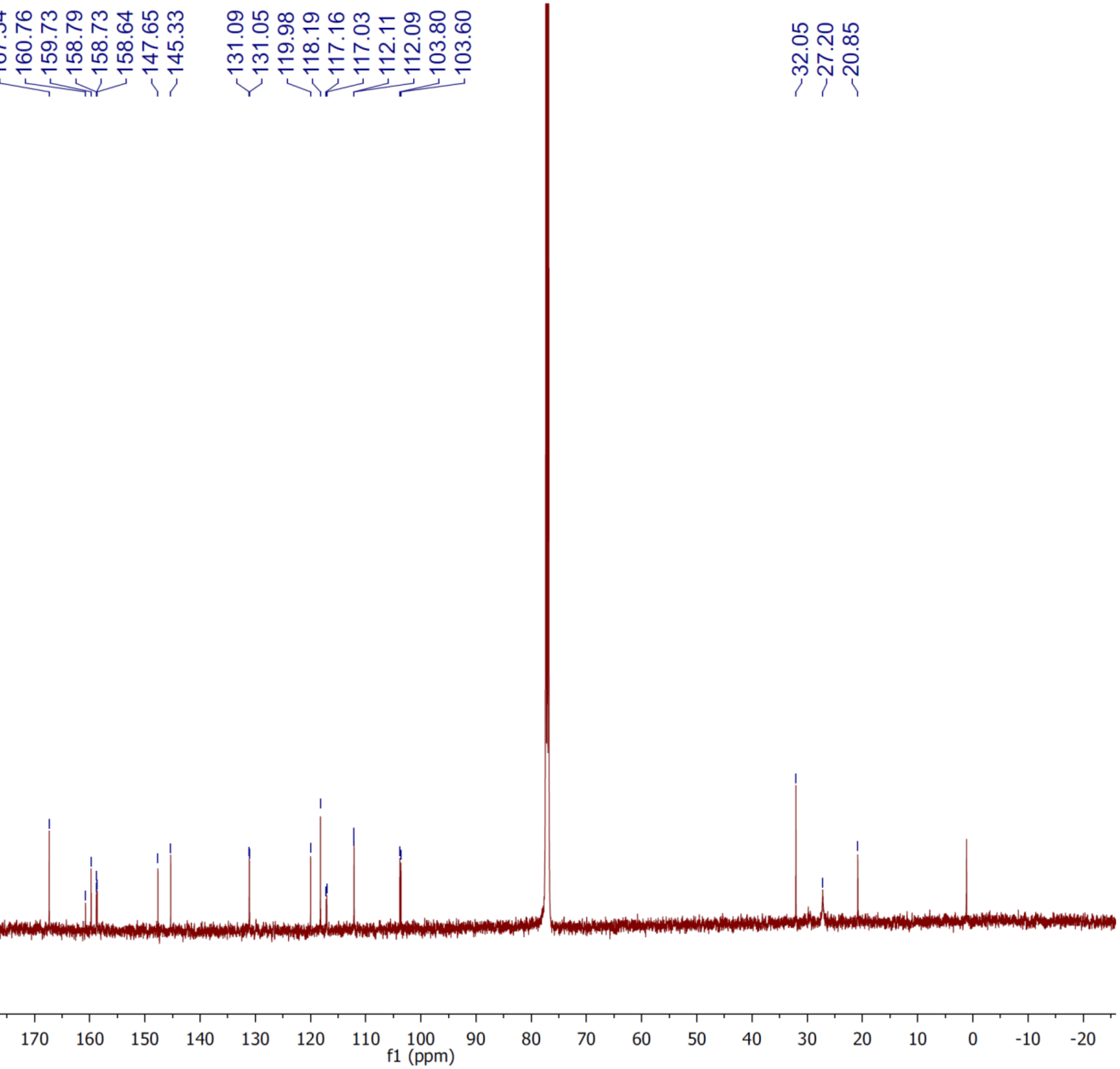
